# Population genetics of polymorphism and divergence in rapidly evolving populations

**DOI:** 10.1101/2021.06.28.450258

**Authors:** Matthew J. Melissa, Benjamin H. Good, Daniel S. Fisher, Michael M. Desai

## Abstract

In rapidly evolving populations, numerous beneficial and deleterious mutations can arise and segregate within a population at the same time. In this regime, evolutionary dynamics cannot be analyzed using traditional population genetic approaches that assume that sites evolve independently. Instead, the dynamics of many loci must be analyzed simultaneously. Recent work has made progress by first analyzing the fitness variation within a population, and then studying how individual lineages interact with this traveling fitness wave. However, these “traveling wave” models have previously been restricted to extreme cases where selection on individual mutations is either much faster or much slower than the typical coalescent timescale *T_c_*. In this work, we show how the traveling wave framework can be extended to intermediate regimes in which the scaled fitness effects of mutations (*T_c_s*) are neither large nor small compared to one. This enables us to describe the dynamics of populations subject to a wide range of fitness effects, and in particular, in cases where it is not immediately clear which mutations are most important in shaping the dynamics and statistics of genetic diversity. We use this approach to derive new expressions for the fixation probabilities and site frequency spectra of mutations as a function of their scaled fitness effects, along with related results for the coalescent timescale *T_c_* and the rate of adaptation or Muller’s ratchet. We find that competition between linked mutations can have a dramatic impact on the proportions of neutral and selected polymorphisms, which is not simply summarized by the scaled selection coefficient *T_c_s*. We conclude by discussing the implications of these results for population genetic inferences.

## I. INTRODUCTION

Models of evolutionary dynamics and population genetics aim to predict how observable properties of the distribution of genotypes within a population are shaped by evolutionary forces such as mutations, natural selection, and genetic drift. We focus here on the simplest possible models of these three core evolutionary forces, which consider a population consisting of *N* individuals, with each individual subject to new mutations at rate *U* per generation, and with the fitness effect of each mutation drawn from some distribution *ρ*(*s*). More complicated models of evolutionary dynamics can incorporate the effects of additional evolutionary forces (e.g., spatial structure and migration, environmental fluctuations in time or space, and ecological interactions). However, the dynamics arising from even the simplest models—incorporating only the forces of mutation, selection, genetic drift—are already rather complex.

Within the context of these simple models, we wish to understand how several types of observables depend on the key parameters. Specifically, we aim to predict the probability, *p*_fix_(*s*), that a mutation with fitness effect *s* will fix. From this quantity, we can compute the rate at which mutations will accumulate (i.e. the rate of genotypic divergence from an ancestor) and the rate, *v*, at which the mean fitness of the population changes over time (McCandlish and Stoltzfus, 2014). In addition to these properties of long-term divergence, we wish to predict expected patterns of genetic diversity within the population at any given time. For example, we aim to compute the coalescence timescale *T_c_* (the typical time since individuals are related by common ancestry, which determines the expected overall level of genetic diversity in the population), as well as other readily observable quantities such as the site frequency spectrum (which describes the relative abundances of polymorphisms at different allele frequencies). Obtaining predictions for patterns of genetic diversity is of particular interest because of the potential to use these predictions, in combination with measured levels of contemporary sequence diversity, to infer the strength of the various evolutionary forces which have acted on a population.

Much classical work proceeds by assuming the different loci in the genome evolve completely independently of one another (i.e., by ignoring the physical *linkage* of mutations within the same genomic segment). The dynamics are then relatively straightforward to analyze (Fisher, 1931; Wright, 1931). This assumption is reasonable in sexually evolving populations where recombination is sufficiently rapid compared to the timescales on which mutation, selection and genetic drift generate correlations between loci (Falconer and Mackay, 1996). In this case, a given mutation is shuffled onto a large number of genetic backgrounds over the course of its evolutionary trajectory, and any correlations in the fates of mutations at different loci become negligible.

In asexual populations, however, mutations are are fully linked and do not evolve independently of one another. Even in the genomes of obligately sexual organisms, loci that are physically close within a chromosome will be broken up by recombination only over very long timescales, and cannot necessarily be treated as independently evolving (Weissman and Barton, 2012). A useful opposite regime to consider is then the limit of a genomic region that is small enough that recombination can be entirely neglected. One can imagine that on sufficiently short genomic distance scales (within so-called “linkage blocks”), evolution proceeds entirely asexually, while on larger genomic distance scales evolution acts on some collection of recombining blocks. It is not entirely clear how this division between asexual linkage blocks and larger-scale regions consisting of multiple recombining blocks is best modeled, though some recent work has begun to address this question (Good et al., 2014; Neher et al., 2013; Weissman and Hallatschek, 2014). However, we focus in this paper on an even simpler question: how to model the evolution within a single nonrecombining linkage block. As we will see, even within the context of the simple models described above, evolutionary dynamics even within perfectly asexual linkage blocks are surprisingly complex, and remain incompletely understood.

There is a long history of efforts to analyze the dynamics of purely asexual evolution, and numerous approaches have been proposed to analyze the dynamics in different limiting regimes of the parameter space. Extensive work has focused on the *neutral limit*, in which natural selection can be neglected entirely (Kimura, 1968). In this case, genealogical approaches such as coalescent theory can provide a complete description of the expected rates of divergence and patterns of diversity (Kingman, 1982; Wakeley, 2005). These methods can then be used as the basis for methods to include recombination (e.g., using the ancestral recombination graph (Griffiths, 1997), or using variations of the sequentially Markovian coalescent (McVean and Cardin, 2005)), and can also incorporate other complications such as geographic subdivision, population structure, and demographic changes (Nordborg, 2004). However, the backwards-time nature of the coalescent approach makes it difficult to incorporate the effects of selection: genealogies cannot be considered independently of the selected mutations which occur on their branches (Kaplan et al., 1988). Efforts to do so have largely been limited to simulation-based or essentially numerical approaches, such as the ancestral selection graph (Krone and Neuhauser, 1997), though analytical progress has been made in certain cases, particularly in the presence of purifying selection on deleterious mutations (Charlesworth, 1994; Hudson and Kaplan, 1995).

To model the effects of selection on beneficial mutations, much work has instead been done using forward-time approaches. Broadly speaking, these approaches seek to characterize the trajectories of mutant lineages in a probabilistic sense. Provided selection is sufficiently strong and selected mutations are sufficiently rare (more precisely, if *Ns* ≫ 1 and *NU* log *Ns* < 1), a beneficial mutation typically either sweeps to fixation or is purged before another such mutation arises within the same linkage block (Desai and Fisher, 2007). Thus, within this *strong-selection weak-mutation* (SSWM) regime, multiple selected mutations are unlikely to segregate simultaneously within the block. In this case, the dynamics of each selected mutation can be treated independently of one another, using the single-locus methods described above (Gillespie, 1983). The impact of these mutations on linked neutral diversity can also be calculated from these dynamics (e.g., to analyze the signature of an isolated selective sweep) (Smith and Haigh, 1974). However, recent work has shown that in a wide range of microbial and viral populations, and potentially in many linked regions of the genomes of obligately sexual organisms such as humans, multiple beneficial mutations often segregate simultaneously (Lieberman et al., 2014; Miralles et al., 1999; Nourmohammad et al., 2019; Strelkowa and Lässig, 2012). In these *rapidly adapting* populations, clonal interference (i.e. competition between multiple distinct adaptive lineages) and genetic hitchhiking can be critical to the dynamics, and analyzing the evolution of multiple linked loci simultaneously is critical (Buskirk et al., 2017; Gerrish and Lenski, 1998; Kao and Sherlock, 2008; Lang et al., 2013).

Over the past two decades, numerous authors have analyzed the evolutionary dynamics of many linked loci in rapidly adapting populations (reviewed by Neher (2013)). Similar ideas have also been used to analyze the dynamics of populations rapidly *declining* in fitness due to Muller’s ratchet (Neher and Shraiman, 2012), as well as those maintained in a dynamic steady-state balance between beneficial and deleterious mutations (Goyal et al., 2012). Collectively, we can think of this body of work as studying the evolutionary dynamics of rapidly *evolving* populations—those populations in which there are typically multiple linked mutations, either beneficial or deleterious, in the population at once. The central idea of this work is to first analyze how the collective effects of many linked mutations generate variation in fitness within the population. This leads to a time-dependent within-population fitness distribution that can be described as a *traveling wave*, and which maintains a constant steady-state shape *f*(*x*) while its mean fitness changes at a constant rate *v* (potentially with fluctuations) (Rouzine et al., 2008, 2003; Tsimring et al., 1996). Extensive work has characterized how the velocity and steady-state shape of the wave depends on the key parameters, which can be used to compute observables related to long-term divergence such as *p*_fix_(*s*) or the distribution *ρ_f_*(*s*) of *fixed* fitness effects (Good et al., 2012). More recent studies have then used this traveling wave picture as the basis for tracing the genealogical history of individual mutations (Desai et al., 2013; Neher and Hallatschek, 2013) or tracking their trajectories forward in time (Kosheleva and Desai, 2013) to calculate diversity statistics such as the coalescence timescale or the site frequency spectrum. We provide a schematic of the steady-state traveling wave shape *f*(*x*), as well as an example *ρ*(*s*) and corresponding *ρ_f_*(*s*) in Fig. 1.

**FIG. 1.**
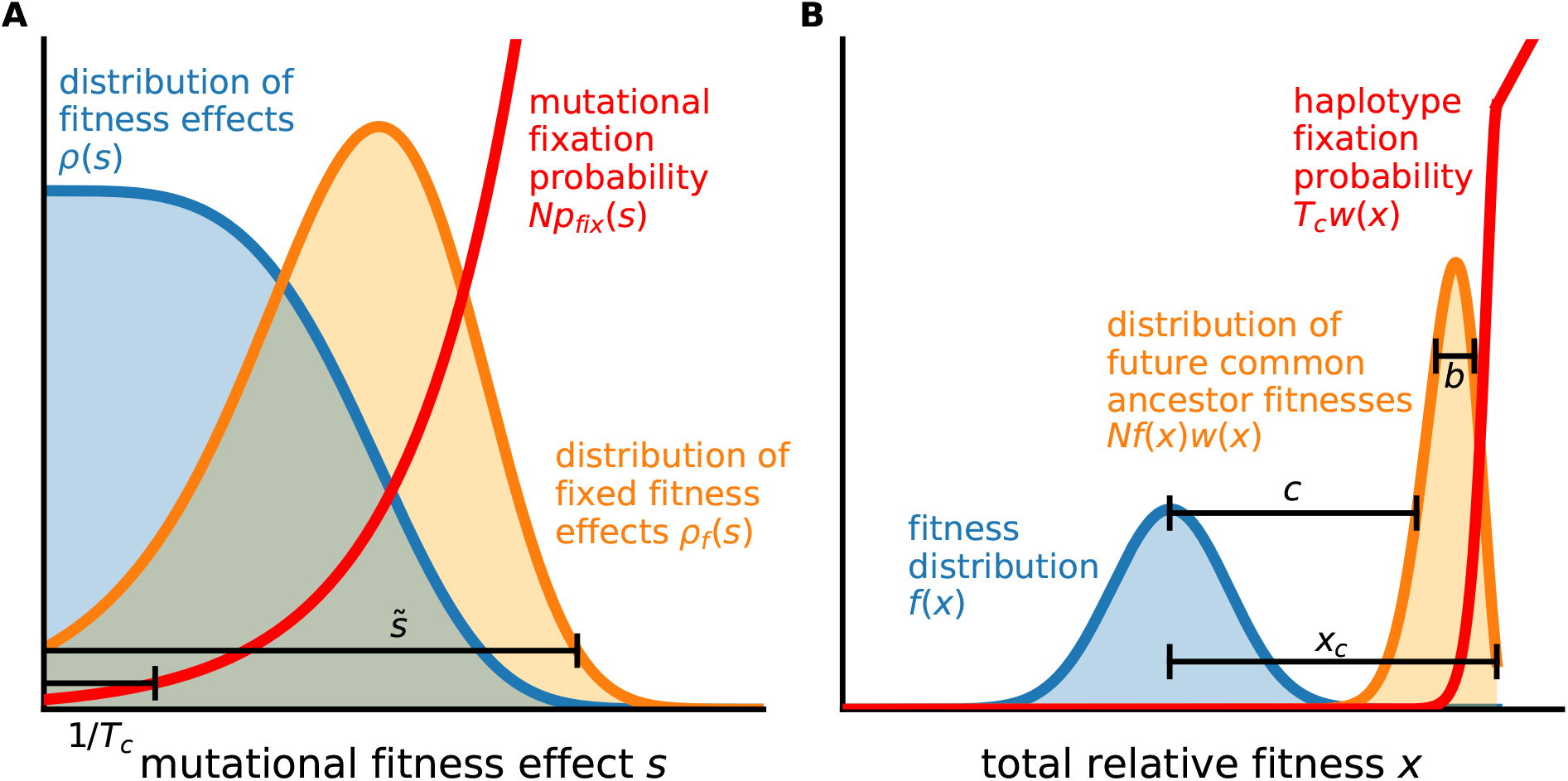
(A) Example distribution *ρ*(*s*) of fitness effects, distribution *ρ_f_*(*s*) of fixed fitness effects, and mutational fixation probability function *p*_fix_(*s*), scaled by the population size *N*. The scale 1/*T_c_*, to be defined precisely in Section IV, will be seen to correspond with the scale on which *Np*_fix_(*s*) varies; selection is strong on individual mutations when 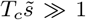. (B) Example distribution *f*(*x*) of individual (haplotype) relative fitnesses, distribution *Nf*(*x*)*w*(*x*) of fitnesses of future common ancestors, and haplotype fixation probability function *w*(*x*), scaled by *T_c_*. The “nose” of the fitness distribution (its high-fitness edge) lies at a fitness advantage *x_c_* compared to the mean; the haplotype fixation probability function *w*(*x*) has a shoulder at the same fitness advantage, with *w*(*x*) ≈ *x* for *x* > *x_c_*. The scales *b* and *c*, to be introduced in Section IV, will be seen to correspond roughly to the width and average, respectively, of the distribution *Nf*(*x*)*w*(*x*).

While this work has led to substantial progress, it has been focused primarily on two limiting cases. One body of work (Cohen et al., 2005; Hallatschek, 2011; Neher and Hallatschek, 2013; Tsimring et al., 1996) has analyzed the *infinitesimal limit*, in which the the selective effects of individual mutations are infinitesimal, but selected mutations occur extremely frequently (more precisely, in which *s* → 0 and *U* → ∞ with the product *U* 〈*s*^2^〉 held fixed). The corresponding *infinitesimal approximation* is often thought to be valid when *s* ≪ *U*, though a more accurate condition of its validity is that *T_c_s* ≪ 1 (Good and Desai, 2014; Good et al., 2014); in both conditions, the relevant *s* can be taken as the rootmean-squared effect 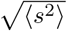. As a result, within the infinitesimal regime, selection on individual mutations can be neglected: the timescale 1/*s* on which selection can substantially alter the fate of a mutation with effect size *s* is longer than the coalescent timescale *T_c_* on which common ancestry is determined (and which we define more precisely in Section IV). At the same time, the population as a whole maintains substantial variance in fitness, *σ*^2^, resembling in some ways infinitesimal models in quantitative genetics (Barton et al., 2016); selection on *haplotypes* in the infinitesimal regime can be strong. This work has led to analytical results for both divergence-related quantities and, more recently, diversity-related quantities (Neher and Hallatschek, 2013), valid for populations subject to beneficial mutations, to deleterious mutations, or to some combination of the two.

A second body of work has analyzed the opposite limit of *strong* selection on individual mutations, such that 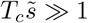; here 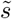 denotes the typical effect size of mutations which fix (with 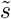 defined more precisely in Section IV). When selection is strong on deleterious mutations, Muller’s ratchet clicks slowly and beneficial mutations can much more easily dominate the dynamics. While some work has been done on the rate of Muller’s ratchet when 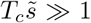 (Neher and Shraiman, 2012), the majority of this work has focused exclusively on beneficial mutations (Desai and Fisher, 2007; Desai et al., 2013; Fisher, 2013; Good et al., 2012; Kosheleva and Desai, 2013) or on the case in which deleterious mutations can be considered a perturbative correction (Good and Desai, 2014). This work can be further divided into two regimes: the “moderate-speed” regime, in which 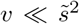, as well as the “high-speed” regime, in which 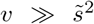 (Fisher, 2013). Qualitatively, the “moderate-speeds” and “high-speeds” regimes can be distinguished based on the range of background fitnesses Δ*x_f_* which typically produces an eventual common ancestor of the population: in the “moderate-speeds” regime, 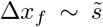, while in the “high-speeds” regimes, 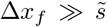. That is, within the “moderate-speeds” regime, the individual which will eventually fix is likely within one mutational effect 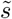 of the “nose” (the high-fitness edge) of the fitness distribution; in the “high-speeds” regime, individuals can catch up and fix from further behind by rapidly acquiring multiple mutations. Analytical results for divergence-related quantities have been obtained within both the “moderate-speeds” and “high-speeds” regimes (Fisher, 2013; Good and Desai, 2014); diversity-related quantities have also been studied within the “moderate-speeds” regime (Desai et al., 2013; Kosheleva and Desai, 2013).

Although the parameters *N*, *U* and *s* are natural quantities to use in specifying a model of the evolutionary dynamics, in many applied settings the combined quantities *T_c_s* and *T_c_U* can be probed more directly. While dynamical interpretations of *T_c_s* and *T_c_U* are less immediately clear, these can be considered independent properties of the evolution that together can describe many aspects of a population. Existing methods to infer *T_c_s* from observed levels of polymorphism and divergence among populations typically make the assumption that different loci evolve *independently* of one another (Bustamante et al., 2001; Sawyer and Hartl, 1992). For example, a classic result shows that in a population with a constant size *N*, the ratio between selected and neutral site frequency spectra is given by

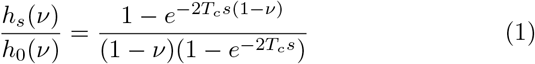

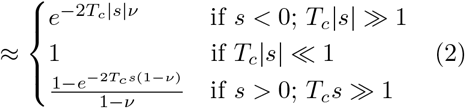

where *T_c_* = *N* is the coalescent timescale (Sawyer and Hartl, 1992). Using this result, the magnitudes of the scaled selection coefficients (*T_c_s*) can be identified from the transitions in the site frequency spectra at low frequencies (*ν_c_* ~ 1/*T_c_*|*s*|) for deleterious mutations, or at high frequencies (1 – *ν_c_* ~ 1/*T_c_s*) for beneficial mutations.

However, previous work has shown that these classical results can lead to substantial errors in the inferred selection strengths when natural selection is widespread (Messer and Petrov, 2013). This makes it difficult to identify the relevant values of *T_c_s* in rapidly evolving populations. For deleterious mutations, the intermediate regime where *T_c_s* ~ 1 is particularly critical: it is precisely mutations with effect sizes on the order of 1/*T_c_* which are expected to have the largest impact on patterns of genetic diversity within a population (Good et al., 2014). Thus when there is a relatively broad distribution of effect sizes of new mutations (including strong, intermediate, and weak-effect mutations), it may often be the case that the intermediate-effect mutations with *T_c_s* ~ 1 have the largest impact on observed patterns of diversity. More generally, an understanding of the intermediate regime in which *T_c_s* ~ 1 is critical for understanding the transition between strong selection and neutral-like behavior in site frequency spectra. Our understanding of these effects is limited, however: except for special cases—such as when a single strong beneficial fitness effect is available (Kosheleva and Desai, 2013)—we lack analytical predictions for the diversity statistics of selected mutations in rapidly evolving populations. In particular, we lack predictions for how mutations with a distribution of beneficial and/or deleterious selective effects contribute to the site frequency spectrum, which has in turn limited our ability to apply PRF-based inference methods (e.g. of Sawyer and Hartl (1992)) to rapidly evolving populations.

To address this gap, in this work we reexamine the approach used by Fisher (2013) to study the “high-speeds” regime. We demonstrate that the key ideas of that approach can be applied more generally to the case in which 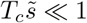 (as well as the intermediate regime in which 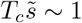. As a result, we argue that the infinitesimal regime and the “high-speeds” regime—previously studied separately—can be unified into a single *moderate selection, strong mutation* (MSSM) regime, which includes populations subject to a distribution of beneficial mutations, deleterious mutations, or some combination of the two. The key requirements for validity of our MSSM approximation are that 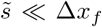 (which ensures that selection is moderate, or weak, on single mutations) and that *T_c_*Δ*x_f_* ≫ 1 (which ensures that mutation is strong, and that selection on haplotypes is strong). Dynamically, these conditions imply that mutational “leapfrogging” is important to the dynamics: *some* individuals routinely catch up to the high-fitness “nose” despite an initial fitness disadvantage, relative to the nose, of several mutations. These requirements are satisfied in the limit *N* → ∞ with *Uρ*(*s*) held fixed—as long as *ρ*(*s*) falls off faster than exponentially with large positive *s*—as well as in other cases we describe. In particular, we show that this approach can model the dynamics of deleterious mutations with 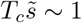, to which patterns of diversity and divergence are particularly sensitive.

Using this approach, we compute divergence-related statistics such as the fixation probabilities *p*_fix_(*s*) of new mutations and the average rate *v* of adaptation, or—if *v* < 0—of Muller’s ratchet. We also compute diversity-related statistics such as the coalescence timescale *T_c_* and the site frequency spectrum. We show that at high frequencies, the neutral site frequency spectrum corresponds to that of the Bolthausen-Sznitman coalescent (BSC) (Bolthausen and Sznitman, 1998), which has previously been found to describe genealogies of populations in the infinitesimal regime (Neher and Hallatschek, 2013) and “moderate-speeds” regime (Desai et al., 2013), as well as in evolving populations modeled as FKPP waves (Brunet et al., 2007). We identify the frequency scale above which this correspondence holds and analytically describe departures from the BSC at lower frequencies. Importantly, we find that the low-frequency portion of the neutral site frequency spectrum is much more useful for distinguishing different parameter combinations than the high-frequency portion, which depends on the parameters *N*, *U* and *ρ*(*s*) via a single overall scale factor. Over a broad range of intermediate and high frequencies—extending beyond the frequency range at which a correspondence with the BSC exists—we demonstrate that the neutral and selected site frequency spectra are simply proportional to one another, with a constant of proportionality equal to the ratio of fixation rates of the two types of mutations. This proportionality reflects the fact that, in the presence of widespread linkage, the fates of even strongly selected mutations can be considered *conditionally neutral* at long times *t* > *T_c_*: at this point, their initial fitness effects have been “forgotten”, and their lineage trajectories are indistinguishable from those of neutral mutations with the same age (and frequency at age *T_c_*). We discuss implications of our results for inferring the strength and/or frequency of selection in natural populations using observed levels of polymorphism and divergence.

### Outline of this Paper

The remainder of this work is organized as follows. In Section II, we describe the model of the population dynamics and briefly summarize our simulation methods. In Section III, we review previous approaches to model evolutionary dynamics using traveling wave theory. We begin Section IV by presenting our MSSM approximation, which we use to obtain results for the steady-state distribution of fitnesses, fixation probabilities, and the rate of adaptation. We then provide and discuss conditions of validity of the MSSM approximation, apply the MSSM approximation to several specific examples of DFEs, and compare our analytical results to the results of simulations. In Section V, we use our MSSM approximation to analyze statistics of genetic diversity, with particular focus on the site frequency spectra of neutral mutations and of selected (non-neutral) mutations. In Section VI, we provide heuristic interpretations of several fitness scales and timescales that emerge in our analysis, and compare dynamical aspects of populations described by the MSSM approximation to those of populations in other regimes. In the Discussion, we consider implications of our results for making inferences from population genetic data.

## II. MODEL

We aim to understand the evolutionary dynamics of an asexual population, given the simplest possible model incorporating the effects of mutations, natural selection, and genetic drift. To this end, we consider a population of a constant number *N* asexual haploid individuals, within which new mutations arise at a rate *U* per genome per generation. We assume that all mutations have additive effects on (log) fitness, *X* (i.e., that the log fitness of each individual is the sum of the fitness effects of all the mutations it or its ancestors have acquired). An individual’s offspring number distribution is determined by its relative fitness 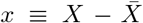, where by 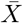 we denote the population-wide mean fitness. We write down the particular stochastic birth/death process we carry out in simulations in Subsection II.A; in our analytical treatment, we assume a slightly different birth/death process, which we write down in Section III. We assume the fitness effect of each new mutation is drawn at random from a time-independent distribution of fitness effects (DFE), *ρ*(*s*). Our model thus assumes that while *epistasis* may exist among individual mutations, there are no overall differences in the DFE among different genotypes, so we can treat *ρ*(*s*) as a constant that remains the same as the population evolves. Because most dependence on the parameters *U* and *ρ*(*s*) will be mediated by the product *μ*(*s*), we define *μ*(*s*) ≡ *Uρ*(*s*) as the *mutational fitness spectrum*. We assume neutral mutations occur at rate *U_n_* (not included in *U*).

The actual DFEs relevant to natural populations may be broad and complex, with mutations conferring a wide range of fitness effects that occur at a variety of different rates, and these empirical distributions are difficult to measure precisely (Eyre-Walker and Keightley, 2007). Much of our analysis is conducted for an arbitrary DFE (provided it meets our conditions of validity). For concreteness, we also focus on a few simplified forms of *μ*(*s*) in order to gain intuition. For example, we consider the cases in which (i) all mutations are beneficial, (ii) all mutations are deleterious, and (iii) all mutations confer the same effect size. The analysis of these simplified forms of *μ*(*s*) can provide useful intuition about how mutations of different types and effect sizes affects various aspects of the evolutionary dynamics. Further, we find that most of our results are sensitive to the assumed *μ*(*s*) only within a limited region of *s*. That is, given values of the other parameters, the evolutionary dynamics are often dominated by a subset of mutations that have a narrow range of effect sizes, and hence can be predicted based on simplified forms of *μ*(*s*) (Hegreness et al., 2006).

### A. Simulation methods

To test our analytical predictions, we conduct individual-based Wright-Fisher simulations. These simulations, which we describe in more detail in Appendix H, separately track all of the individuals in the population, as well as the mutations they have acquired. These simulations make it possible to measure divergence-related and diversity-related quantities in the population, including the rate of change in the mean fitness, the rate of mutation accumulation, the heterozygosity of neutral mutations and of selected (i.e. non-neutral) mutations, and the site frequency spectrum.

Simulations consist of a series of two steps repeated each generation. First, each individual acquires a Poisson-distributed number of mutations (with mean number *U*, and with the fitness effects of those mutations each drawn from *ρ*(*s*)). Mutations increment an individual’s log-fitness *X* by an amount *s*. Second, in the selection/reproduction step, individuals in the population are resampled (with replacement) to form the population in the following generation. Each individual is sampled with with probability proportional to its (exponential) fitness *e^X^*.

Each simulated population is initialized clonally. We record the number of generations until a mutation has fixed within the population, which we define as the length of an *epoch*. At the conclusion of each subsequent epoch, we record the state of the population, including its mean fitness, site frequency spectrum, and fixed mutations. Simulations are run for a prespecified number of epochs, with results from the first 10 epochs discarded.

Simulated populations are subject to stretched exponential DFEs of the form

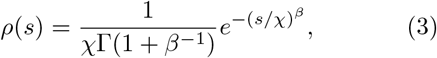

or to mutations consisting of a single fitness effect (while our analysis applies for more general *ρ*(*s*)). DFEs of the form in Eq. (3) have been considered in previous theoretical work (Desai and Fisher, 2007; Fogle et al., 2008; Good et al., 2012). Note that if *β* > 1, *ρ*(*s*) in Eq. (3) falls off faster than exponentially with large *s*, while *ρ*(*s*) is an exponential distribution if *β* = 1. For *β* < 1, *ρ*(*s*) in Eq. (3) falls off more slowly than exponentially with large *s* (though for reasons we describe below, we do not consider this case extensively here).

## III. THE TRAVELING-WAVE APPROACH

In this Section, we review the basic traveling-wave approach which underlies our work, and which has been used to study the dynamics of rapidly evolving populations in multiple regimes of the parameter space. Readers familiar with the traveling-wave literature may wish to skip directly to the Analysis section. We emphasize that the details of the traveling-wave approach have differed across studies—particularly in the treatment of fluctuations—and that our review is not entirely comprehensive. Our presentation, which largely follows the approach of Good and Desai (2014), neglects some of the fluctuations which are discussed at length in Fisher (2013), but enables us to streamline computations of average quantities such as *v*, *p*_fix_(*s*), *T_c_* and the site frequency spectrum.

A key quantity of interest is the distribution of fitnesses within the population, *f*(*X, t*), which gives the fraction of the population at absolute (log) fitness *X*. In our model, this fluctuating distribution evolves in time according to the nonlinear stochastic differential equation (Good and Desai, 2013),

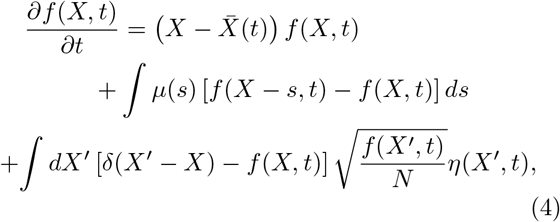

where 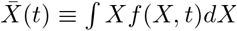, and *η*(*X, t*) is a Brownian noise term (Appendix A).

The last term in Eq. (4), which captures birth/death number fluctuations, is important. Without this term, even in the simple case of a single beneficial fitness effect, the solution *f*(*X, t*) would have a rate of fitness increase 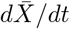 that grows without bound (Tsimring et al., 1996). In contrast, in stochastic simulations, Tsimring et al. (1996), Rouzine et al. (2003) and others have found that after an initial transient period, the distribution of fitnesses in a population attains a steady state “traveling wave” profile, *f*(*X*–*vt*) which moves through fitness space at a roughly constant rate 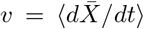; the steady state can be understood as a type of “mutation-selection balance” in which fitness variation, purged by selection, is repleted by new mutations (Desai and Fisher, 2007). In large populations, this average (or more precisely, *typical*) profile *f*(*x*) can then be approximated using a “quasi-deterministic” approach, in which

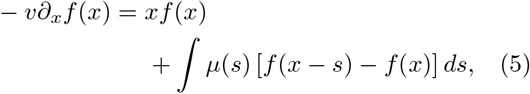

for *x* less than some cutoff *x_cut_*, and *f*(*x*) = 0 for *x* > *x_cut_* (Fisher, 2013; Good et al., 2012; Neher et al., 2010; Rouzine et al., 2008; Rouzine and Coffin, 2005; Rouzine et al., 2003; Tsimring et al., 1996). This defines a distribution which is normalized such that

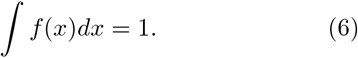

The solution to Eq. (5) will depend on the unknown average rate of mean fitness change *v*. To determine this quantity, we can express *v* in terms of the stochastic accumulation of new mutations,

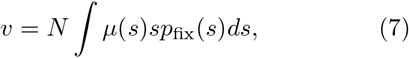

where *p*_fix_(*s*) describes the fixation probability of a mutation with fitness effect *s*. In our traveling wave framework, it is useful to express this fixation probability as

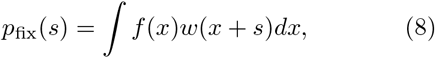

where *w*(*x*) denotes the fixation probability of a lineage founded by a single individual with relative fitness *x*. In this way, Eq. (8) averages over all of the possible fitness backgrounds on which a mutation can occur.

A key simplification is that *w*(*x*) can often be approximated by modeling the population as a collection of independent branching processes, which compete with each other only through the average rate of fitness change *v* (Fisher, 2013; Good and Desai, 2014; Good et al., 2012; Neher et al., 2010). Each of these lineages founds its own stochastic fitness wave, *g*(*x, t*), whose dynamics are described by a related differential equation,

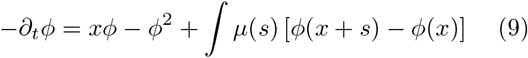

for the generating functional 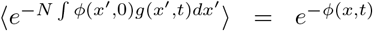 (Appendix A). The fixation probability *w*(*x*) then follows from

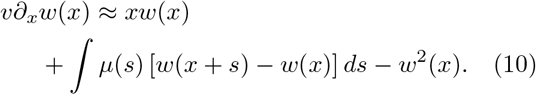

Together, Equations (5) and (10) can be solved given a particular *μ*(*s*) and an assumed rate of adaptation *v*. Using these solutions *f*(*x*) and *w*(*x*), Eq. (7) can be enforced as a self-consistency relation to determine *v* in terms of *N* and *μ*(*s*). In practice, it will be useful to enforce a related self-consistency condition,

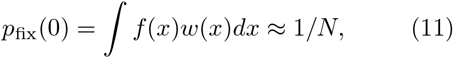

which ensures that neutral mutations fix with unbiased probability 1/*N*. Approximating the solutions to Eq. (5) and Eq. (10), or similar equations, given the parameters *N* and *μ*(*s*), has thus been the subject of extensive theoretical work (Fisher, 2013; Good and Desai, 2014; Good et al., 2012; Hallatschek, 2011; Neher et al., 2010).

A central complication is that Eq. (10) is nonlinear. Furthermore, the nonlinear *w*^2^ term is the only term in Eq. (10) which reflects the stochasticity of births and deaths. As a result, dropping the *w*^2^ term in Eq. (10) essentially amounts to a deterministic approximation, and leads to fundamentally wrong predictions. However, a *dominant balance* approach can be taken to treat this nonlinearity (Fisher, 2013; Good and Desai, 2014; Good et al., 2012; Neher et al., 2010). The basic idea is that because the fixation probability *w* must be an increasing function of *x*, at very large relative fitness (i.e. large *x*), *xw* and *w*^2^ are the dominant terms in Eq. (10). Thus at large *x* we have *w*(*x*) ≈ *x*. In contrast, for sufficiently small fitness, the nonlinear term must be negligible compared to the other terms. Thus at these small *x* we can neglect the *w*^2^ term in Eq. (10) and solve the linear equation

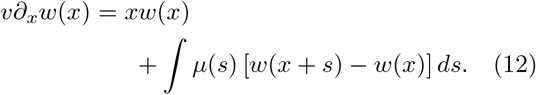

An approximation where the large-*x* and small-*x* solutions are simply *patched* together is then widely valid (Fisher, 2013; Good and Desai, 2014; Good et al., 2012). Specifically, a boundary *x_c_* is identified such that the large-*x* result (*w*(*x*) = *x*) is valid for *x* > *x_c_* and the small-*x* result (the solution of Eq. (12)) is valid for *x* < *x_c_*, without any intervening shoulder region (or more precisely, with a shoulder region that is narrow on relevant scales). A schematic of this patched solution *w*(*x*) is depicted in Fig. 1B. The details of how to determine the boundary *x_c_* can still be quite complicated and case-dependent (Fisher, 2013); however, in the regime of interest to us, a simple heuristic—ensuring that both *w*(*x*) and its derivative are continuous at the boundary *x_c_*—is adequate (Good and Desai, 2014). The boundary *x_c_* is often referred to as the *interference threshold*, since individuals with relative fitness *x* > *x_c_* fix with probabilities largely unaffected by interference (i.e. with their establishment probabilities ≈ *x*).

We note that solving Eq. (5) leads to negative values for *f*(*x*) at sufficiently large *x* (Fisher, 2013; Rouzine et al., 2008; Tsimring et al., 1996). This is an artifact of neglecting the fact that a finite set of fitnesses are represented in a population at any given time. To avoid this pathology, a common approach is to implement a “cutoff” by assuming that *f*(*x*) = 0 for *x* > *x_cut_* (Rouzine et al., 2008, 2003; Tsimring et al., 1996). The value *x_cut_* can then be interpreted as a typical maximum relative fitness of the individuals in a population, or as the fitness advantage of the “nose” of the population-wide fitness distribution. Consistent with Good and Desai (2014), Fisher (2013), and others, we take *x_cut_* = *x_c_* throughout. At a heuristic level, this choice can motivated by the fact that established lineages cannot typically exist above the interference threshold (if so, they would interfere); a more rigorous justification is given by Fisher (2013). Imposing a cutoff in *f*(*x*) can also be motivated using the tunable constraint framework introduced in Hallatschek (2011). In that work, the underlying model is changed in such a way to yield moment closure of the equations for the stochastic time-dependent fitness distribution; the resulting analog of *f*(*x*) does not display the above-mentioned pathological behavior, and instead exhibits an exponential decay beyond a fitness scale related to *x_c_* (suggesting that imposing a cutoff at *x_c_* is reasonable). We note further that the “tunable constraint” imposed by Hallatschek (2011) turns out to be equivalent to the requirement *p*_fix_(0) = 1/*N* that we enforce to ensure a self-consistent rate of adaptation *v* (Good et al., 2012). Ultimately, our results are relatively insensitive to whether a cutoff is taken in *f*(*x*) or a tunable constraint model is assumed.

Analysis of the linearized equations (5) and (12) is still not straightforward, however, since both equations contain mutation terms which are *nonlocal*. While exact solutions for suitably defined Laplace transforms 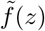 and 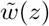 can be obtained (Fisher, 2013), the subsequent inversion of these transforms requires approximation. A related strategy has been to approximate Eq. (5) and Eq. (12) by a set of *local* differential equations, which can then be solved straightforwardly. For example, in the infinitesimal regime, the relevant fitness effects *s* are assumed sufficiently small to approximate

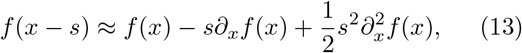

and

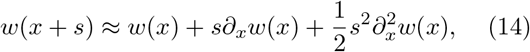

in Eq. (5) and Eq. (12), respectively. The resulting *f*(*x*) and *w*(*x*) can be obtained using the Airy equation (Cohen et al., 2005; Hallatschek, 2011; Tsimring et al., 1996). Rouzine et al. (2003) and Rouzine et al. (2008) employ a similar approach, instead Taylor approximating the logarithm of *f*(*x* – *s*).

Our focus in this article is on a related approach to approximate Eq. (5) and Eq. (12) by a set of local equations, used by Fisher (2013) to study the “high-speeds” regime. In Section IV, we begin by reviewing this approximate calculation of *f*(*x*) and *w*(*x*). We proceed to provide novel solutions to a number of questions within a *moderate selection, strong mutation* (MSSM) regime, within which this approximate calculation is valid, and which includes both the “high-speeds” regime and the infinitesimal regime as special cases. We first solve for divergence-related quantities including fixation probabilities *p*_fix_(*s*) and the average rate of fitness change *v*, and discuss conditions of validity of our approach that are obtained by ensuring *f*(*x*) and *w*(*x*) are well-approximated in the region of *x* making a dominant contribution to these quantities. We then consider statistics of genetic diversity within the MSSM regime, with a focus on the site frequency spectra of selected mutations and of neutral mutations. By calculating the neutral site frequency spectrum, we demonstrate a partial correspondence between genealogies within the MSSM regime and those of the Bolthausen-Sznitman coalescent (BSC), and analytically describe departures from the BSC that are apparent in the low-frequency portion of the site frequency spectrum. Finally, we conclude Section IV by discussing the various fitness and time scales that emerge in our analysis, and provide a heuristic description of the dynamics of lineages within the MSSM regime.

We summarize some of our key notation used throughout in Table I.

**TABLE I.**
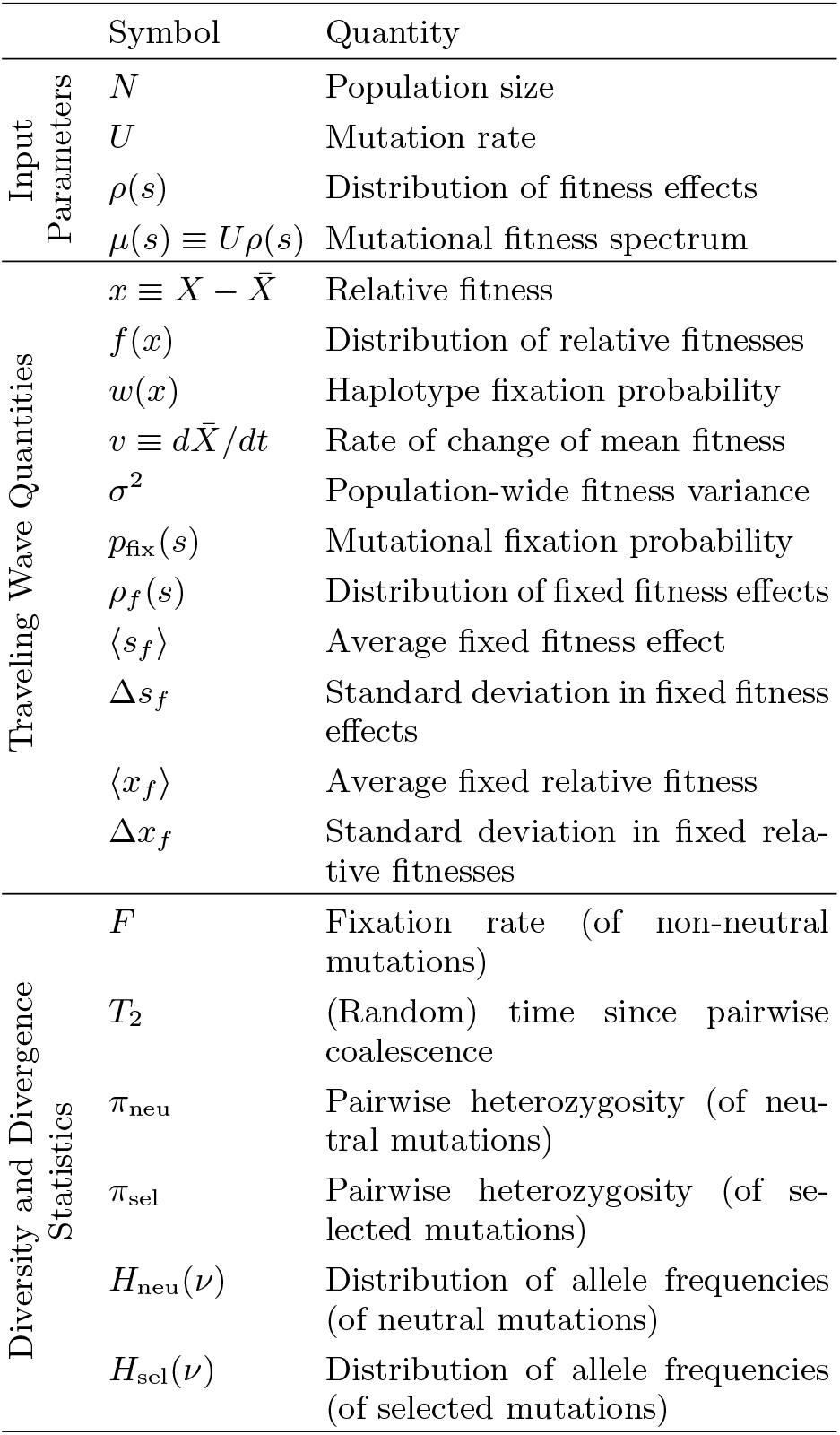
Notation Used

**TABLE II.**
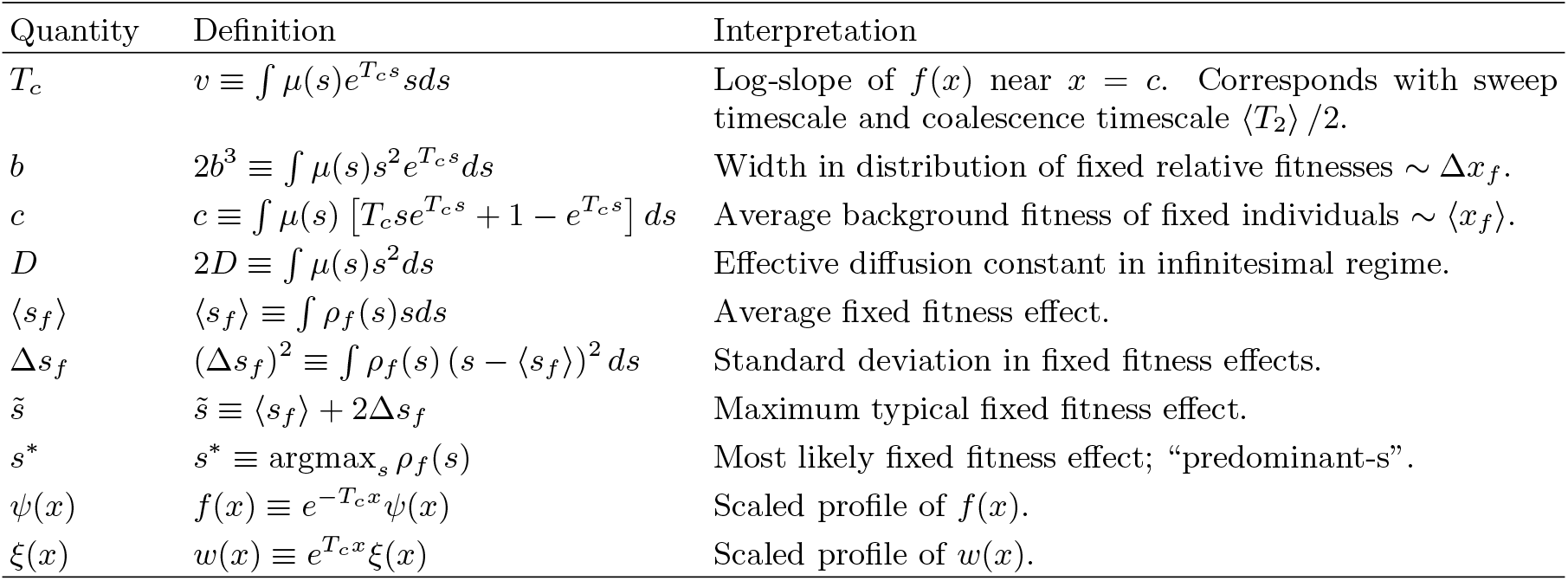
Key Definitions

## IV. ANALYSIS

As noted above, our approach is based on a key approximation employed by Fisher (2013) to study the “high-speeds” regime. The key idea is that we can approximate *f*(*x* – *s*) and *w*(*x*+*s*) in Eq. (5) and Eq. (10), respectively, by first pulling out rapid exponential prefactors varying at the appropriate rate, and then performing Taylor approximations of the remaining factors. To do so, we define

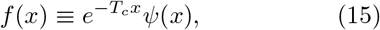

and

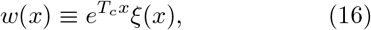

with *T_c_* defined by

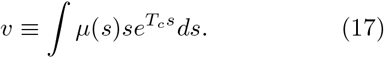

Note that *T_c_* is not a parameter of our model; Eq. (17) uniquely defines a specific *T_c_* given the model parameters *N* and *μ*(*s*). If *μ*(*s*) falls off *slower* than exponentially with *s* at large positive *s*, then *T_c_* is not well-defined and our approach breaks down; in the marginal case of an exponential DFE of beneficial mutations (with mean effect *s_b_*), Eq. (17) implies that *T_c_s_b_* < 1. We will later show in Section V that, within the MSSM regime, the quantity *T_c_* as defined in Eq. (17) is approximately equal to 〈*T*_2_〉 /2—one half the average time since pairwise coalescence among individuals in a population—which motivates our interpretation of *T_c_* as a *coalescence* timescale. However, we emphasize that *T_c_* is a derived quantity, defined by Eq. (17), which we will relate to the underlying parameters *N* and *μ*(*s*) in Section IV.A. [For example, for the case of single beneficial fitness effect, *μ*(*s*) = *Uδ*(*s* – *s_b_*), we will have 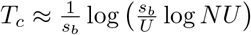 provided that *T_c_s_b_* ≫ 1.]

Plugging Eq. (15) and Eq. (16) into Eq. (5) and Eq. (10), we find that for *x* < *x_c_*, *ψ*(*x*) and *ξ*(*x*) satisfy

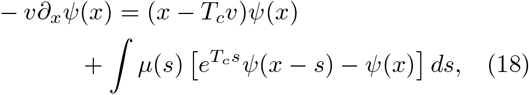

and

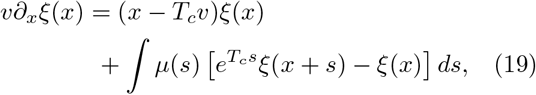

respectively. Taylor expanding *ψ*(*x* – *s*) in Eq. (18) and keeping the lowest two nonzero orders in *s*, we find

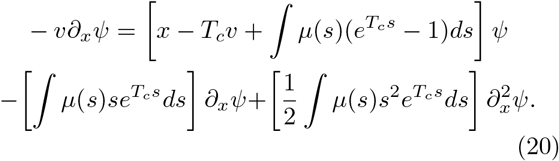

It will be useful to define the fitness scales *b* and *c* according to

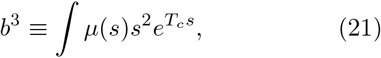

and

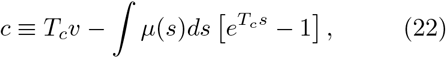

respectively, which enable us to rewrite Eq. (20) in the compact form

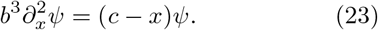

Similar manipulations can be carried out to approximate *ξ*(*x*) in Eq. (19), with both equations solved by

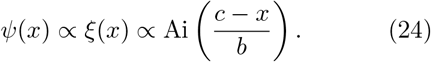

This implies that for *x* < *x_c_*,

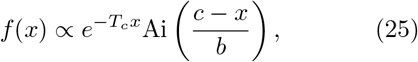

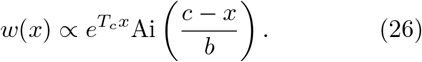

As discussed in Section III, we will take *w*(*x*) ≈ *x* and *f*(*x*) ≈ 0 for *x* > *x_c_*. Provided that *T_c_b* ≫ 1, the transition between these solutions will occur over a narrow boundary layer, such that *x_c_* can be determined by enforcing continuity of *w*(*x*) and *w*′(*x*) at *x* = *x_c_* (as done by Good et al. (2014), for instance). This calculation is presented in Appendix B, with the result

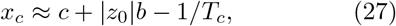

where *z*_0_ ≈ –2.34 is the least negative zero of Ai(*z*). We note that *x_c_* is obtained by Fisher (2013) in the “high-speeds” regime using an alternative “solvability condition” for *w*(*x*) which yields the same result.

### Relation to infinitesimal approximation

We note that in the limit *T_c_s* → 0 for relevant fitness effects *s* (with *T_c_b* ≫ 1 fixed), the solutions *f*(*x*) and *w*(*x*) in Eq. (25) and Eq. (26), as well as *x_c_* in Eq. (27), reduce to the corresponding quantities obtained using the infinitesimal approximation. In this limit,

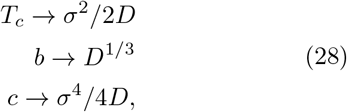

where here, *σ*^2^ ≡ *v* – ∫ *μ*(*s*)*sds* corresponds to the population-wide fitness variance, and 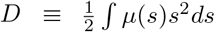 corresponds to a mutational *diffusion* constant (an interpretation of which we provide in Subsection VI). The above approximation can be considered a *generalized infinitesimal* approximation; we further discuss the relation between our approximation detailed above and the infinitesimal approximation in the following Subsections.

#### A. The relation between *T_c_* and *N*

Our derivation above made use of the phenomenological quantities *T_c_*, *b*, *c* and *v*, which are functions of the underlying parameters *N* and *μ*(*s*). We now derive an additional equation relating these quantities, which allows us to solve for *T_c_* (and thus *v*, as well as *b* and *c*) in terms of *N* and *μ*(*s*). To do so, we enforce the condition ∫ *f*(*x*)*w*(*x*)*dx* = 1/*N* in Eq. (11), using the approximate *f*(*x*) and *w*(*x*) in Eq. (25) and Eq. (26), respectively. We emphasize that the expressions in Eq. (25) and Eq. (26) are only *local* approximations, valid within some range of *x*. In the next Subsection, we will obtain and discuss conditions which ensure that Eq. (25) and Eq. (26) are approximately valid within the important region dominating 1/*N* = ∫ *f*(*x*)*w*(*x*)*dx*. The quantity *Nf*(*x*)*w*(*x*) can be interpreted as a distribution of fitnesses of future common ancestors, and given *f*(*x*) and *w*(*x*) in Eqs. (25) and (26), evaluates to

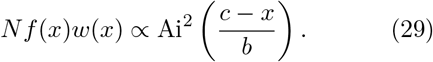

The integral ∫ *f*(*x*)*w*(*x*)*dx* thus receives a dominant contribution from the region 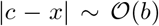. This motivates us to refer to the region 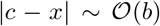 probed by our condition of validity as the *fixation class*, since collectively, this region of fitness space produces a future common ancestor of the population with probability 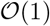, despite comprising a small fraction of the total population.

To evaluate the integral in Eq. (11), we need the overall constants of proportionality of *f*(*x*) and *w*(*x*). Recall that the constant of proportionality for *w*(*x*) was fixed by matching to the solution *w*(*x*) ≈ *x* at *x* = *x_c_*. On the other hand, the overall constant of proportionality of *f*(*x*) must be determined by the normalization condition ∫ *f*(*x*)*dx* = 1. However, while our local approximation for *f*(*x*) in Eq. (25) is valid near the nose, it is not necessarily valid throughout the entire normalization integral ∫ *f*(*x*)*dx*, which is dominated by fitnesses near the mean. Instead, we show in Appendix C that an analogous local approximation can be applied to the Laplace transform of *f*(*x*), which allows us to obtain the relevant normalization,

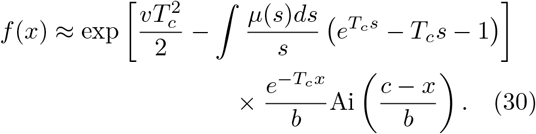

Using this expression, we can obtain a relation between *T_c_* and *N*, *μ*(*s*) and *x_c_*,

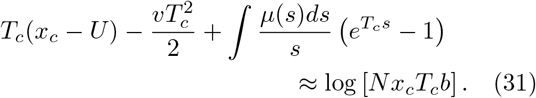

(Appendix B). By combining Eq. (27) and Eq. (31), along with the definitions for *T_c_*, *b* and *c* in Eqs. (17), (21) and (22) it is possible to solve for *T_c_* in terms of *N* and *μ*(*s*). We therefore defined a simple numerical routine to solve Eq. (27) and Eq. (31) for a given choice of *N* and *μ*(*s*) (Appendix I). Although we discuss analytical approximations to this solution in Subsection IV.C, we use this numerical solution for comparison with simulations throughout the rest of this paper. We note that Eq. (31) can be rearranged to provide the relationship between *T_c_*/*N* and the combinations *T_c_U* and the distribution of scaled effects *T_c_s*. In this way, our approximation is well suited to describing the dynamics corresponding to given values of the quantities *T_c_U* and *T_c_s*, parameter combinations more often inferred in natural settings (albeit often problematically, as our analyses show) than *NU* and *Ns*.

We illustrate the fact that Eq. (30) is only a locally valid approximation—and the importance of normalizing *f*(*x*) appropriately—in Fig. 2. There, we compare our predicted *f*(*x*) with the same quantity observed in simulations, for three example parameter choices. In the same Figure, we also plot histograms of future common ancestors fitnesses–that is, the empirically measured distribution *Nf*(*x*)*w*(*x*)—obtained from simulations. We can see that the prediction for *f*(*x*) in Eq. (30) matches simulation results well in the region dominating ∫ *f*(*x*)*w*(*x*)*dx*. Outside, this region, however, the prediction in Eq. (30) breaks down, and would yield an incorrect constant of proportionality if normalized “directly”—particularly in the region of the MSSM regime that lies outside the infinitesimal regime. For comparison, we also plot predictions for *f*(*x*) obtained using the infinitesimal approximation (for details, see Appendix I), as well as a numerical saddle point approximation we present in Appendix C. While our numerical saddle point approximation does yield improved accuracy in predicting the simulated *f*(*x*) throughout its “bulk”, it has a negligible impact on global quantities of interest such as *v* and *p*_fix_(*s*), which are dominated by the behavior of *f*(*x*) and *w*(*x*) in the fixation class. We therefore use only our analytical prediction for *f*(*x*) in Eq. (30) throughout the remainder of this article.

**FIG. 2.**
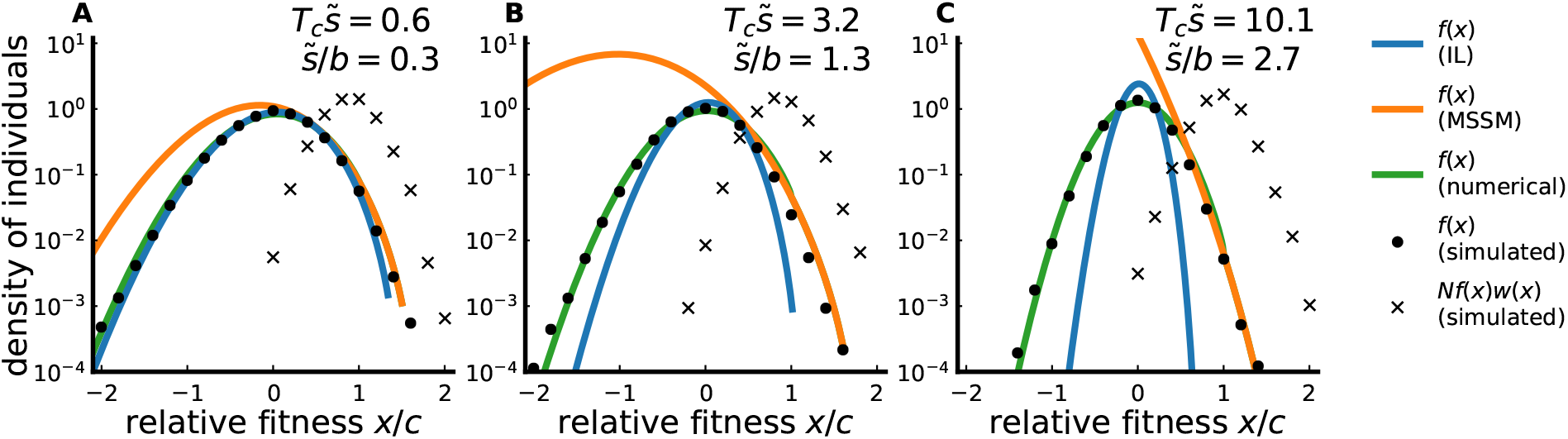
Averaged distributions of relative fitnesses, *f*(*x*), for simulated populations subject to beneficial mutations with an exponential DFE, with values of 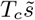 and 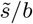 denoted above. A precise definition of 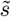 is given in Subsection IV.B. Note the logarithmic scale of the vertical axis. Filled circles represent the simulated distribution of relative fitnesses, *f*(*x*), as obtained by averaging over measurements from 490 different epochs. Black ‘x’ markers represent the simulated distribution of future common ancestor relative fitnesses, *Nf*(*x*)*w*(*x*). Blue and orange lines denote theoretical predictions for *f*(*x*) obtained with the infinitesimal (IL) and MSSM approximations, respectively; green lines denote predictions for *f*(*x*) obtained using the numerical saddle point approximation detailed in Appendix C. In all cases *f*(*x*) is best-predicted by the numerical saddle point approximation, but the MSSM approximation adequately predicts the behavior of *f*(*x*) within the region of *x* dominating ∫ *f*(*x*)*w*(*x*)*dx*. In panel A the infinitesimal approximation adequately predicts *f*(*x*) for all *x*, which is expected since 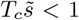. Panels B and C correspond to larger values of 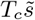 for which the infinitesimal approximation begins to break down, particularly in the region of *x* dominating ∫ *f*(*x*)*w*(*x*)*dx*.

#### B. Conditions of validity: a “moderate selection, strong mutation regime”

The approximate solutions *f*(*x*) and *w*(*x*) given in Eq. (25) and Eq. (26) are valid only within a limited region of fitness space. Our primary motivation in solving for *f*(*x*) and *w*(*x*) is that these quantities can be used to compute other dynamical quantities such as *v* and *ρ_f_*(*s*). These computations involve integrals of the form ∫ *f*(*x*)*w*(*x* + *s*)*dx* (and our computations of genetic diversity statistics in Section V involve similar integrals as well). We can thus obtain conditions of validity of our approach by demanding that both *f*(*x*) and *w*(*x*) are well-approximated in the region of fitness space dominating these integrals. We will find that these conditions are most readily expressed in terms of *ρ*(*s*) and the derived quantities *T_c_* and *b*, although these conditions can straightforwardly be expressed in terms of the model parameters *N* and *μ*(*s*) using the relation between *T_c_* and *N* given above.

In particular, as noted in the previous Subsection, ∫ *f*(*x*)*w*(*x*)*dx* is dominated by the region 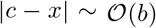. The same region dominates the integral used to compute *ρ_f_*(*s*),

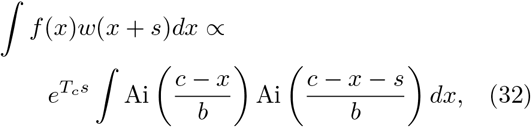

as long as *s* ≪ *b*. For *s* ≪ *b*,

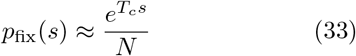

follows from Eq. (32), and 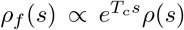 (Appendix D). We therefore obtain a condition of validity of our approximation by ensuring that *f*(*x*) and *w*(*x*) are well-approximated within the region 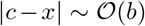. We will see below that this condition requires that *s* ≪ *b* for typical fixed effects *s* (i.e. for the region of *s* which dominates ∫ *s*^2^*ρ_f_*(*s*)*ds*), so that *f*(*x*) and *w*(*x*) are also well-approximated within the region dominating ∫ *f*(*x*)*w*(*x* + *s*)*dx* for relevant *s*.

We obtain this condition of validity in Appendix C. The basic idea is to ensure that the inverse Laplace transforms of the linearized *f*(*x*) and *w*(*x*) reduce to the approximate expressions in Eq. (25) and Eq. (26). For relevant *x*, this will be true provided that a particular term can be neglected. By ensuring, for 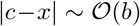, that this additional term yields small corrections near saddle points of the inverse Laplace integral, we obtain the condition

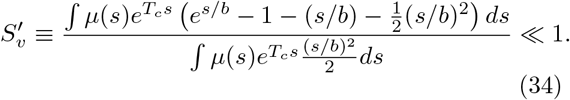

Note that the condition *T_c_b* ≫ 1 is required to justify the dominant balance approximation for *w*(*x*). These conditions can be applied to determine the suitability of the above approximations for populations subject to beneficial mutations, deleterious mutations, or some combination of the two.

As anticipated above, under relatively mild assumptions on *ρ*(*s*), the condition 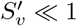 is satisfied if

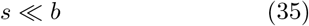

for *s* dominating ∫ *ρ_f_*(*s*)*s*^2^*ds* (i.e. if typical fixed fitness effects *s* are much smaller than *b* in magnitude). For bookkeeping purposes, it is useful to define a maximum typical fixed fitness effect, 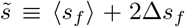, where 〈*s_f_*〉 and Δ*s_f_* are the average and standard deviation, respectively, of fixed mutation effects. From Eq. (29), we also note that Δ*x_f_* ~ *b* (where 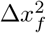 denotes the fitness variance of future common ancestors). Given these definitions, the conditions 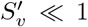 and *T_c_b* ≫ 1 can then be written more compactly as

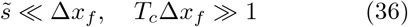

These conditions have dynamical interpretations which we discuss in Subsection VI. Notably, because 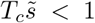 is not required by our conditions, selection on single mutations need not be weak. Because 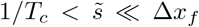 (or 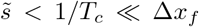) is permitted, we refer to the regime of validity of our approximation as a *moderate selection, strong mutation* (MSSM) regime. [“Moderate selection” refers to the requirement that 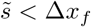, while “strong mutation” refers to the requirement that clonal interference is strong.]

#### C. Specific example DFEs

We can gain more intuition about the MSSM regime by solving for *T_c_* (and therefore *v*) for a few concrete example DFEs.

### Single beneficial fitness effect

The simplest such scenario is that of a single beneficial fitness effect *s_b_*. We begin by noting that when *T_c_s_b_* ≪ 1 Eq. (31) reduces to the well-known results

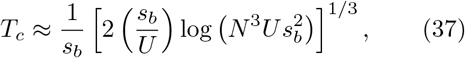

and

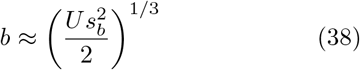

from the infinitesimal limit (Good et al., 2014; Neher and Hallatschek, 2013). We can see that Eq. (37) is self-consistently valid (i.e. *T_c_s_b_* ≪ 1, *T_c_b* ≫ 1 and *s_b_* ≪ *b*) when

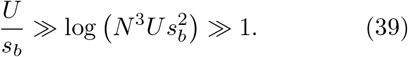

Note that this condition requires more than just *U* ≫ *s_b_*; *U*/*s_b_* must also be larger than the large parameter combination 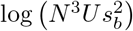.

In the opposite regime *T_c_s_b_* ≫ 1, Eq. (31) reduces to

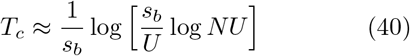

and

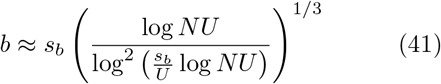

which coincides with the “high-speeds” regime considered by Fisher (2013) (Appendix B). This is self-consistently valid when

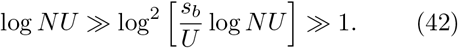

By combining Eq. (42) and Eq. (40), we can see that there is a relatively smooth crossover between the infinitesimal and “high-speeds” regimes when *U*/*s_b_* ~ log *NU*, where 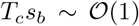. For fixed *U* and *s_b_*, the infinitesimal limit always breaks down for sufficiently large *N*, while the “high-speeds” regime is eventually valid for any *s_b_* and *U*.

### Distributions of beneficial fitness effects

More generally, as long as the DFE *ρ*(*s*) falls off *faster* than exponentially with large positive *s*, convergence to the MSSM regime is obtained in the limit *N* → ∞. In this limit, previous work has shown that the relevant integrals over *μ*(*s*)*e^T_c_s^* become increasingly sharply peaked around a characteristic value, such that

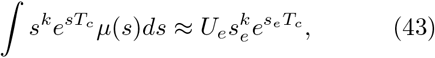

where *s_e_* coincides with the predominant fitness effect *s*\* ≡ argmax*_s_ρ_f_*(*s*) (Fisher, 2013). This shows that short-tailed DFEs can be approximated by an effective DFE with *μ*(*s*) = *U_e_δ*(*s* – *s_e_*) (Desai and Fisher, 2007; Fisher, 2013; Good et al., 2012; Hegreness et al., 2006). Moreover, under rather general conditions, one can show that *s*\* becomes increasingly small compared to *b* as *N* (and therefore *T_c_*) increases, so that the conditions of validity of the MSSM approximation will be satisfied (Appendix G).

On the other hand, if *ρ*(*s*) falls off slower than exponentially with large positive *s*, the integrals in Eq. (43) no longer converge, and the MSSM approximation cannot be applied. The case of an exponential DFE (with mean effect size 〈*s*〉) is a marginal case, since convergence will depend on the relative values of *T_c_* and 〈*s*〉, or equivalently, of *T_c_* and 〈*s_f_*〉. When *T_c_* 〈*s_f_*〉 ≪ 1, the MSSM approximation reduces to the infinitesimal approximation, as in the case of a single beneficial effect. In the opposite case that *T_c_* 〈*s_f_*〉 ≫ 1, we find that

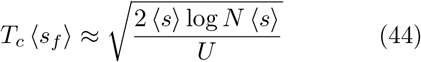

and

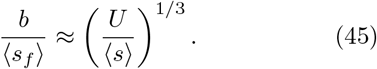

(Appendix G). The conditions *T_c_* 〈*s_f_*〉 ≫ 1, 〈*s_f_*〉 ≪ *b* and *T_c_b* ≫ 1 are then jointly satisfied, and the MSSM approximation is valid, when

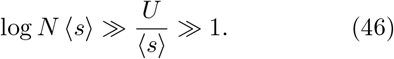

Note that since *T_c_* 〈*s_f_*〉 ≈ *T_c_*〈*s*〉 /(1 – *T_c_*〈*s*〉) for an exponential DFE, (1 – *T_c_*〈*s*〉) is a small and positive quantity in this case, such that the relevant integrals converge. Note that the conditions in Eq. (46) differ from those for a single beneficial fitness effect in Eq. (42) in that *U* ≫ 〈*s*〉 is now explicitly required, no matter how large the value of *N*. The opposite case, in which 〈*s*〉 > *U*, cannot be described using the MSSM approximation, and remains only partially understood; for a discussion of this case, see Fisher (2013).

### Deleterious mutations in adapting populations

The previous two examples have mostly recapitulated earlier results for beneficial mutations in the infinitesimal (Cohen et al., 2005; Hallatschek, 2011; Neher and Hallatschek, 2013; Tsimring et al., 1996) and high-speed regimes (Fisher, 2013). A key advantage of our MSSM approximation is that it can also be applied to scenarios with large numbers of deleterious mutations, where few analytical results are currently available.

As an example, we first consider a scenario where beneficial and deleterious mutations have the same effect size *s*, and occur at rates *U_b_* and *U_d_* respectively. The infinitesimal approximation will once again apply in the limit that *T_c_s* ≪ 1, but qualitatively new behavior starts to emerge when *T_c_s* ≫ 1. In this case, a useful simplification occurs in Eq. (31): the contributions to the left-hand side are exponentially suppressed for deleterious mutations, with the exception of the *U_d_*/*s* term. As a result, the overall solution for *T_c_* reduces to the purely beneficial case in Eq. (40), but with an *effective population size*

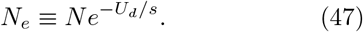

in place of *N*. In particular, the conditions of validity in Eq. (42) imply that the MSSM approximation will be valid for sufficiently large *N* for any choice of *U_b_*, *U_d_* and *s*. Note that while we have assumed that *s_d_* = *s_b_* = *s* for simplicity above, this same argument can be generalized to unequal selection strengths as long as *T_c_s_d_* ≫ 1. For example, if *T_c_s_b_* ≪ 1, then *T_c_* will instead be described by the infinitesimal approximation in Eq. (37), with *N_e_* = *Ne*^−*U_d_*/*s_d_*^.

We emphasize that the effective population size in Eq. (47) does *not* denote a coalescence timescale, as is often implied in the literature. Indeed, the asymptotic expressions in Eqs. (40) and (37) show that *T_c_* will typically be much less than *N_e_* in the MSSM regime. The interpretation of such a reduced effective population size is clear, however: strongly deleterious mutations rarely fix, but they are not purged from a population immediately. The effective population size in Eq. (47) resembles the classical prediction for the number of mutation-free individuals that would exist at mutation-selection balance (Haigh, 1978). Our results suggest that this simple approximation continues to apply in certain non-equilibrium settings as well, even when the deleterious fitness costs are small compared to the typical fitness variation in the population. Although these strongly deleterious mutations are unable to fix, they can still affect the evolutionary dynamics through the size of the unloaded class, similar to the “background selection” (Charlesworth, 1994) or “ruby in the rubbish” (Peck, 1994) behavior that has been observed for single beneficial mutations. We emphasize that this effective population size approximation was a direct consequence of our mathematical expressions in Eqs. (21) and (31). This provides a more rigorous justification for previous ad-hoc approaches that assumed that strongly deleterious mutations can be treated in this way (Good et al., 2014; Söderberg and Berg, 2007).

### Background selection

Finally, we can also apply our framework to the case of purely deleterious mutations (often referred to as “background selection”). For simplicity, we will focus on the well-studied case where deleterious mutations have a single fitness cost *s_d_*. As noted in previous studies (Good et al., 2014; Neher and Hallatschek, 2013; Neher et al., 2013), this scenario is well-described by the infinitesimal limit when *T_c_s_d_* ≪ 1, with *s_d_* replacing *s_b_* in Eq. (37). In the opposite case where *T_c_s_d_* ≫ 1, our MSSM approximations yield an alternative solution, in which

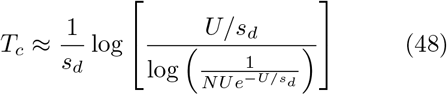

and

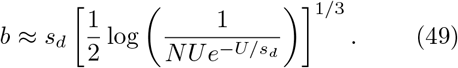

This solution is self-consistently valid (*T_c_s_d_* ≫ 1, *s_d_* ≪ *b* and *T_c_b* ≫ 1) when

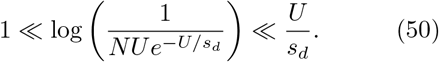

For fixed values of *U* and *s_d_*, this condition will always be violated for sufficiently large population sizes (*NUe*^−*U/s_d_*^ ≳ 1). This breakdown is consistent with previous theory (Charlesworth, 1994; Cvijović et al., 2018; Good et al., 2014) which predicts that the large-*N* limit of the background selection model approaches a nearly neutral regime, with a coalescent timescale,

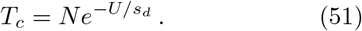

Interestingly, Eq. (50) shows that the MSSM approximation can remain valid even for arbitrarily large values of *T_c_s_d_*, as long as the corresponding values of *U/s_d_* are also sufficiently large. Thus, even for purely deleterious mutations, the MSSM approximation does not necessarily require that all mutations have weak effects (i.e. that *T_c_s_d_* ≪ 1). In principle, strong clonal interference can occur even for large scaled fitness effects (*T_c_s_d_* ≳ 1), but only in an increasingly narrow region of the underlying parameter space.

This last point suggests an alternative way of looking at the relation between *T_c_* and *N* in Eq. (31). In addition to solving for *T_c_* as a function of the bare parameters *N*, *U* and *s_d_*, it is also possible to directly solve for *NU* as a function of the phenomenological variables *T_c_s_d_* and *T_c_U*:

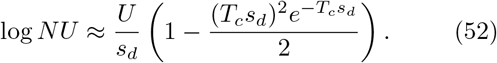

This expression gives the underlying value of *NU* that would be required for the MSSM regime to apply for a given value of *T_c_s_d_* and *T_c_U*. This alternative view of the parameter space, in which *T_c_s_d_* and *T_c_U* are the “independent” parameters, is often more natural in applied settings, where the phenomenological parameters are usually estimated from contemporary patterns of genetic diversity. This perspective will be useful for our discussion of nonsynonymous site frequency spectra below.

#### D. Simulation results

To complement our analytical predictions, we simulated populations subject either to only beneficial mutations (adapting populations) or only deleterious mutations (ratcheting populations). Simulated parameters *N* and *μ*(*s*) are chosen to correspond with a grid with linearly spaced *T_c_b* values, and with logarithmically spaced *T_c_*〈*s_f_*〉 values (for adapting populations) or *T_c_*〈*s*〉 values (for ratcheting populations), subject to the constraints *T_c_b* > 1,1 < *NU* < 10^5^, 1 < *N* 〈*s*〉 < 10^3.5^ and *U*/ 〈*s*〉 ≤ 10^4^. For ratcheting populations we add a further constraint that *U*/ 〈*s*〉 > 1. These constraints are chosen both to limit attention to the MSSM regime (and the region of parameter space where it begins to break down) and to ensure feasibility of our individual-based simulations. For each point on these constrained grids, we (separately) simulated populations subject to an exponential DFE, and to a stretched-exponential DFE with steepness parameter *β* = 2. Details of the numerical implementation of these simulations and choice of parameters are given in Appendix H; in Fig. S1 we plot our constrained grid of simulation parameters in the space of *NU* vs. *N*〈*s*〉.

In Fig. 3, we plot our constrained grid in the space of *T_c_b* vs. 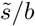, with the color of each point denoting the accuracy of either the MSSM approximation or infinitesimal approximation in predicting the rate of change *v* in the mean fitness. For clarity, we include only populations subject to an exponential DFE in Fig. 3. Plotting our simulation grid in the space of *T_c_b* vs. 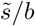 enables us to verify that predictions of the MSSM approximation are accurate when our conditions of validity are met (roughly speaking, when *T_c_b* ≫ 1 and 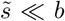). From Fig. 3, we can see that our predictions for *v* are reasonably accurate even for 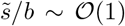, as long as *T_c_b* > 1. Moreover, we can see that the infinitesimal approximation breaks down even for small 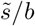 when 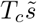 is large.

**FIG. 3.**
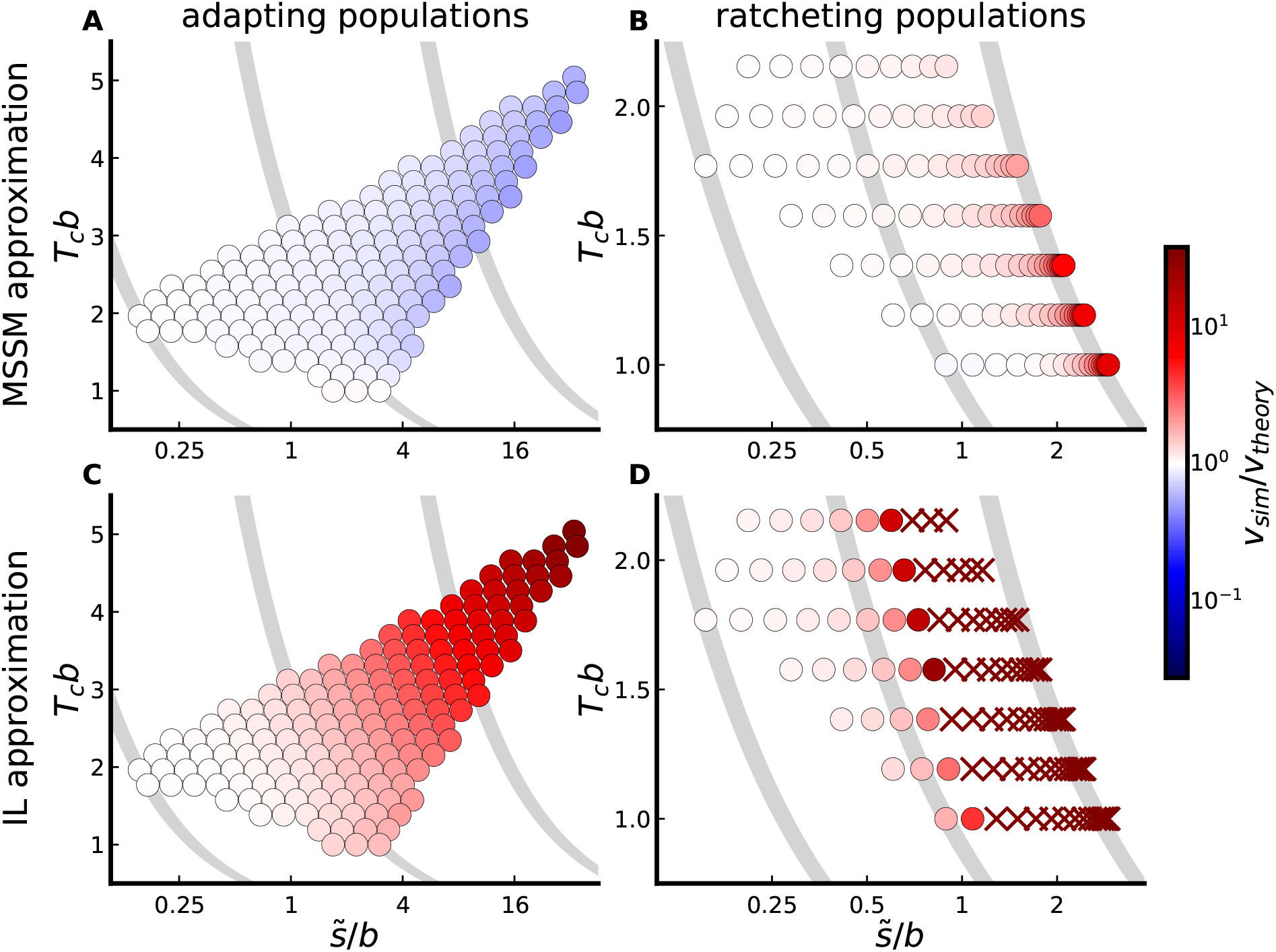
Comparison of theoretical predictions for the rate of change in mean fitness, *v*, to the corresponding rates observed in simulations, for populations subject to an exponential DFE of beneficial mutations or deleterious mutations. The color of each point denotes the ratio of simulated to predicted *v* for a set of parameters (with red and blue indicating that theory underestimates or overestimates *v*, respectively, according to the scale at right). In panels A and B, predictions are obtained using the MSSM approximation, while in panels C and D, predictions are obtained using the infinitesimal (IL) approximation. The location of a point along the vertical axis denotes the value of *T_c_b* for its corresponding set of parameters (with *T_c_* and *b* computed using the MSSM approximation in all panels), and the horizontal axis denotes the value of 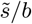 for that set of parameters (with 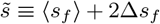); curves of constant 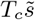 are denoted in gray. Panels A and C involve simulations of populations subject only to beneficial mutations (adapting populations), and panels B and D involve simulations of populations subject only to deleterious mutations (ratcheting populations). Simulated parameters are those lying on the constrained grid described below, and depicted in the space of *NU* vs. *N* 〈*s*〉 in Fig. S1. The ‘x’ markers in D denote populations for which the infinitesimal approximation predicts a rate of fitness change of the incorrect sign.

To visualize the quantitative agreement between simulations and our predictions more directly, we plot the ratio *v*_sim_/*v*_theory_ as a function of the single quantity 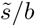 in Fig. 4, with points colored according to their values of *T_c_b*. We include, in Fig. 4, simulated populations subject to a *β* = 2 stretched-exponential DFE, as well as simulated populations subject to an exponential DFE. In the same Figure, we compare theoretical predictions of the *fixation rate* of new mutations, given by

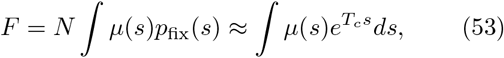

as well as the average fixed effect 〈*s_f_*〉 and standard deviation Δ*s_f_* in fixed effects, to measurements of these quantities in simulations. Predictions for the quantities 〈*s_f_*〉 and Δ*s_f_* are obtained using the simulated *ρ*(*s*) and *p*_fix_(*s*) in Eq. (33). For each of these quantities, we observe highly quantitative agreement between simulations and theory for small and moderate values of 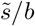.

**FIG. 4.**
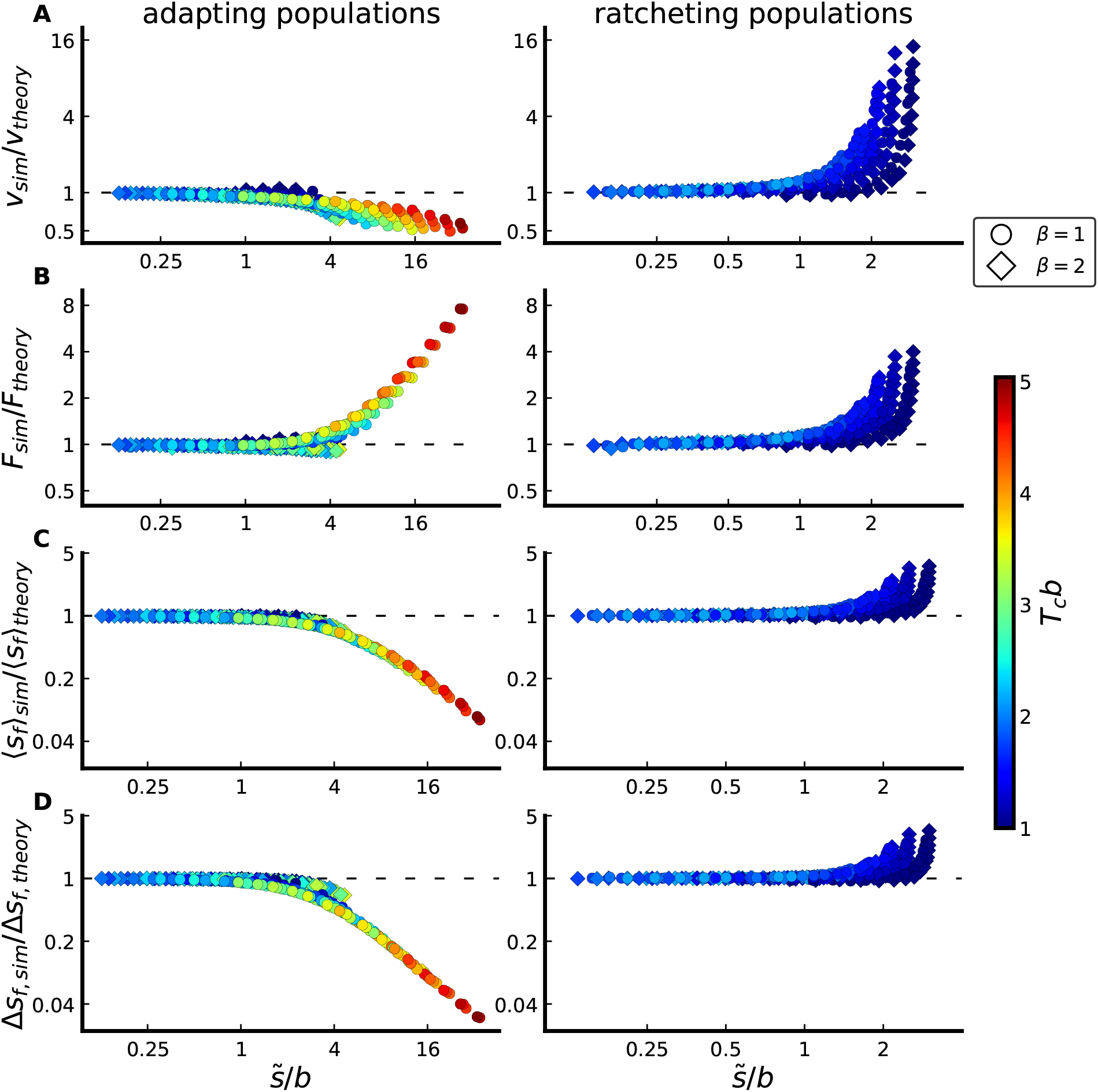
Comparison between simulated and predicted rate *v* of fitness change (A), rate *F* of mutation accumulation (B), average fixed effect 〈*s_f_*〉 (C), and standard deviation Δ*s_f_* in fixed effects (D). Populations in the left column are subject to only beneficial mutations; populations in the right column are subject to only deleterious mutations. Predictions are obtained using the MSSM approximation; quantitative agreement between simulations and predictions is obtained as long as 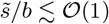. Simulated sets of parameters lie on the same (constrained) grid considered in Fig. 3, although here we include, for each point on that grid, both an exponential DFE (circles) and *β* = 2 stretched-exponential DFE (diamonds).

## V. STATISTICS OF GENETIC DIVERSITY

We now consider statistics of genetic diversity within the MSSM regime. Our central quantity of interest will be the frequency spectrum *h_s_*(*ν*) for a single site with fitness effect *s*, which is defined such that

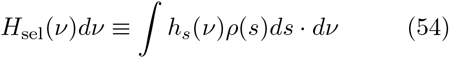

gives the expected number of selected (non-neutral, including both beneficial and deleterious) mutations per site with frequencies between *ν* and *ν* + *dν*. We will also consider an analogous aggregate quantity

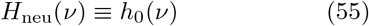

defined for a subset of putatively neutral sites (e.g. synonymous sites, short introns, etc). These site frequency spectra are important statistics that are often used to make inferences regarding the evolutionary forces acting within a population (Nielsen, 2005). Several empirical studies have drawn inferences on the presence and strength of selection acting in a population based on differences in the site frequency spectrum among synonymous (i.e. mostly neutral) and nonsynonymous (i.e. more selected) mutations (Eyre-Walker et al., 2006). The key idea underlying these approaches is simple: because synonymous and nonsynonymous sites (which are interdigitated throughout the genome) share the same demographic history, differences in their patterns of diversity can be attributed to (positive or negative) selection on nonsynonymous mutations (McDonald and Kreitman, 1991). The ability to make inferences using this line of thinking requires predictions for *h_s_*(*ν*) (Hartl et al., 1994; Sawyer and Hartl, 1992), which are lacking for rapidly evolving populations.

In the following subsections, we develop analytical predictions for these neutral and selected site frequency spectra in the MSSM regime, noting important departures from the classical intuition in Eq. (1). We demonstrate that *h_s_*(*ν*) and *h*_0_(*ν*) are simply related by a constant factor above a characteristic frequency *ν_c_* ≪ 1, which is not simply summarized by the scaled selection coefficient *T_c_s*. Building on previous work (Desai et al., 2013; Fisher, 2013; Neher and Hallatschek, 2013), we also demonstrate a partial correspondence between the genealogies in our model and the Bolthausen-Sznitman coalescent, and we analytically describe the departures from the BSC SFS that can be observed at low frequencies. Compared to the BSC SFS, we find that the functional form of these departures provides substantially greater power to distinguish different parameter combinations. Finally, we compute the pairwise heterozygosity for selected and neutral mutations, and demonstrate a correspondence between the pairwise coalescence time 〈*T*_2_〉 and the quantity *T_c_* defined in Eq. (17). In the Discussion, we consider implications of our results in the context of the population genetic inference methods described above.

### 1. The site frequency spectrum: basic formalism

To calculate *h_s_*(*ν*), it will be useful to first consider the discrete version,

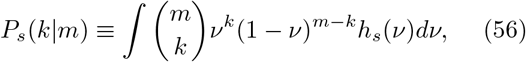

which gives the normalized probability of observing a mutation in exactly *k* individuals in a random sample of size *m*. The continuous version can be recovered by taking the limit of large sample sizes:

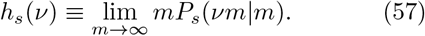

One advantage of switching to *P_s_*(*k*|*m*) is that it can be rewritten as an average over lineages defined at different times in the past:

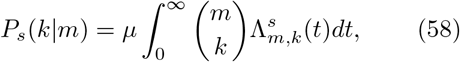

where *μ* is the per-site mutation rate and

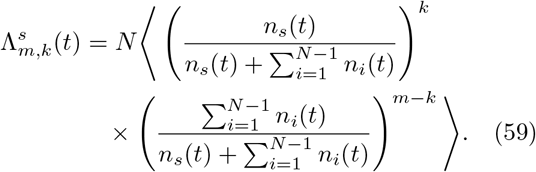

In this equation, *n_s_*(*t*) denotes the present-day size of a lineage founded by a mutation with effect size *s* that occurred *t* generations ago, while the 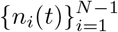 represent the lineages founded by the remaining *N* – 1 individuals that were alive at that time. Eq. (58) can be interpreted as integrating over the possible times a mutation last occurred at a given site. This mutation could have occurred on any of the *N* possible genetic backgrounds in the population. Given that it did occur, it will be observed in *k* individuals in the present with probability 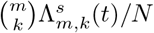.

We note that in the special case of a neutral site (*s* = 0), the quantity 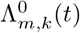 can be interpreted as a *merger probability* within a corresponding coalescent model. That is, 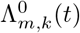 gives the probability that in a sample of size *m*, a particular set of *k* individuals share a common ancestor at *t* generations into the past, and the remaining *m* – *k* individuals do *not* trace back to that same ancestor. Related merger probabilities (defined slightly differently) are considered by Neher and Hallatschek (2013) to show a correspondence between genealogies in the infinitesimal regime and those of the Bolthausen-Sznitman coalescent (BSC). Below, we follow a similar approach to simplify the quantities 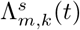 used in our cal-culation of the selected site frequency spectrum. In Appendix E, we use our calculated 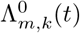 to explicitly demonstrate a (partial) correspondence with the BSC in the MSSM regime, and highlight key ways in which our results differ from those of Neher and Hallatschek (2013).

To simplify Eq. (59), we make the key approximation that each of *n_i_*(*t*) and *n_s_*(*t*) evolve as a collection of independent branching processes described by Eq. (9) (Desai et al., 2013; Fisher, 2013; Neher and Hallatschek, 2013). That is, we assume the fates of lineages are coupled only through the average rate of adaptation, which is shaped by interference as calculated in Section IV. The identity 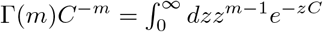 then yields

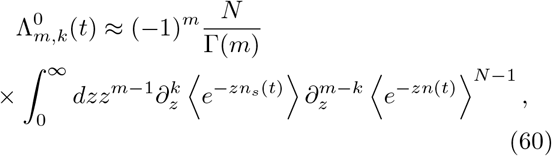

where, since the 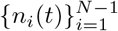 are identically distributed, we have dropped the subscripts *i* in writing *n*(*t*). As we discuss in Appendix A,

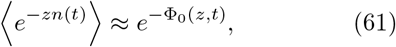

where 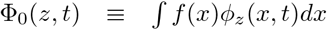, and where *ϕ_z_*(*x*, *t*) satisfies

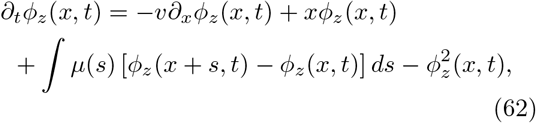

with initial condition *ϕ_z_*(*x*, 0) = *z* > 0; more generally,

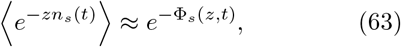

with 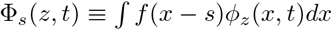.

A detailed analysis of Eq. (62) is conducted by Fisher (2013), which we review and extend in Appendix A. The key results we use here are that

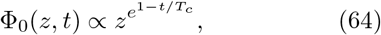

for *b*(*t* – *T_c_*) ≫ 1, over the region of *z* that will turn out to dominate the integral in Eq. (199), and that for 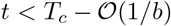, Φ_0_(*z*, *t*) is approximately linear in *z*. In Appendix E, we show, using the properties of *ϕ_z_*(*x*, *t*) discussed in Appendix A along with our result for *f*(*x*) in Eq. (25), that

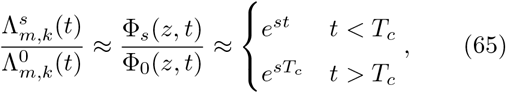

provided that |*s*| ≪ *b*. The relevant site frequency spectra then can then be obtained by substituting these expressions into Eq. (58) and integrating over time.

### 2. The ratio between the selected and neutral SFSs

The simple behavior of Eq. (65) suggests that the *ratios* of neutral and selected site frequency spectra may take on a particularly simple form in certain regimes. For example, if we consider frequencies *ν* = *k/m* that are sufficiently large that the integrals in Eq. (58) are dominated by times greater than *T_c_* (we discuss the minimum frequencies required for this assumption below), then the time-independent behavior of Eq. (65) immediately implies that

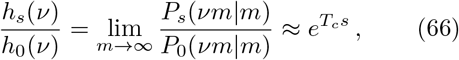

when |*s*| ≪ *b*. Since 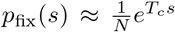, we can also write this as

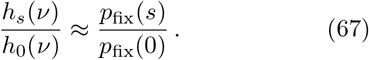

In this range of frequencies, Eq. (67) predicts that neutral and selected site frequency spectra are simply proportional to one another, and that the constant of proportionality is equal to the ratio of their fixation probabilities.

By summing over sites, we can obtain an analogous result for the aggregate site frequency spectra,

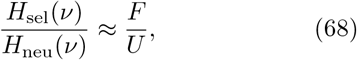

where *F* ≡ *N* ∫ *μ*(*s*)*p*_fix_(*s*)*ds* is the total fixation rate of the selected mutations defined in Eq. (53). Recall that we have defined these aggregate quantities so that a (rough) analogy can be drawn between *H*_neu_(*ν*) and the distribution of *synoynmous* site frequencies, and between *H*_sel_(*ν*) and the distribution of *nonsynonymous* site frequencies. Under this analogy, the right-hand side of Eq. (68) corresponds to *dN/dS*—the ratio of nonsynonymous to synonymous divergence rates (adjusted, as usual, for differences in mutation rates among the two types of mutations) (Yang and Bielawski, 2000).

In Fig. 5, we plot the ratio between *H*_sel_(*ν*) and *H*_neu_(*ν*) as measured in simulations for a subset of the populations considered in Fig. S2. In all cases, the ratio *H*_sel_(*ν*)/*H*_neu_(*ν*) approaches *F/U* as *ν* → 1. This can be understood as a consequence of the fact that mutations already present at the very highest frequencies will drift neutrally to (or away from) fixation. In contrast, as *ν* → 0, *H*_sel_(*ν*)/*H*_neu_(*ν*) → 1 for all cases depicted in Fig. 3 (and thus Eq. (67) breaks down). This is also to be expected: mutations observed at sufficiently low frequencies will have occurred at short enough times into the past that their fates have not yet been substantially impacted by selection.

**FIG. 5.**
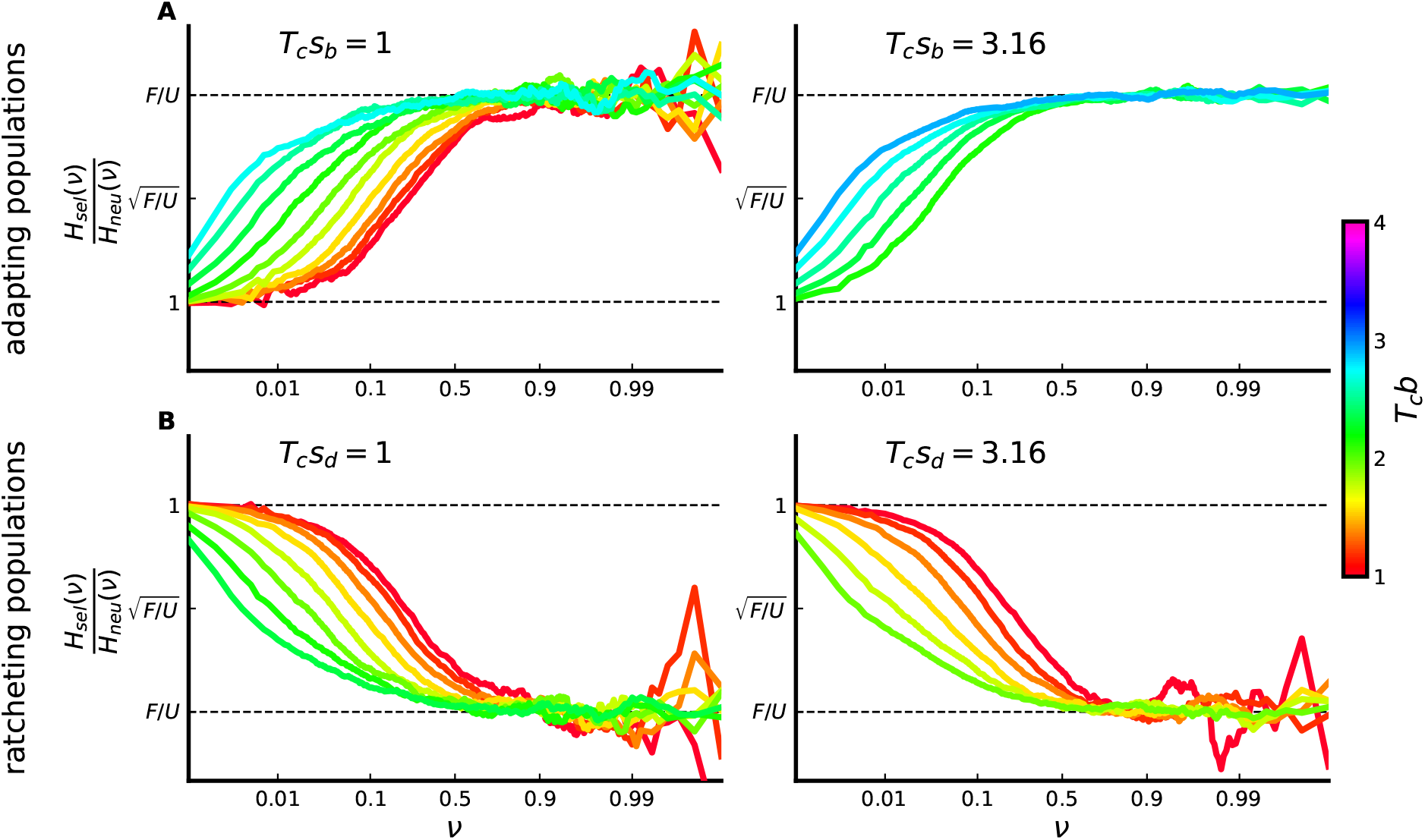
Ratio between selected and neutral SFSs, scaled by the ratio between selected and neutral mutation rates, for adapting populations (A) and ratcheting populations (B) subject to a single-effect DFE with *T_c_s* values denoted in each panel. Ratio curves are plotted on a log scale, and are scaled such that tick marks correspond to the values of 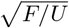 and *F/U* averaged over simulation runs, for each curve (with *F* the fixation rate of selected mutations, either beneficial or deleterious). Each curve corresponds to a simulated population with parameters lying on the constrained grid depicted in Fig. S2; for each panel, the represented *T_c_b* values are linearly spaced.

The behavior we see in these extreme limits is also observed in the independent sites model in Eq. (1) (Hartl et al., 1994; Sawyer and Hartl, 1992). However, these classical results predict that the high frequency limit only applies for extremely large frequencies (1 – *ν* ≪ 1/|*T_c_s*| ≪ 1). In contrast, Fig. 5 shows that *H*_sel_(*ν*)/*H*_neu_(*ν*) ≈ *F/U* over a much broader range of intermediate frequencies in the MSSM regime. This has important consequences for the interpretation of population-genetic data, and in particular for application of the “asymptotic alpha” approach introduced by Messer and Petrov (2013); we comment on this further in the Discussion. Similarly, the independent sites model in Eq. (1) predicts that the deleterious site frequency spectrum will start to differ from its neutral counterpart when *ν* ≳ 1/*T_c_*|*s*|. In contrast, Fig. 5 shows that the frequency scale at which *H*_sel_(*ν*) and *H*_neu_(*ν*) start to differ is not given by 1/*T_c_*|*s*| in the MSSM regime; we discuss related implications for estimates of *T_c_s* in the Discussion.

Importantly, from Fig. 5 it is clear that the shapes of SFS ratio curves—and in particular, the frequency scale on which they transition from 1 to *F/U*—do not depend simply on *T_c_*〈*s*〉 (or, equivalently, on *T_c_*〈*s_f_*〉); a given value of *T_c_*〈*s*〉 is compatible with SFS ratio curves which differ substantially, and the parameter *T_c_b* is equally if not more important in determining the crossover frequency scale. Empirically, we can see that variation of SFS ratio curves (and the crossover frequency scale) is largely mediated by the quantity *Nσ*, another population-level quantity; note that *σ*^2^ = *v* – *U*〈*s*〉 denotes the population-wide fitness variance. To see this across a broad range of parameters, for each observed SFS ratio curve we identified, using spline interpolation, the frequency *ν_c_* at which *H*_sel_(*ν*)/*H*_neu_(*ν*) reaches 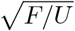 (the half-maximum of *H*_sel_(*ν*)/*H*_neu_(*ν*) in log-space). In Fig. 6, we plot these crossover frequencies *ν_c_* a function of our MSSM approximation predictions for *Nσ*, with points colored by *T_c_*〈*s_f_*〉 values (for adapting populations) or *T_c_*〈*s*〉 values (for ratcheting populations). We can see a simple power law dependence of *ν_c_* on *Nσ*; parameter combinations with similar *Nσ* values have similar *ν_c_* values, even if their values of *T_c_*〈*s*〉 differ substantially. This echoes findings of Good et al. (2014) in the in-finitesimal regime; in that work, a similar collapse is found for the dependence of the neutral heterozygosity. In Fig. S3, we plot the full SFS ratio curves for the same set of parameter combinations, colored by *Nσ* values; there we can see that variation in full SFS ratio curves (in addition to the crossover frequency *ν_c_*) is largely mediated by variation in *Nσ*.

**FIG. 6.**
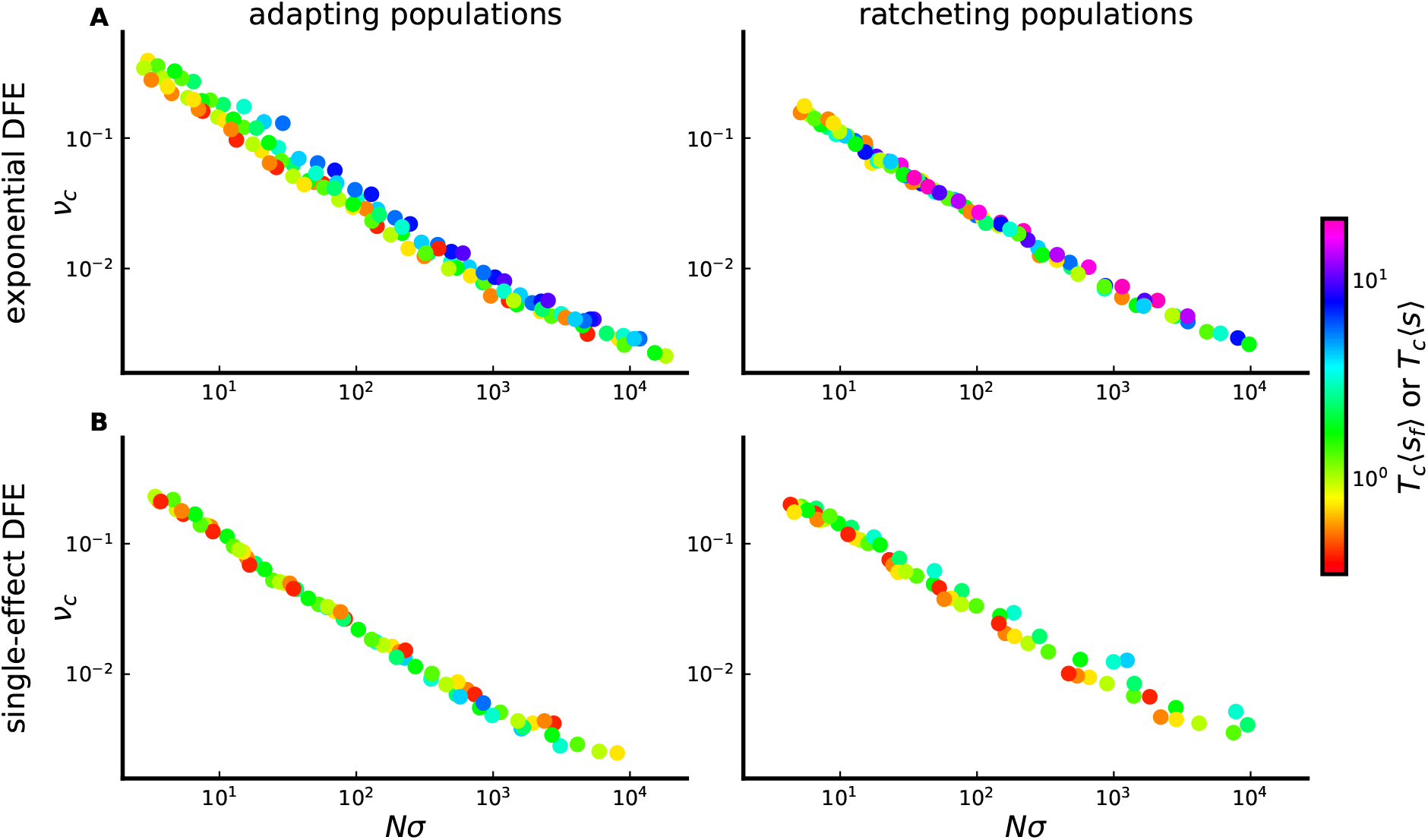
Dependence of the crossover frequency *ν_c_* on *Nσ*. Simulated populations in (A) are subject to an exponential DFE, while simulated populations in (B) are subject to a single-effect DFE. For adapting populations (left) points are colored by their values of *T_c_* 〈*s_f_*〉, while for ratcheting populations (right) points are colored according to their values of *T_c_*(*s*). Values of *Nσ* are computed using the MSSM approximation with *σ*^2^ = *v* – *U*〈*s*〉. For the exponential DFE case, parameters correspond to those on the constrained grid described above and depicted in Fig. S1. For the single-effect DFE case, parameters correspond to the points on a similar constrained grid in the space of *T_c_b* vs. *T_c_s*, depicted in the space of *NU* vs. *Ns* in Fig. S2. For clarity, we have displayed only points corresponding to simulated populations with 〈*s_f_*〉 < 3*b* (such that the MSSM approximation does not break down) and *T_c_* 〈*s_f_*〉 > 1/4 (such that the neutral and selected SFSs differ substantially at high frequencies).

Together, these results suggest that efforts to infer the distribution of scaled effects *T_c_s* using existing approaches (e.g Bustamante et al. (2001); Eyre-Walker et al. (2006)) which “fit” the neutral and selected SFSs may fail when applied to rapidly evolving populations such as those considered here. In particular, these classical results cannot explain the marked dependence of SFS ratio curves on *T_c_b* and/or *Nσ* observed above for fixed values of *T_c_*〈*s*〉.

### 3. The neutral and selected site frequency spectra

We now proceed to compute the neutral and selected SFSs directly. In Appendix E, we use the merger probabilities 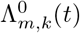 calculated using Eq. (60) and Eq. (64) to simplify Eq. (58), with the result

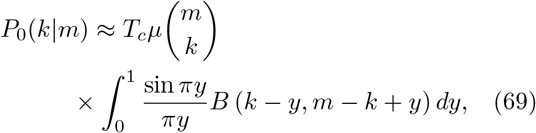

where *B*(*x, y*) is the Beta function satisfying *B*(*x, y*) = Γ(*x*)Γ(*y*)/Γ(*x* + *y*). Up to an overall scale factor, *P*_0_(*k*|*m*) in Eq. (69) matches the SFS corresponding to the BSC, recently calculated by Kersting et al. (2019) directly from the BSC partition structure. Thus, similar to previous work (Desai et al., 2013; Kosheleva and Desai, 2013; Neher and Hallatschek, 2013), Eq. (69) implies a correspondence between genealogies in the MSSM regime and those of the BSC—at least for aspects of genealogies which determine the average SFS at the moderate to high frequencies for which Eq. (69) is valid. We will refer to the large-*m* limit of *mP*_0_(*νm*|*m*) with *P*_0_(*k*|*m*) given by Eq. (69) as 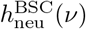, and provide an expression for 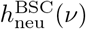 in terms of special functions in Appendix E.

The quantity 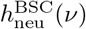 well approximates the actual SFS *H*_neu_(*ν*) only for *ν* such that typical observed mutations have ages *t* satisfying *b*(*t*–*T_c_*) ≫ 1. The time integral yielding Eq. (69) is dominated by times *b*(*t* – *T_c_*) ≫ 1 only when log(1/*ν*) ≪ *T_c_b*. At lower frequencies such that log(1/*ν*) ≫ *T_c_b*, mutations with ages 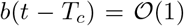 make a dominant contribution to the SFS. Fortunately at these lower frequencies, fluctuations in the size of a focal lineage make a small contribution to the denominator of Eq. (59). As a result, the SFS can be calculated by considering the marginal distribution of a single lineage, which is encoded by the generating function 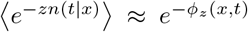. In Appendix A we carry out an analysis of *ϕ_z_*(*x, t*) on fitness scales 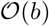 and time scales 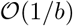, extending the previous analysis of *ϕ_z_*(*x, t*) conducted by Fisher (2013). In Appendix E we show how this generating function can be inverted to obtain an approximation for *h*_neu_(*ν*),

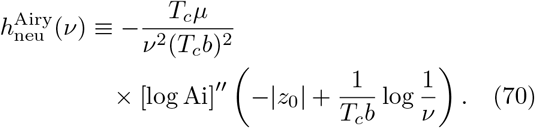

In the limit 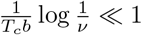,

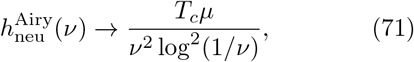

which is precisely the asymptotic behavior of 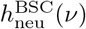 as *ν* → 0; thus the two functions 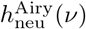 and 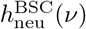 have a smooth crossover at 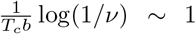. In principle, our predictions 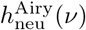 and 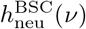 could be connected by asymptotic matching. Notably, 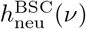 depends on the parameters *N* and *μ*(*s*) only via the overall scale factor *T_c_μ*, which reflects the overall fixation rate of neutral mutations over the timescale *T_c_* of coalescence. In contrast, 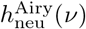 depends separately on the quantities *T_c_μ* (which again sets an overall scale factor) and *T_c_b*. This suggests that for the purposes of distinguishing between (and potentially inferring) evolutionary parameters, an understanding of the low-frequency portion of the SFS is particularly important.

Both 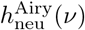 and 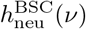, however, neglect the contribution of mutations with ages *t* such that *b*(*t*–*T_c_*) is large and negative. To capture the contribution of mutations with ages *t* < *T_c_*, a deterministic approximation is useful. Under such an approximation, the frequency of an observed mutation—and its age—precisely determine the relative fitness of the ancestral background on which the mutation must have arisen. The contribution to the SFS from ages *t* < *T_c_* can then be obtained by integrating over the times at which a mutation may have occurred, weighted by the corresponding probabilities with which the mutation arose on a background with the respective requisite fitness. A particularly simple approximation, introduced by Neher and Shraiman (2011), can be made by assuming that mutations arise within the “bulk” of the fitness distribution (which is well-described as a Gaussian 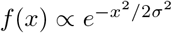, with variance *σ*^2^ = *v* – *U*〈*s*〉) and by approximating *n*(*t*|*x*)—the deterministic lineage size at time *t*, given a founding background fitness *x*—as 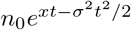 (where *n*_0_ denotes the size of a lineage upon *establishing*, at which point its lineage begins to grow deterministically, under this approximation). This yields a resulting SFS approximated by

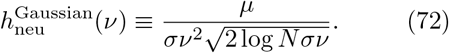

In Appendix E, we generalize the above argument to the regime in which mutations arise outside the “bulk” of the fitness distribution, or at times such that the approximation 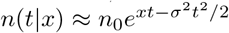 breaks down by the time the mutation is observed. A key simplification arises because of a relation between the deterministic lineage sizes *n*(*t*|*x*) and the Laplace transform 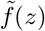 of *f*(*x*). We find that more generally, the deterministic contribution to the SFS is given by

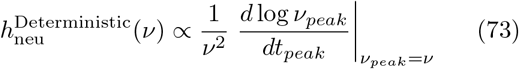

where *ν_peak_*(*t_peak_*) is the frequency at which a lineage peaks in size, given its peak size occurs at time *t_peak_* (which in turn can be expressed as a function of the lineage’s initial relative fitness *x*); the derivative in Eq. (73) is evaluated at *t_peak_* such that *ν_peak_* = *ν*. The applicability of Eq. (73) reflects the fact that the contribution of any particular lineage to the time-averaged SFS is typically dominated by the time the lineage spends near its peak in size; this intuition has been used to calculate the SFS in the presence of purifying selection, for example (Cvijović et al., 2018). As we show in Appendix E, Eq. (73) can easily be approximated when *st_peak_* ≪ 1 for relevant *s* in *μ*(*s*), or when *b*(*t_peak_* – *T_c_*) ≪ 1, and can be evaluated more generally by numerically solving a simple equation for *t_peak_* in terms of *ν_peak_*. In the limit *st_peak_* ≪ 1 (which occurs for sufficiently low *ν*), we find that 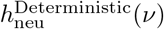 tends to 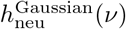 given in Eq. (72), as expected.

Finally, at the very lowest (and highest) frequencies, the SFS is dominated by completely neutral genetic drift. A well-known result is that 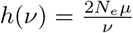 for a neutrally evolving population with effective population size *N_e_* (Crow et al., 1970); at sufficiently low frequencies, mutations contribute in the same way to the SFS, since selection has not yet had sufficient time to alter their fates substantially (Cvijović et al., 2018). As a rough heuristic, we might expect this result to hold with *N_e_*/*N* equated to 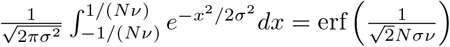, the fraction of individuals in the “bulk” of the fitness distribution which will typically reach a frequency *ν* in the population *before* establishing (i.e., reach a frequency *ν* primarily by genetic drift as opposed to by deterministic forces, contingent on reaching a frequency *ν*). We thus define

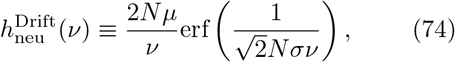

which has a relatively smooth crossover to 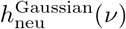 at *ν* ≈ 2/(*Nσ*), roughly the threshold frequency which most mutations founded in the “bulk” of the fitness distribution will not reach before establishing. A similar argument suggests that *h*_neu_(*ν*) → 2*Nμ* as *ν* → 1, although we have not worked out a heuristic for the dependence on *ν* in this limit.

The transitions between these frequency regimes are smoothly varying crossovers. To obtain concrete predictions, it is useful to consider a piecewise approximation,

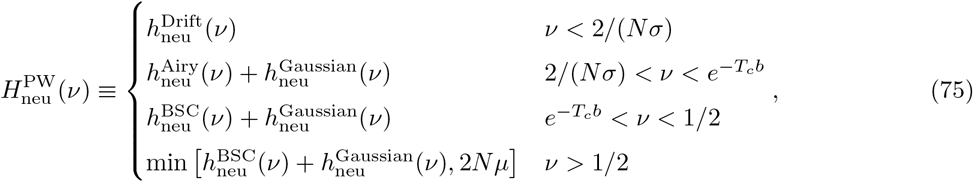

where 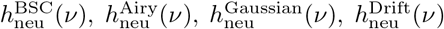 are given above. Note that mutations with ages *t* < *T_c_* (accounted for by 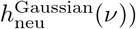 and with ages *t* > *T_c_* or 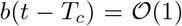 (accounted for by 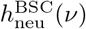 and 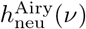, respectively) contribute additively to the SFS, which motivates the inclusion of the sums in Eq. (75). 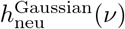 is important primarily for *ν* < *e*^−*T_c_b*^—and over a broad range of log *ν*, both 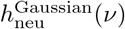 and 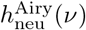 are important—but is retained for *v* > *e*^−*T_c_b*^ to ensure a smooth piecewise curve. In Fig. 7, we compare the predicted SFSs given by Eq. (75) to neutral SFSs observed in simulations. In the same Figure, we compare simulated *selected* SFSs to predicted selected SFSs obtained using a piecewise-defined function 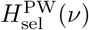. This function is defined completely analogously to 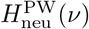 in Eq. (75), with analogous contributions 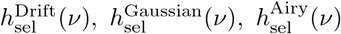, and 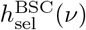 from selected mutations. The only differences are that *μ* is replaced by *μF/U* in 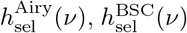, and in the upper limit to 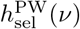 imposed by 2*Nμ*. These replacements are justified because the contributions to 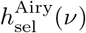 and 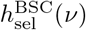 from a mutation with fitness effect *s* both involve overall factors of *e^T_c_s^* (which, integrated over *ρ*(*s*), yield a factor *F/U*). We provide further comparison of 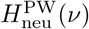 and 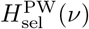 with simulated site frequency spectra in Fig. S4, Fig. S5, Fig. S6 and Fig. S7.

**FIG. 7.**
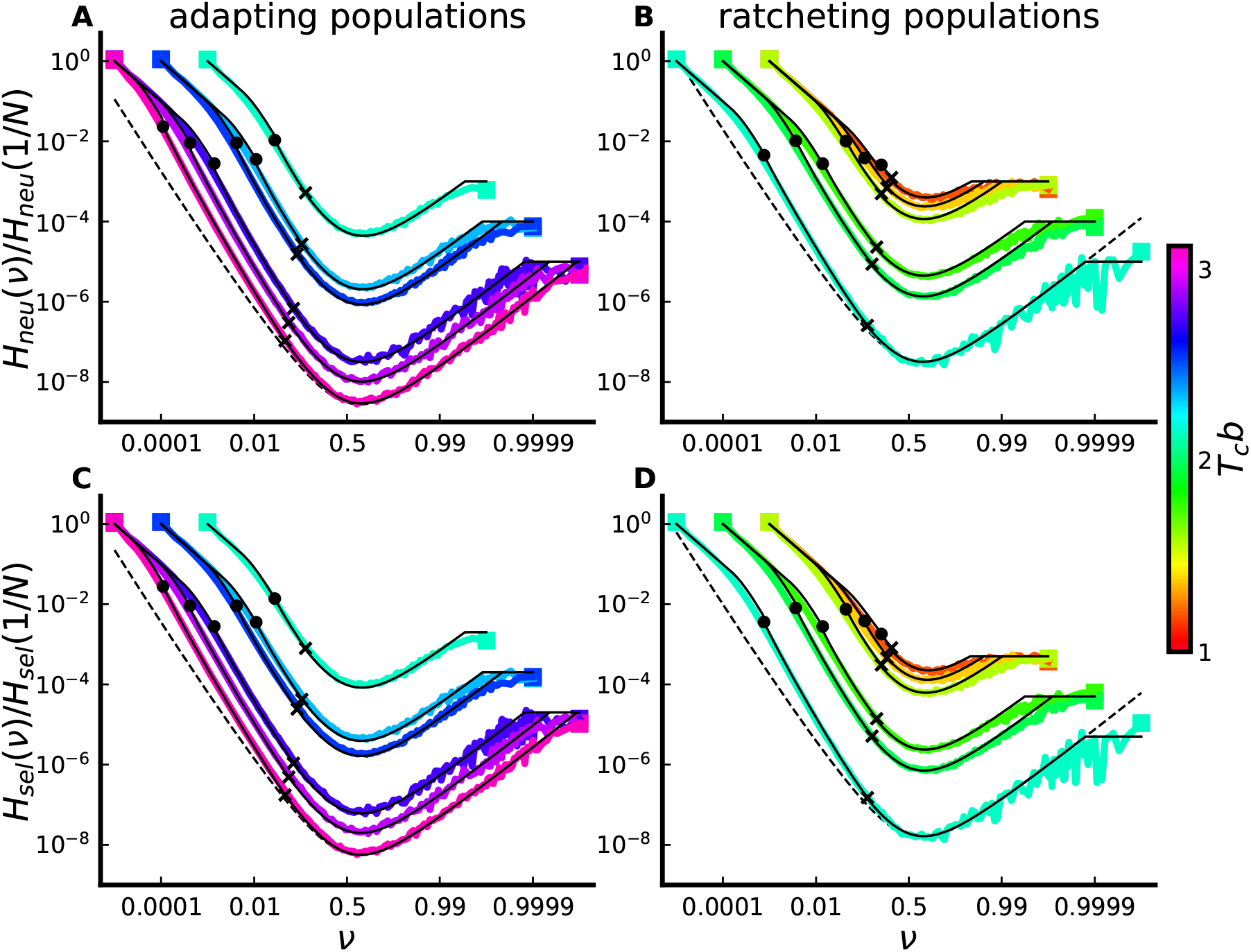
Site frequency spectrum of neutral mutations (A and B) and of selected mutations (C and D). Colored lines denote SFSs observed in simulations, averaged over at least 800 epochs and smoothed using a moving average box kernel smoother. Solid black lines denote the corresponding theory predictions of the piecewise-defined function in Eq. (75) or its generalization to the selected SFS; for each theory curve, the ‘x’ marker denotes the point at which *v* = *e*^−*T_c_b*^, and the circle marker denotes the point at which *ν* = 2/(*Nσ*). Dashed black lines denote the BSC prediction *h*^BSC^(*ν*) for the parameter combination with the largest value of *T_c_b* in each panel (with BSC predictions for other parameter combinations simply shifted by a constant factor). All SFSs and theory curves are normalized by our theoretical prediction for *h*(1/*N*), which is 2*N*^2^*U* for selected mutations and 2*N*^2^*U_n_* for neutral mutations. In all cases populations are driven by an exponential DFE. Simulated parameters are chosen with *T_c_*〈*s_f_*〉 = 1 for adapting populations, and with *T_c_*〈*s*〉 = 1 for ratcheting populations; in both cases *T_c_b* values are linearly spaced and denoted by the color of each curve. Note that each simulated SFS terminates at the frequencies *ν* = 1/*N* and *ν* = 1 – 1/*N* at which we denote simulated SFS values by square markers.

Based on these considerations, *ν_c_* should lie at a frequency such that 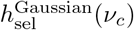 is of comparable magnitude to 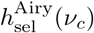, and should obey the approximate bound

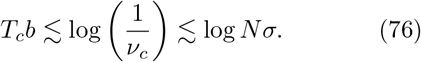

In Fig. S8 we verify that Eq. (76) is satisfied for the populations shown in Fig. 6 for large *Nσ*. We leave a more complete analytical description of the crossover frequency *ν_c_*, as well as the precise frequency-dependence of *h*_sel_(*ν*)/*h*_neu_(*ν*) at the lowest frequencies 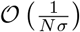, for future work.

### 4. Pairwise heterozygosity and coalescence times

A special case of the neutral site frequency spectrum is the pairwise neutral heterozygosity π_neu_ ≡ *P*_0_(1|2). More generally, we can consider the pairwise heterozygosity *P_s_*(1|2) for a single selected site as well as the aggregate heterozygosity π_sel_ ≡ ∫ *P_s_*(1|2)*ρ*(*s*)*ds*. With these definitions, π_neu_ and π_sel_ can both be computed using Eq. (69) and the SFS ratio given in Eq. (67). These expressions, however, are obtained under the assumption that typical contributing mutations have ages *t* > *T_c_*. In reality, π_neu_ (as well as, rather generally, π_sel_) receives a substantial contribution both from mutations with ages *t* > *T_c_* and from mutations with ages *t* < *T_c_*. In Appendix E we show, using the approximate behavior of Φ*_s_*(*z, t*) for *t* > *T_c_* and for *t* < *T_c_*, that

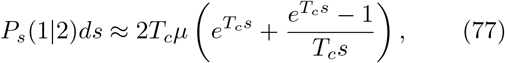

where the second term in Eq. (77) can be considered the contribution to *P_s_* (1|2) from mutations with ages *t* < *T_c_*. This term, which can alternatively be written as 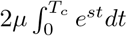, can be interpreted as follows: mutations with ages *t* ≪ *T_c_* are present at expected frequency *e^st^*/*N*, with a negligible fraction of mutations shared among sampled individuals. We note that the *s* → 0 case of Eq. (77) reduces to

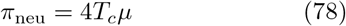

Since π_neu_ is related to the mean time 〈*T*_2_〉 to pairwise *coalescence* according to π_neu_ = 2*U_n_*〈*T*_2_〉, Eq. (78) then implies that

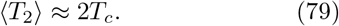

That is, the defined quantity *T_c_* corresponds with (one-half) the average time 〈*T*_2_〉 to pairwise coalescence, motivating our interpretation of *T_c_* as a coalescence timescale. In Appendix E, we obtain the same result by considering the time-dependent pairwise merger probability *Q*_2_(*t*) = Λ_2,2_(*t*). Our calculation in Appendix E also yields the *distribution* of times to pairwise coalescence (and relatedly, the distribution of the pairwise neutral heterozygosity); we find the same exponential distribution of pairwise coalescence times, following an initial delay period of time *T_c_* during which coalescence events are negligible, observed by Neher and Hallatschek (2013) in the infinitesimal regime (with a different overall timescale).

In Fig. 8, we compare our predictions in Eq. (77) for π_neu_ and π_sel_ to averages of these quantities measured in simulations. We find good agreement between simulations and our prediction, provided the MSSM approximation conditions of validity are met, though agreement appears to require larger values of *T_c_b* than for the quantities considered in Fig. 4. We note also that a separate prediction for π can be obtained from the piecewise approximation to the site frequency spectrum in Eq. (75), using the relation 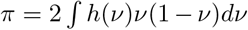. We compare this prediction to the observed values of π_neu_ and π_sel_ in Fig. S9; a similar level of agreement is obtained.

**FIG. 8.**
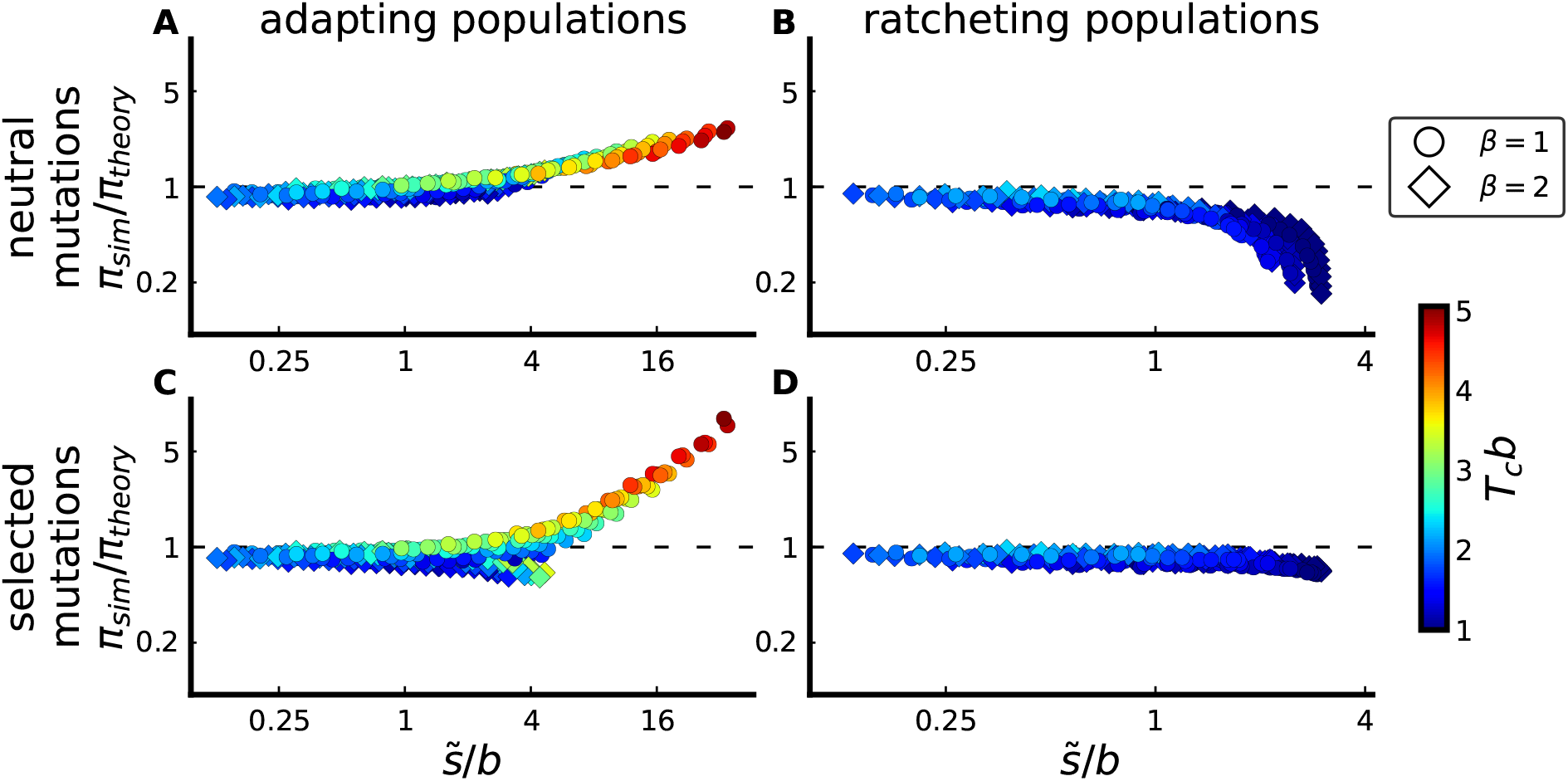
Comparison between simulated and predicted heterozygosity π of neutral mutations (A and B), and between simulated and predicted heterozygosity π of selected mutations (C and D), for adapting populations (A and C) and ratcheting populations (B and D). Parameters are identical to those simulated in Fig. 4.

## VI. KEY FITNESS SCALES AND TIMESCALES: SIMPLE HEURISTICS

We now turn to summarize the different interpretations that can be given to the quantities *T_c_*, *b* and *c* defined above, and provide a heuristic description of the dynamics within the MSSM regime. We begin by noting that the distribution *Nf*(*x*)*w*(*x*) of future common ancestors fitnesses is peaked with width 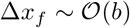 around 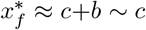. This is our motivation for defining the region 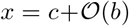 as the *fixation class*, since collectively, this region (of width approximately |*z*_0_|*b*) fixes with probability 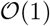, despite its small size relative to the total population. In Appendix C, we show how our approximate solutions *f*(*x*) and *w*(*x*) in Eq. (25) and Eq. (26) can be considered local approximations, valid precisely around *x* = *c*; we obtain a condition of validity for the MSSM regime by ensuring the region of validity of this *local* approximation encompasses the entire fixation class.

### 1. Competition among the fixation class, and the fates of “doomed” lineages

Our condition of validity for the MSSM regime, expressed in Eq. (34), essentially requires that *s* ≪ Δ*x_f_* throughout the region dominating ∫ *s*^2^*ρ_f_*(*s*)*ds*. In this sense, mutational effects can be considered “infinitesimal” in the MSSM regime: the typical fixed fitness effect 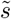 must be much smaller in magnitude than the range of fitness space which typically produces a future common ancestor (i.e., the width of the fixation class). Dynamically, this means that individuals routinely fix despite having fitness disadvantages, compared to the most-fit individuals in the population, of several times (up to 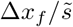 times) the typical fixed fitness effect 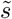. These individuals catch up and fix not by acquiring a single large-effect beneficial mutation, but rather by rapidly acquiring several mutations (and/or by *avoiding* deleterious mutations) so as to “leapfrog” above their more-fit competitors. Less-fit individuals are exponentially less likely to fix—with 1 /*T_c_* the fitness scale on which fixation probabilities vary—but there are also exponentially more of such individuals; over the fitness scale Δ*x_f_* ≈ *b*, the two exponential factors cancel, leaving a relatively broad range of background fitnesses which routinely supply a future common ancestor. Intuitively, it makes sense that this behavior is obtained for high mutation rates (relative to the size of selective effects), but as we have shown, this behavior also arises for sufficiently large population sizes (as long as *ρ*(*s*) falls off faster than exponentially with large, positive *s*). The infinitesimal regime constitutes a special case of the MSSM regime in which mutations are “infinitesimal” in a more narrow sense: in the infinitesimal regime, selection acts in a negligible way on any individual mutation.

In contrast, in the “moderate-speeds” regime, 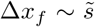: nearly all future common ancestors come from within one “predominant” effect 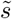 of the nose (Fisher, 2013; Good and Desai, 2014). The lineages of individuals lying more than one multiple of 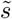 below the nose are essentially “doomed” to eventually go extinct, and the chance with which these individuals leapfrog above their more-fit competitors can be ignored. As a result, in analyzing the “moderate-speeds” regime several simplifications can be made. For instance, in treating a population as a set of discrete fitness classes (separated by predominant-effect mutations), mutations are only important only via their potential to establish a new “lead” fitness class (Desai and Fisher, 2007), and can thus be ignored in fitness classes below the current lead class. Once a new lead class is established, the frequencies of lineages, within the previous lead class, can be treated as “frozen”. Each prior lead class will grow in size—while losing relative fitness at rate *v*—and will eventually comprise a large fraction of the total population, and at that point the frequencies of individual lineages within the class will have changed negligibly. This approximation has been used to simplify calculations of genetic diversity statistics and sojourn/fixation times of mutations (Desai et al., 2013; Kosheleva and Desai, 2013).

In the MSSM regime, it is less useful to model a population as a set of discrete fitness classes, since lineage frequencies are at no point “frozen” within fitness classes. Instead, lineages continue to acquire mutations, and *spread out* in relative fitness as they fall behind the nose. To connect the dynamics of the fixation class with the dynamics of the bulk—which we have done by ensuring *f*(*x*) is appropriately normalized, in enforcing the condition 1/*N* = ∫ *f*(*x*)*w*(*x*)*dx*—accounting for this spread is important. Using a deterministic approximation (carried out in Appendix F) we can gain further intuition on the dynamics of lineages as they fall behind the nose. In the special case of the infinitesimal regime, we review how lineage-wide relative-fitness distributions evolve according to a *reaction-diffusion* equation with diffusion constant 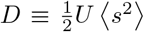, advection rate −*σ*^2^, and local growth rate *x*. The infinitesimal regime has thus long been recognized as a limit of “mutational diffusion” (Tsimring et al., 1996); along the line of descent, fitness follows a biased random walk with diffusion constant *D* (Neher et al., 2014a). More generally in the MSSM regime, we can see that a diffusion approximation may *not* be adequate in describing the trajectories of lineages. In particular if 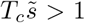 then at later times, further mutations acquired by the lineage will have begun to shape its fitness distribution in a nondiffusive way (i.e. its fitness variance will grow faster than linearly with time).

### 2. Interpretations of the timescale *T_c_*

A key result of the Section V is that *T_c_* ≈ 〈*T*_2_〉/2. This motivates our interpretation of *T_c_* (as defined in Eq. (17)) as a *coalescence* timescale. The timescale *T_c_* can also be interpreted in a few different ways. In Appendix F, we show that under a deterministic approximation, the descendants of the fixation class—initially consisting of 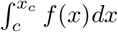 individuals—will collectively sweep through and comprise an 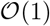 fraction of the population at *T_c_* generations. This motivates us to interpret *T_c_* as a *sweep* timescale. In the “high-speeds regime,” a related interpretation has been given to *T_c_* as the time required for fluctuations near the high-fitness nose of the fitness distribution to substantially affect its bulk dynamics (Fisher, 2013).

Since the (linearized) operators governing the forward-time and backward-time dynamics are adjoints of one another, it is not particularly surprising that the timescale *T_c_* can also be interpreted as a *delay* timescale of coalescence: looking backwards in time, a given pair of individuals is unlikely to coalesce until *T_c_* generations have elapsed. We can understand this correspondence heuristically by viewing the fixation class as an exponentially expanding subpopulation, among the total population, within which all coalescence events must occur (since the fixation class is destined to eventually fix). It is well known that in rapidly *expanding* populations, coalescence events occur primarily at the very beginning of exponential growth (i.e., when the population was small in size) and that genealogical trees are starlike, with long terminal branches (Slatkin and Hudson, 1991); thus, looking backward in time, we expect few coalescence events to occur until near the end of the sweep timescale.

We can also gain intuition on the delay timescale *T_c_* by considering, for a randomly chosen individual, the distribution *A*(*x, t*) of its ancestor’s relative fitness *x* as the time *t* recedes into the past. As we describe in Appendix A,

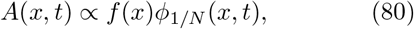

with lim_*t*→∞_ *A*(*x, t*) = *Nf*(*x*)*w*(*x*) corresponding to the distribution of fitnesses of *eventual* common ancestors described above. Importantly, we can see that at *T_c_* generations into the past, the distribution 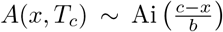 comes to resemble (and, subsequently, rapidly converges to on a timescale 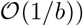) the eventual distribution of ancestor fitnesses 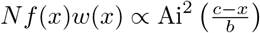. Thus, the ancestors of typical individuals (which start out near “bulk” of the fitness distribution) migrate upward in relative fitness and reach the fixation class on the timescale *T_c_*. After this point, the ancestors of a typical sample of individuals begin to coalesce *within* the fixation class. This matches the interpretation given by Neher and Hallatschek (2013) for the delay timescale in the infinitesimal regime, using a heuristic argument: in that case, ancestors migrate upwards in fitness at initial rate *σ*^2^, slowing down at later times to reach a fitness *σ*^4^/4*D* at the delay timescale *σ*^2^/2*D*.

It is not entirely clear why the delay timescale should match the coalescence timescale of the ensuing BSC process—that is, why coalescence *within* the fixation class requires approximately the same amount of time as that required to *reach* the fixation class. The correspondence between these two timescales appears to be a relatively universal feature of rapidly evolving populations, observed in the moderate-speeds regime (Desai et al., 2013; Kosheleva and Desai, 2013) and the infinitesimal regime (Neher and Hallatschek, 2013). However, this is not generically the case: for instance, in the adapting populations modeled by Brunet et al. (2007) as FKPP waves, the delay timescale is much shorter than the coalescence timescale, even though the genealogies of those populations are described by the BSC. The key difference is that in the model considered by Brunet et al. (2007), the growth rate of a lineage does not depend linearly on its fitness advantage *x*. Instead, all individuals lying above minimum fitness cutoff survive until the next generation, and individuals therefore have a reduced benefit of being much fitter than average.

### 3. Conditional neutrality at long times

The simple exponential dependence 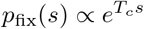 suggests that selection acts in a substantial way to amplify the frequency of a mutation only over the timescale *T_c_*. This in turn suggests a picture of *conditional neutrality* in the fates of mutations at times *t* > *T_c_*. This conditional neutrality can also be seen through our result for the SFS of selected mutations. As we have seen, the SFS of selected mutations with effects between *s* and *s* + *ds* (and ages *t* > *T_c_*) is simply scaled by an overall factor *e^T_c_s^*, relative to the neutral SFS (as well as a scaling factor reflecting the difference in the mutation rates). Thus, we can think of selection acting to amplify the probability with which a mutation is observed only over timescale *T_c_*. At least up to the information contained in the SFS, frequency trajectories of selected mutations at times *t* > *T_c_* are indistinguishable from those of neutral mutations with the same frequency at time *T_c_*. We note that a *stronger* picture of conditional neutrality is seen in the infinitesimal regime. In the infinitesimal regime, *T_c_* ≪ 1/*s* for relevant *s*, so that even during the initial period *t* < *T_c_*, the fate of a typical mutation (or even of a typical *fixed* mutation) is influenced by its selective effect in a negligible way. In the MSSM regime, the selective effect of a mutation can influence its fate in a substantial way, but only during the first *T_c_* generations of the mutation’s lifetime.

At a heuristic level, it makes sense that *T_c_* is the relevant timescale on which selection acts. To see this, we note that looking backward in time, a given pair of individuals which happen to be sampled from the fixation class will coalesce at average time *T_c_* (because they are sampled from the fixation class, they skip the initial delay period). Looking forward in time, then, some individual will have grown to comprise a macroscopic fraction of the fixation class on the same timescale *T_c_*. Note that individuals do not *fix* within the fixation class over the timescale *T_c_*—fixation requires a time 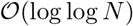 multiples of *T_c_*, given the distributions of times to coalescence of a large sample under the BSC (Berestycki, 2009; Desai et al., 2013). At the timescale *T_c_*, however, lineages originally founded in the fixation class (or more precisely, the portions of these lineages still *in* the fixation class) have spread out enough in fitness so that initial fitness differences of size *s* ≪ *b* have been “forgotten”; selection thus no longer acts to amplify the frequency of a mutation which occurred in that region. Because the fixation class will eventually take over the population, this implies conditional neutrality in the full population on the same timescale.

A similar period of conditional neutrality—after an initial delay—is noted by Kosheleva and Desai (2013) in the “moderate-speeds” regime. In that case, Kosheleva and Desai (2013) note that a mutation observed at a macroscopic frequency *x*_0_—large enough to essentially guarantee the mutation occurred in the “nose” class—will go on to fix with the same probability *x*_0_. This is explained by the fact that lineage frequencies are “frozen” within fitness classes in that case, so that a mutation currently at macroscopic frequency *x*_0_ in the population was once present in the “nose” class at the same frequency *x*_0_, a delay time *T_d_* ago. The fraction of the nose class occupied by any given lineage evolves *neutrally* as a Markov process (in that the *future* fraction of the nose class occupied by any given lineage depends only on the fraction of the nose class it *currently* occupies) and so a period of conditional neutrality emerges after the delay timescale *T_d_*.

In the MSSM regime, the same argument does not quite apply, even upon replacing the “nose” class with the fixation class. The key difference is that the fraction of the fixation class occupied by a given lineage does not evolve as a Markov process: fitness differences among individuals in the fixation class are important in predicting their eventual fates (since *T_c_b* ≫ 1). However, the same argument *does* hold if we instead consider the *effective lead frequency ν_L_* of a lineage, defined as

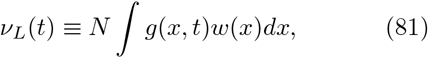

where *g*(*x, t*) is the relative fitness distribution of the lineage (normalized such that its total size at time *t* equals *N* ∫ *g*(*x, t*)*dx*). The generating function 〈*e*^−*θν_L_*(*t*)^〉 is considered by Fisher (2013)—though in that work, *g*(*x, t*) in Eq. (81) is primarily taken as a fitness distribution of an entire *population* whose total size can fluctuate. In Appendix E, we briefly review key features of 〈*e*^−*θν_L_*(*t*)^〉 discussed by Fisher (2013). An implication of these results is that the frequency *n*(*t*)/*N* of a lineage in a population closely mirrors its effective lead frequency *ν_L_*(*t* – *T_c_*) a time *T_c_* into the past (provided the lineage was founded in the fixation class). Furthermore, the effective lead frequency *ν_L_*(*t*) evolves as a neutral Markov process, in that 〈*e*^−*θν_L_*(*t*)^〉 can be obtained from *ν_L_* at any earlier time (i.e., the distribution of *ν_L_* at later times is mediated by its current value, and does not explicitly depend on the full lineage-wide distribution *g*(*x, t*)). Together these results give rise to a type of conditional neutrality at long times similar to that observed by Kosheleva and Desai (2013).

Using the generating function 〈*e*^−*θν_L_*(*t*)^〉, we can obtain the *transition density G_t_*(*ν_L_*(*t*)|*v_L_*(0))—the probability a lineage has effective lead frequency *ν_L_*(*t*) at time *t*, given its effective lead frequency *ν_L_*(0) at time 0—with the result

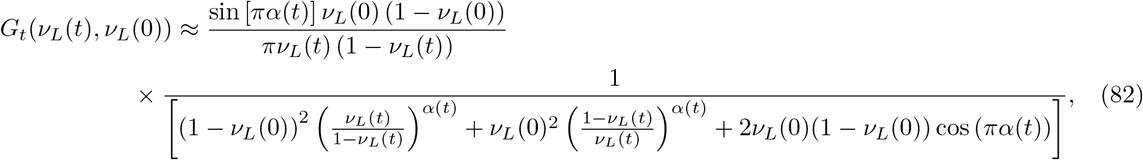

where *α*(*t*) ≡ *e*^−*t/Tc*^. We carry out this calculation in Appendix E. Up to a change in timescale, *G_t_*(*ν_L_*(*t*), *ν_L_*(0)) in Eq. (82) is the same as the transition density for the *actual* lead frequency (i.e the frequency in the “nose” class) found in the “moderate-speeds” regime (Desai et al., 2013; Kosheleva and Desai, 2013). The distributions of sojourn times found by Kosheleva and Desai (2013) using the transition density thus carry over to the MSSM regime as well, after making an overall change in the timescale, and replacing the actual lead frequency with the effective lead frequency. Interestingly, the same transition density is obtained by Hallatschek (2018) in a purely neutral model of a population with a 1/*n*^2^ offspring number distribution from one generation to the next (and which can be considered a forward-time dual to the BSC). As discussed by Hallatschek (2018), this transition density manifests as an apparent *frequency-dependent selection*—a typical bias favoring majority alleles is balanced by rare compensating jumps of low-frequency alleles to maintain overall neutrality. We thus expect to see a similar apparent frequency-dependence among mutational trajectories in the MSSM regime at long times.

Upon incorporating selection into the model described above, Hallatschek (2018) finds the same exponential dependence *p*_fix_(*s*) ∝ *e^T_c_s^* that we have shown is obtained in the MSSM regime. In the model considered by Hallatschek (2018), a mutation with effect *s* is shown to effectively increment the logit frequency log[*ν*/(1 – *ν*)] of a lineage by an amount *s* (i.e., a lineage which just acquired a mutation with effect *s* behaves identically to a neutral lineage whose logit frequency is larger by an amount *s*). In our case, a mutation of effect *s* increments log *ν_L_* by an amount *T_c_s*, provided it occurs within the fixation class (and has effect *s* ≪ *b*). Thus, the model considered by Hallatschek (2018) replicates quite well the long-time dynamics of the MSSM regime, coarse-grained on a timescale of *T_c_* generations. A key difference, however, is that in our model, the timescale *T_c_*—the length of an effective generation—is shaped by selection in a complicated way; in the model considered by Hallatschek (2018), *T_c_* is a fixed input parameter, independent of the action of selection.

The above results emphasize that while the effective frequency a mutation in the fixation class evolves as a neutral Markov process, it does *not* evolve as a neutral Wright-Fisher diffusion process. Thus, even during the conditionally neutral period, we do not expect mutational trajectories to resemble those of a purely neutral Wright-Fisher population—we only expect that lineages have “forgotten” their initial fitness effect. This can be contrasted with a type of quasi-neutrality discussed by Cvijović et al. (2018) in a model of strong purifying selection (|*T_c_s*| ≫ 1). In that model, the fitness advantage of individuals in the “lead” class is balanced, on average, by the rate of mutations *out* of the “lead” class. As a result, fluctuations in lineage frequencies within the “lead” class closely resemble those in a purely neutral Wright-Fisher population. The same fluctuations, smoothed on a certain timescale, are then mirrored by the fluctuations in overall mutational frequencies, after a delay period. Relatedly, the SFS for this model has a “quasi-neutral” region scaling as 1/*ν* for moderate frequencies *ν*, and differs from the SFS of the BSC in important ways (Cvijović et al., 2018); in contrast, in the MSSM regime deviations from the SFS of the BSC are only observed at very low frequencies with log(1/*ν*) > *T_c_b* (and to some extent, at the very highest frequencies). This difference highlights the fact that the presence of multiple selected mutations in a population at once—which is the case for the model considered by Cvijović et al. (2018)—is not sufficient to give rise to the seemingly universal correspondence to the BSC we have described above. Rather, the struggle among fit lineages to increase fitness through new mutations, and the jackpot dynamics this gives rise to, appear to be important features in giving rise to this correspondence.

## VII. DISCUSSION

As we have seen, evolutionary dynamics within asexual genomes can be complex, even within the simplest models that include only the effects of mutations, natural selection, and genetic drift. The central difficulty is that when the mutation rate is sufficiently high and the population size is sufficiently large, multiple selected mutations often segregate simultaneously and their dynamics are not independent. We refer to this scenario as *rapid evolution*, because evolution is not primarily limited by the waiting time for new mutations to arise. Instead, numerous mutations arise in a variety of linked combinations, and selection can only act on these combinations as a whole. The resulting complex dynamics of clonal interference and hitchhiking can limit the efficiency of natural selection, and dramatically alter evolutionary dynamics and population genetics.

We note that rapid evolution does not necessarily have to involve adaptation. The key components of rapid evolution are simply that numerous selected mutations segregate simultaneously within a linkage block—such that the population maintains substantial variation in fitness—and that the population moves through fitness space over time as selected mutations arise and fix. It can therefore involve both beneficial and deleterious mutations, and in particular can result when the accumulation of beneficial and deleterious mutations balances so that *v* = 0, and the population on average neither increases or decreases in fitness (Goyal et al., 2012). It can even occur in scenarios where only deleterious mutations are possible (and hence the rate of change in mean fitness, *v*, will be negative), as long as deleterious mutations routinely fix (Neher and Hallatschek, 2013).

In the past two decades, many authors have analyzed evolutionary dynamics in rapidly evolving populations using *traveling wave* models. However, previous work on these models has largely been focused on two limiting cases: the case in which selection is *strong* on single mutations (the “moderate-speeds” regime and “high-speeds” regime), and the case in which selection is *weak* on single mutations (the infinitesimal limit). These two limits correspond to the cases where 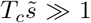 and 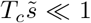 respectively, where *T_c_* is the coalescence timescale and 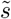 is the typical fitness effect of a fixed mutation. In other words, this previous work has assumed a strong separation of scales between the timescale *T_c_* on which common ancestry is determined, and the timescale 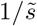 on which selection can act on a typical fixed mutational effect 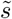.

In reality, however, any population is likely to experience mutations with a wide range of selective effects, including many in the intermediate regime between these two extremes. Our lack of understanding of the dynamics of those mutations for which *T_c_s* ~ 1 thus represents an important gap. This is particularly problematic because it is natural to expect that “nearly neutral” mutations with effects on the order of the inverse coalescence timescale (i.e. for which *T_c_s* ~ 1) may have the largest impact on patterns of genetic diversity (they are strong enough that their effects are felt, but not so strong that they immediately sweep or are purged) (Akashi et al., 2012; Ohta, 1973).

This expectation is broadly consistent with numerous empirical studies that have attempted to infer distributions of selection coefficients in natural populations based on population genetic data (Eyre-Walker and Keightley, 2007). These studies typically do not infer selection strengths directly, but rather the product *T_c_s* (Sawyer and Hartl, 1992; Sawyer et al., 2003) (this is often referred to as *N_e_s* under the assumption that *T_c_* corresponds to an effective population size, but we have avoided this terminology since the dynamics differ in many important ways from those of a neutral population with an appropriately sized effective population size (Neher, 2013)). Many of these studies find mutations with *T_c_s* ~ 1 are quite prevalent and comprise a large proportion of fixed mutations. This includes mutations involved in viral and mitochondrial DNA (Nielsen and Yang, 2003), amino-acid substitutions in *Drosophila* (Sawyer et al., 2007), synonymous mutations affecting codon usage in *E. coli* (Hartl et al., 1994) and *Drosophila* (Akashi, 1995; Machado et al., 2020; Zeng and Charlesworth, 2009), as well as among mutations occurring within animal mitochondria (Nachman, 1998).

It has remained unclear whether these results actually imply that typical selective coefficients are of order 1/*T_c_*, or whether emergent aspects of the evolutionary dynamics tend to generate patterns of variation that are most sensitive to the subset of mutations in this regime. In addition, because these inference approaches typically assume free recombination (Bustamante et al., 2001; Sawyer and Hartl, 1992; Sawyer et al., 2003), it is unclear whether interference may confound these results (see e.g. McVean and Charlesworth (2000)).

These considerations highlight the importance of understanding the evolutionary dynamics and population genetics of rapidly evolving populations in cases where a relatively broad range of selective effects is relevant, including effects *s* with *T_c_s* ~ 1. In this work, we make progress in this direction by extending existing methods to apply within a broader regime of the population-genetic parameter space, which we refer to as a *moderate selection, strong mutation* (MSSM) regime. In the MSSM regime, no assumption is made about the magnitude of 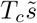; instead, we require that *T_c_*Δ*x_f_* ≫ 1 and 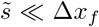, where Δ*x_f_* is the standard variation of fitness advantages among individuals which eventually take over the population. The first of these conditions is also required within the infinitesimal regime, and implies that selection is strong among *haplotypes* competing for fixation. The second of these conditions differs from the condition 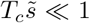 required within the in-finitesimal regime. Instead of requiring that typical fixed mutational effects are weak (and fix essentially neutrally), within the MSSM regime we require only that typical fixed mutational effects are weak compared to the scale of fitness variation among potential future common ancestors.

Our results are not a complete solution to the problem, since our analysis does make other assumptions, and in particular it does not apply to populations in the “moderate-speeds” regime. However, in combination with earlier work, our analysis helps to provide a more complete picture of how mutations with effects on a wide range of scales shape the evolutionary dynamics of rapidly evolving populations. By explicitly demonstrating an approximate correspondence between genealogies in the MSSM regime and those of the BSC, our analysis further supports previous claims that certain aspects of the evolutionary dynamics of rapidly evolving populations are relatively universal (Neher and Hallatschek, 2013). Given this apparent universality, we expect that many of our qualitative conclusions (e.g. for selected and neutral site frequency spectra) may apply in rapidly evolving populations more generally. With this in mind, the MSSM regime is a particularly useful regime of the parameter space to study: as in the infinitesimal regime, the evolutionary dynamics are relatively tractable, even for a full distribution of fitness effects, but unlike in the infinitesimal regime, the dynamics of neutral mutations and of selected mutations differ in a substantial way.

In particular, in this work we have computed the selected and neutral site frequency spectra in the MSSM regime, from which predictions for *dN/dS*, π_*N*_/π_*S*_ and related statistics can readily be obtained. While these quantities are used extensively to infer the strength and presence of selection in natural populations, our analytical understanding of these quantities has previously remained limited when linked selection is widespread. These quantities are considered by Kosheleva and Desai (2013) in the “moderate-speeds” regime, although that analysis is limited to the case in which mutations each confer a single strongly beneficial effect *s_b_* (with *T_c_s_b_* ≫ 1). Our present results allow us to calculate these polymorphism and divergence statistics for full distributions of beneficial or deleterious fitness effects, including those with mutations near 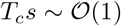. This revealed that the MSSM regime can produce dramatic departures from existing intuition based on independently evolving sites.

At sufficiently low frequencies, the ratio between nonsynonymous and synonymous site frequency spectra, *h_N_*(*ν*)/*h_S_*(*ν*), approaches the ratio of the underlying mutation rates, as expected under neutrality. For deleterious mutations, the frequency scale at which the shape of *h_N_*(*ν*) starts to deviate from *h_S_*(*ν*) is often used as a rough estimate of 1/|*T_c_s*|. Here, we have found that in the MSSM regime, this transition does not occur for frequencies *ν* ~ 1/|*T_c_s*|, but instead depends strongly on population-level quantities such as *T_c_*Δ*x_f_* or *Nσ*. This suggests that naive estimates of *T_c_s* based on deviations of synonymous from nonsynonymous site frequency spectra may severely overestimate the underlying selection strengths.

We have also shown that over a broad range of higher frequencies, nonsynonymous and synonymous frequency spectra are again related by a constant factor, equal to the ratio of the fixation rates of the two types of mutations (*F_N_* and *F_S_*). As a consequence, the frequency-resolved McDonald-Kreitman statistic,

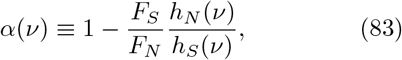

approaches 0 for moderate to large frequencies *ν*, even for large |*T_c_s*|, and regardless of the rate of adaptation *v*. This quantity has previously been used to estimate the fraction of fixed mutations which were strongly beneficial, based on the value to which *α*(*ν*) asymptotes at large frequencies (Messer and Petrov, 2013). Our present results suggest that in the MSSM regime, this “asymptotic alpha” approach may severely underestimate the actual fraction of adaptive substitutions, and might even give the impression that a population is evolving nearly neutrally.

Throughout our analysis, we have assumed that the DFE *μ*(*s*) does not change as the population evolves. This will be true provided that there is no *microscopic epistasis* between individual mutations, such that the fitness effect of one mutation depends on the presence or absence of the other. However, our analysis can also apply even in the presence of extensive microscopic epistasis between individual mutations, provided that the overall DFE *μ*(*s*) does not vary across genotypes – that is, provided there is no *macroscopic epistasis* (Good and Desai, 2015).

Recent experimental work suggests that both microscopic and macroscopic epistasis are widespread, at least in the evolution of laboratory microbial populations (Jerison and Desai, 2015). For example, several recent studies have found general patterns of diminishing returns epistasis, where fitness effects of beneficial mutations systematically decline in more-fit genetic backgrounds (Kryazhimskiy et al., 2014). Recent work has also shown an analogous pattern where the fitness costs of deleterious mutations become more severe in more-fit genetic backgrounds (Johnson et al., 2019). These patterns of macroscopic epistasis suggest that *μ*(*s*)—and particularly *ρ*(*s*)—may change substantially and systematically as a population evolves.

Our analysis, like most previous work on traveling wave models, does not directly address these effects of changing *μ*(*s*). However, provided that *μ*(*s*) changes slowly compared to timescale *T_c_* on which mutations sweep through the population, we expect our analysis to provide an accurate description of the evolutionary dynamics at any given moment (given the appropriate *μ*(*s*) at that moment). Thus we can potentially model the effects of diminishing returns and increasing cost epistasis (or any other systematic variation in *μ*(*s*)), provided only that these changes in *μ*(*s*) are sufficiently slow. Analogously, our approach does not explicitly consider situations where changes in environmental condition lead to temporal fluctuations in the DFE (e.g., a time-varying fitness *seascape* (Agarwala and Fisher, 2019; Schiffels et al., 2011)). However, provided that these temporal fluctuations in *μ*(*s*) are sufficiently slow, our approach will appropriately describe the evolutionary dynamics at any given moment. Of course, if *μ*(*s*) changes rapidly, either due to dramatic epistatic effects or environmental shifts, a transient period may occur where the traveling wave has not reached its steadystate shape. These transient dynamics are understood only coarsely at a theoretical level (Fisher, 2013), and we have not analyzed their effects here. If these shifts are sufficiently common that the transients play a significant role in the overall evolutionary dynamics, the traveling-wave approach will break down.

## ACKNOWLEDGMENTS

We thank Ivana Cvijović and Jiseon Min for useful discussions and comments on the manuscript. MJM acknowledges support from the NSF-Simons Center for Mathematical and Statistical Analysis of Biology at Harvard University (NSF grant DMS-1764269) and the Harvard FAS Quantitative Biology Initiative. This work was also supported in part by the NIH (R01-GM104239), the Simons Foundation (Grant 376196), and the NSF (PHY-1914916). Computational work was performed on the Cannon cluster supported by the Research Computing Group at Harvard University.

## Appendix A CHARACTERISTIC EQUATION FOR *ϕ*(*x, τ*)

In this Appendix, we review much of the key formalism used by Good and Desai (2014), Neher and Hallatschek (2013) and Fisher (2013) to analyze traveling wave models of asexual evolution. In particular, we derive equations for the density *f*(*x*) of individuals with relative fitness *x* and the fixation probability *w*(*x*) of an individual with relative fitness *x*. We then compute the generating function 〈*e*^−*zn*(*t*|*x*)^〉 for the stochastic size *n*(*t*|*x*) of a lineage *t* generations after its foundation, given the relative fitness *x* at which it was founded. In Appendix E, we use properties of 〈*e*^−*zn*(*t*|*x*)^〉—as well as 〈*e*^−*zn*(*t*)^〉, where *n*(*t*) does not condition on the relative fitness *x* at which a lineage is founded—discussed here to compute statistics of genetic diversity such as the distribution of times to pairwise coalescence and the site frequency spectrum.

To begin, following the approach taken by Good and Desai (2014), we consider a lineage with initial absolute fitness distribution *g*(*X*, 0) at time *t* = 0. Both the size and fitness distribution of the lineage may vary with time; we denote by *g*(*X, t*) the time-dependent absolute fitness distribution of the lineage, normalized such that the lineage consists of *n*(*t*) ≡ ∫ *g*(*X, t*)*dX* individuals at time *t*. The time-dependent trajectory of the lineage is coupled to the trajectory of the rest of the population, whose time-dependent absolute fitness distribution we denote by *f*(*X, t*). These two quantities obey the following equations:

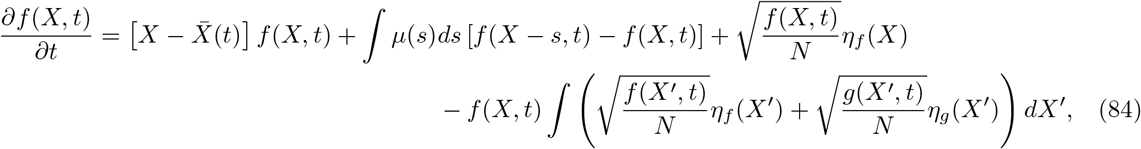

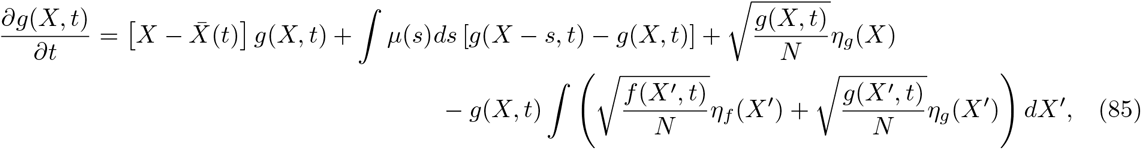

with the stochastic nature of births and deaths captured by the last two terms in Eq. (84) and Eq. (85). Here *η_g_* denotes a Gaussian random variable satisfying 〈*η_g_*(*X*)〉 = 0 and 〈*η_g_*(*X*)*η_g_*(*X*′)〉 = 2*δ*(*X* – *X*′) (and likewise for *η_f_*) (Gardiner et al., 1985). By construction, these noise terms ensure that ∫[*f*(*X, t*) + *g*(*X, t*)]*dX* is time-independent, such that a constant population size is maintained. The quantity 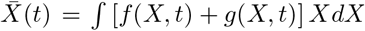 denotes the mean absolute fitness of the population at time *t*, such that 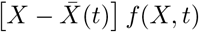 denotes the rate at which *f*(*X, t*) changes due to selection alone. Similarly, ∫*μ*(*s*)*ds* [*f*(*X* – *s*, *t*) – *f*(*X, t*)] denotes the rate at which *f*(*X, t*) changes due to mutation alone. We note that in Eq. (84) and Eq. (85), *f*(*X, t*) and *g*(*X, t*) are treated on a completely equal footing (in the sense that no assumption is made that a ∫ *f*(*X, t*)*dX* ≫ ∫ *g*(*X, t*)*dX* or vice versa). In what follows, however, we focus on the case ∫ *g*(*X, t*)*dX* ≪ 1 (and thus ∫ *f*(*X, t*)*dX* ≈ 1), motivated by our key assumption that the fates of relevant lineages are determined while rare.

Throughout, we follow previous authors in making a “mean-field” approximation to Eq. (84) (Fisher, 2013; Good and Desai, 2014; Good et al., 2012; Neher et al., 2010). That is, we assume that at steady state, fluctuations shape the distribution *f*(*X, t*) only via their effects on the mean fitness 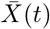 of the population, which can be approximated as increasing at a constant rate *v*. The population-wide fitness distribution *f*(*X, t*) can then be described as a fixed-profile traveling wave, with *f*(*X, t*) = *f*(*X* – *vt*, 0) ≡ *f*(*x*). The quantity *f*(*x*) can be interpreted as the distribution of *relative* fitnesses *x* in the population, with *x* ≡ *X* – *vt*. Without loss of generality, we take 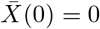; we further assume that *f*(*X*, *t*) has already attained a steadystate traveling wave profile at *t* = 0, potentially following an initial transient period. Given the rate *v*, *f*(*x*) obeys

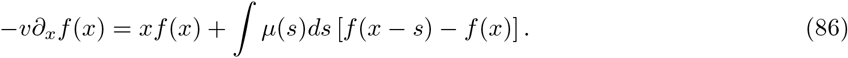

As discussed in the main text, the solution to Eq. (86) attains negative values at large *x* (Fisher, 2013). To alleviate this pathological behavior, we follow previous authors in implementing a cutoff in *f*(*x*), taking *f*(*x*) ≈ 0 for *x* > *x_c_*.

The rate of change of mean fitness, *v*, can be obtained by considering the probabilities with which new mutations eventually fix. This in turn can be done using a stochastic treatment of newly seeded lineages, which are described by Eq. (85). We make the key assumption that the fates of relevant lineages are determined while rare, so that in treating Eq. (85), we can focus on the case ∫ *g*(*X, t*)*dX* ≪ 1. Following the approach taken in Good and Desai (2014), this implies that the last term in Eq. (85) can be neglected, so that *g*(*X, t*) approximately satisfies

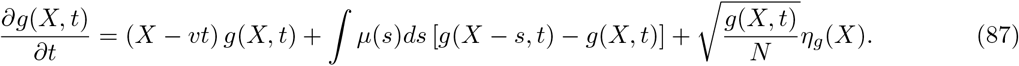

The *relative* fitness distribution *g*(*x, t*) of a lineage approximately then satisfies

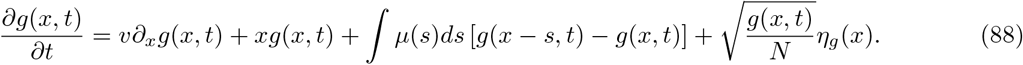

Here, by *Ng*(*x, t*) we denote the number density at time *t* of individuals in the lineage with relative fitness *x* (or, equivalently, with absolute fitness *vt* + *x*). Given Eq. (87) or Eq. (88), fluctuations in lineage-wide fitness distributions can be examined, under the assumption that the background population advances in fitness at a constant rate *v*. In reality, fluctuations in lineage sizes are coupled to fluctuations in the rate of adaptation; this is particularly important at later times, when a lineage may comprise a macroscopic fraction of the total population with substantial probability. In assuming a constant rate *v* at which the population adapts, we neglect this coupling between the rate of adaptation and lineage sizes. Importantly, however, we do ensure that on average, fluctuations in lineage sizes (either to fixation or to extinction) give rise to a self-consistent rate of adaptation *v*.

To examine fluctuations in *g*(*x, t*), we consider the generating functional *H*[*ϕ*(*x*), *t*], defined as

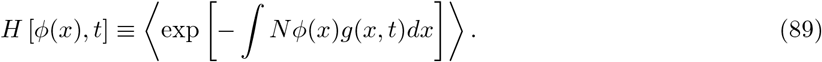

As discussed by Good and Desai (2013), we interpret (88) in the Itô sense, such that

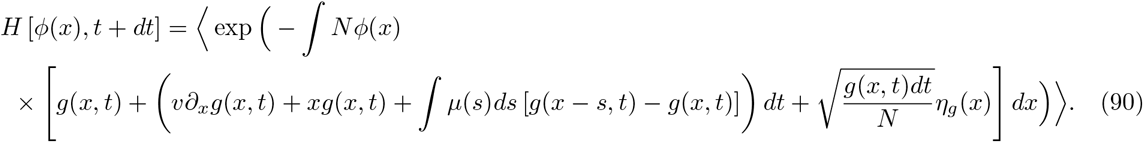

Retaining only terms 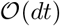 and larger, Eq. (90) simplifies to

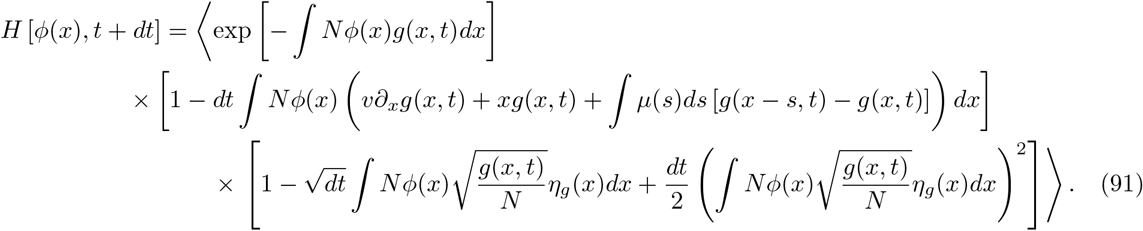

Eq. (91) can be expressed as a differential equation for *H*, which, exploiting the independence of *g* and *η_g_*, takes on the form

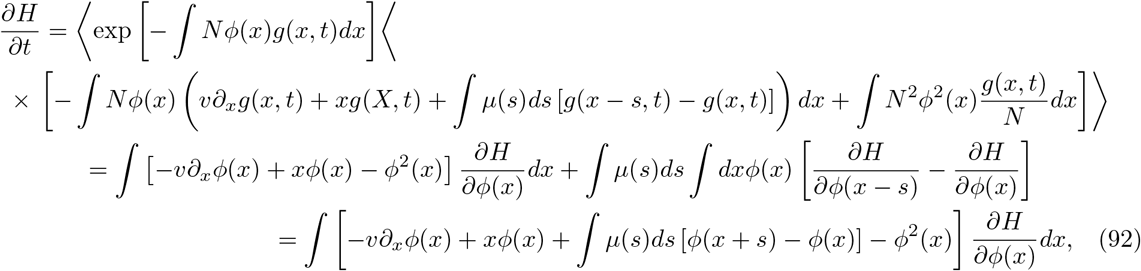

where the second equality is obtained using integration by parts, along with the fact that 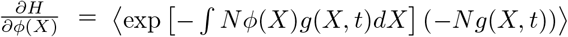.

This enables us to identify characteristic curves *ϕ*(*x, τ*) satisfying

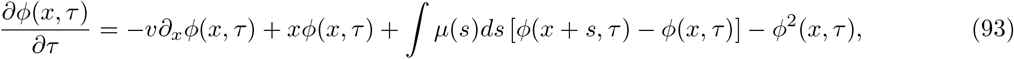

with the property that *H*[*ϕ*(*x*), *t*] = *H*[*ϕ*(*x*, *τ*), *t* – *τ*] for any *τ* > 0. In particular,

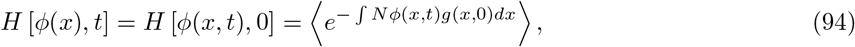

where the averaging in Eq. (94) is only over possible initial conditions—that is, possible instantiations of *g*(*x*, 0)—and not over the stochasticity of the evolutionary process. The problem of calculating *H*[*ϕ*(*x*), *t*] is therefore reduced to solving Eq. (93) for *ϕ*(*x*, *τ*) given the initial condition *ϕ*(*x*, 0) = *ϕ*(*x*).

### A.1 Interpretation of *ϕ_z_*(*x, t*)

In the remaining analysis, we will denote by *ϕ_z_*(*x*, *t*) the solution *ϕ*(*x*, *t*) to Eq. (93) that follows from the initial condition *ϕ*(*x*, 0) = *z*. Then

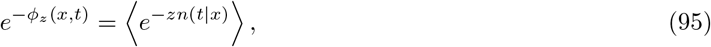

where *n*(*t*|*x*) is the (stochastic) size of a lineage founded at initial relative fitness *x*, after *t* generations (i.e., we can obtain a generating function for *n*(*t*|*x*) by solving for *ϕ_z_*(*x*, *t*)). We note that in general, the initial condition *ϕ*(*x*, 0) need not be independent of *x*. In Appendix E, we will briefly discuss and apply results of Fisher (2013) for the case *ϕ*(*x*, 0) = *θw*(*x*), which yields a generating function for a lineage’s effective lead frequency *ν_L_*(*t*) = ∫ *g*(*x*, *t*)*w*(*x*)*dx*.

A solution for the long-time limit of *ϕ_z_*(*x*, *t*) is particularly useful, since for *z* > 0, lim_*t*→∞_ *ϕ_z_*(*x*, *t*) converges to *w*(*x*) (Fisher, 2013; Good and Desai, 2014). To see this, we note that the above analysis of *H* (and thus of *ϕ_z_*(*x*, *t*)) is done under the branching process approximation, which assumes that lineages are small enough such that population-size constraints are unimportant. Consequently, fixation corresponds to the case in which *n*(*t*|*x*) tends to infinity (rather than the total population size *N*). At sufficiently long times, *e*^−*zn*(*t*|*x*)^ can only approach 1 (provided that the lineage goes extinct) or 0 (provided that the lineage fixes). That is,

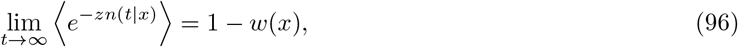

where *w*(*x*) is defined as the fixation probability of an individual with relative fitness *x*. Given a solution *ϕ_z_*(*x*, *t*) to Equation (93), it follows from (94) that

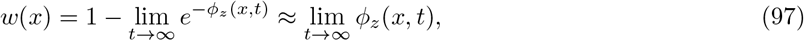

where the last approximation is employed under the assumption that *w*(*x*) ≪ 1 (which holds for *x* ≪ 1, as has been previously assumed in our analysis). Note, then, that *w*(*x*) is the unique long-time limit of *ϕ_z_*(*x*, *t*); at sufficiently long times, *ϕ_z_*(*x*, *t*) does not depend on the initial condition *ϕ*(*x*, 0) = *z*, as long as *z* > 0. The fixation probability *w*(*x*) is therefore the solution *ϕ*(*x*, *t*) to Eq. (93) with the time-derivative term neglected, and satisfies

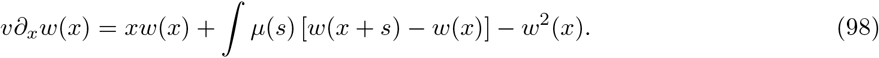

This equation can also be obtained using standard branching process techniques (see e.g. Good et al. (2012); Neher et al. (2010)). In the main text, we discuss approximate solutions to Eq. (98) and Eq. (86) within the MSSM regime.

As discussed by Neher et al. (2014a) in the infinitesimal regime, *ϕ_z_*(*x*, *t*) can additionally be interpreted as a lineage *sampling probability*. In particular, upon sampling *Nz* individuals at time *t*, *ϕ_z_*(*x*, *t*) gives the probability at least one of the sampled individuals is a descendant of a certain individual with relative fitness *x* at time 0. To see this, note that

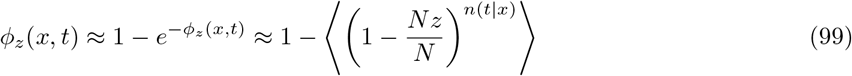

where the right-hand side is precisely the above-described lineage sampling probability (with the expectation value on the right-hand side taken over *n*(*t*|*x*)). The quantity *ϕ_z_*(*x*, *t*) with *z* = 1/*N* is particularly useful to consider, since this gives the probability that a single sampled individual is the descendant of a given individual with relative fitness *x* a time *t* into the past. From a simple application of Bayes’ theorem, we can obtain the probability distribution *A*(*x*, *t*) for the relative fitness *x* of the *ancestor* of a randomly chosen individual (at *t* generations into the past):

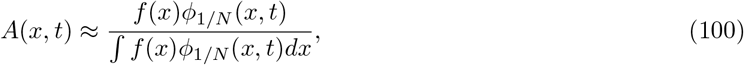

where *f*(*x*) can be interpreted as a prior probability distribution for the ancestor’s relative fitness. Note that

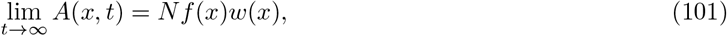

so that, as recognized by Hallatschek (2011), *Nf*(*x*)*w*(*x*) can be considered a distribution of relative fitnesses of future common ancestors of the population. In the main text, we discuss how Eq. (100), together with our solutions for *f*(*x*) and *ϕ_z_*(*x*, *t*) in the MSSM regime, can be used to address the timescale on which individuals in the “bulk” of the fitness distribution descend from its high-fitness “nose”.

We will also consider the quantity Φ_0_(*z, t*) ≡ ∫ *f*(*x*)*ϕ_z_*(*x*, *t*)*dx*. By calculating Φ_0_(*z, t*), we can obtain the generating function for the size *n*(*t*) of a lineage founded by a randomly chosen individual in the population, after *t* generations. That is,

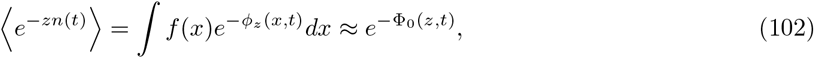

where in the second equality, we used the fact that *ϕ_z_*(*x*, *t*) ≪ 1.

### A.2 Review of properties of *ϕ_z_*(*x*, *t*)

We now review some key properties of the family of time-dependent solutions *ϕ_z_*(*x*, *t*). These properties are discussed by Fisher (2013) in the context of populations subject only to beneficial mutations, in either the moderate-speeds 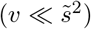 or high-speeds 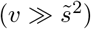 regimes, although the key ideas hold more generally, including throughout the MSSM regime. In the following Subsection, we further analyze *ϕ_z_*(*x*, *t*), explicitly demonstrating that these properties hold within the MSSM regime and extending the analysis conducted by Fisher (2013) to intermediate times that are particularly relevant to our calculation of the site frequency spectrum.

For short times (*t* < *T_c_*), a dominant balance approach is useful (Fisher, 2013). For sufficiently large *x*, the *xϕ_z_* and 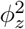 terms balance, and 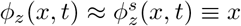. The superscript denotes that 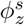 is the approximate solution taken within the *saturation* regime, in which *∂_x_ϕ_z_*(*x*, *t*) saturates approximately to 1. For smaller *x*, the nonlinear term *ϕ*^2^ is subdominant. We denote by 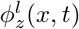 the solution to the linearized equation

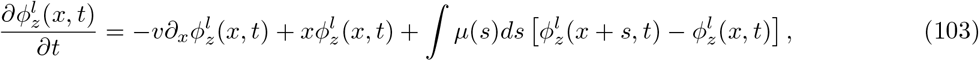

which is solved exactly by

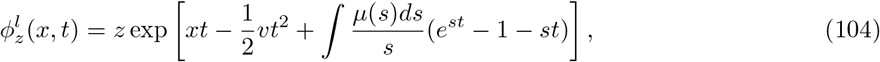

given the initial condition *ϕ_z_*(*x*, 0) = *z*. This suggests the approximate solution

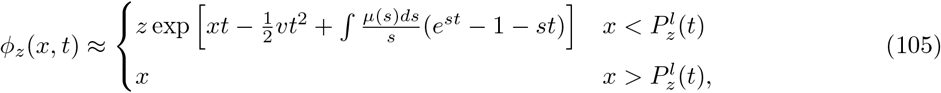

where 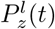 is defined such that 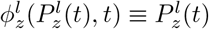. The quantity 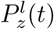 can be interpreted as the location of the time-dependent shoulder of the solution given in Eq. (105).

For 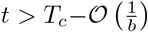, a dominant balance approach is not suitable, and Eq. (105) is not a good approximation. Even if the nonlinear term is subdominant—that is, even if 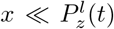, and 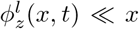 *x*—the linearized solution 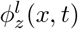 may be a very poor approximation to the actual solution *ϕ_z_*(*x*, *t*). The reason is that in obtaining Eq. (104) from Eq. (103), 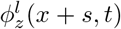 is also assumed to satisfy the same (linearized) equation. For *x* > *P_z_*(*t*) – *s*, the actual *ϕ_z_*(*x* + *s*, *t*) may be better approximated by 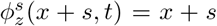 as opposed to 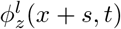. This discrepancy results in a time-derivative *∂_t_ϕ^l^*(*x*, *t*) which differs from that given by Eq. (103). Over time, this discrepancy propagates down to lower *x*, and for *t* > *T_c_*, the actual solution *ϕ_z_*(*x*, *t*) will differ qualitatively from the linearized solution 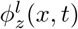 even for *x* well below 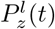.

This issue is considered at length by Fisher (2013), in which the following properties of the solution *ϕ_z_*(*x*, *t*) are identified for *t* > *T_c_*:

1. Shortly after *t* ≈ *T_c_*, on a timescale 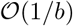, *ϕ_z_*(*x*, *t*) converges to *ϕ*_*P_z_*(*t*)_(*x*), with *ϕ*_*P_z_*(*t*)_(*x*) closely resembling the long-time fixed-point *w*(*x*) ≡ lim_*t*→∞_*ϕ*_*P_z_*(*t*)_(*x*), but with a shoulder at *P_z_*(*t*) rather than at *x_c_* = lim_*t*→∞_ *P_z_* (*t*).
2. The location *P_z_*(*t*) of the time-dependent shoulder converges to its fixed-point value *x_c_* with rate 1/*T_c_*:

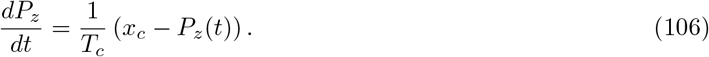
3. At long times, the solution *ϕ_z_*(*x*, *t*) depends on *z* only via the initial condition *P_z_*(*T_c_*) to Eq. (106). This can be approximated by taking 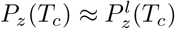, with

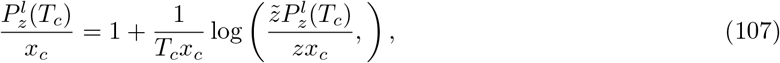

where 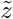 is defined such that 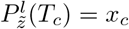. *P_z_*(*T_c_*) can be further approximated by

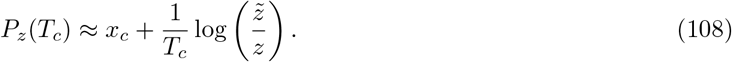
4. The integral Φ_0_(*z*, *t*) ≡ ∫ *f*(*x*)*ϕ_z_*(*x*, *t*)*dx* can be approximated as

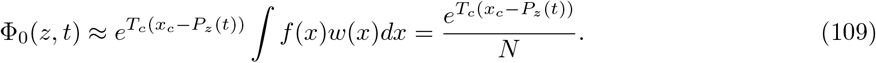 We note that Fisher (2013) considers a model in which the population size is not fixed, and the analog of ∫ *f*(*x*)*w*(*x*)*dx* in Eq. (109) can fluctuate; here, we assume ∫ *f*(*x*)*w*(*x*)*dx* = 1/*N* throughout, since the population size is assumed constant in our model.

These properties imply that for *b*(*t* – *T_c_*) ≫ 1,

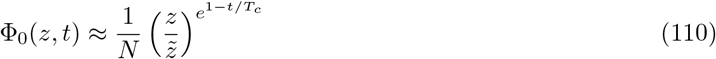

which is a key result we will use in computing statistics of genetic diversity in the MSSM regime (in particular, in computing the site frequency spectrum at moderate and high frequencies). Most of our results follow from the fact Φ_0_(*z, t*) is a simple monomial of *z* with power *e*^1–*t/T_c_*^ (i.e., that 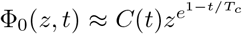, with *C*(*t*) independent of *z*). The full expression in Eq. (110) is useful for ensuring that this simple monomial dependence is valid over the appropriate region. We will see that our calculations of genetic diversity statistics in Appendix E involve integrals over *u* ≡ *N*Φ_0_(*z, t*) which are dominated by the region 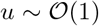. For use in this application, it will therefore be sufficient that Eq. (109) holds for the range 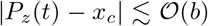 (over which *u* varies from *e*^−*T_c_b*^ to *e*^*T_c_b*^).

### A.3 Analysis of *ϕ_z_*(*x*, *t*)

Here, we provide a detailed analysis of *ϕ_z_*(*x*, *t*), explicitly demonstrating that the above properties hold for *b*(*t* – *T_c_*) ≫ 1, and extending this analysis to the case 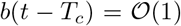. In both cases, we can analyze *ϕ_z_*(*x*, *t*) in a way similar to our analysis of *f*(*x*) and *w*(*x*) by defining

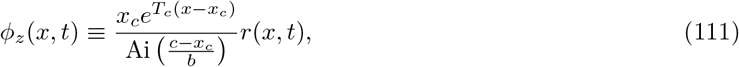

and approximating the resulting differential equation for *r*(*x*, *t*). Note that the constant factors in our definition of *r*(*x*, *t*) are chosen such that 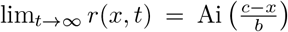; for simplicity in notation, we have suppressed the dependence of *r*(*x*, *t*) on *z*. Making the approximation 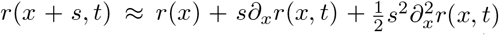 (which can be justified by noting that the resulting solution varies on the scale 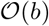 for relevant fitnesses *x*) and neglecting the nonlinear term 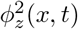, we have

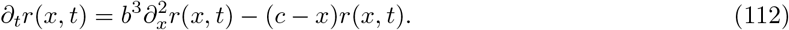

Our solution incorporates the nonlinearity in the equation for *ϕ_z_*(*x*, *t*) through the boundary conditions

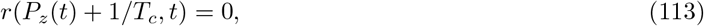

and

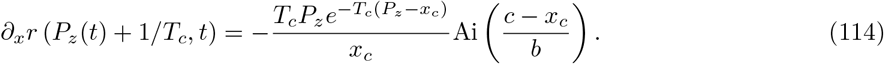

These boundary conditions ensure that *ϕ_z_*(*P_z_*(*t*), *t*) ≈ *P_z_*(*t*), and that rapid variation in *e^T_c_x^* is approximately canceled by variation in *r*(*x*, *t*) at *x* = *P_z_*(*t*), such that *ϕ_z_*(*P_z_*(*t*), *t*) is approximately linear at *x* = *P_z_*(*t*).

An approximate solution to Eq. (112) can be written as

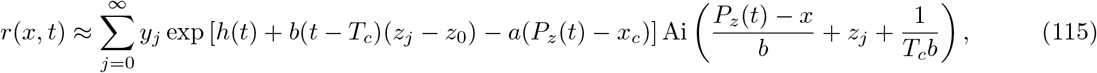

where *h*(*t*) satisfies

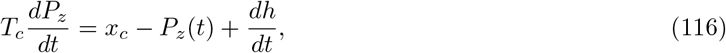

and the *y_j_* are arbitrary coefficients. Here, by *z_j_* we denote the (all negative) zeros of the Airy function satisfying Ai(*z_j_*) = 0, in order of increasing magnitude with *j*. To be more precise, Eq. (115) with *h*(*t*) satisfying Eq. (116) approximately solves Eq. (112), provided that 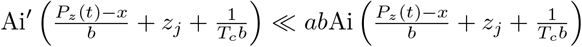, which is satisfied in the region of interest. The boundary condition *r*(*P_z_* + 1/*T_c_*, *t*) = 0 is automatically satisfied, while the boundary condition in Eq. (114) implies that

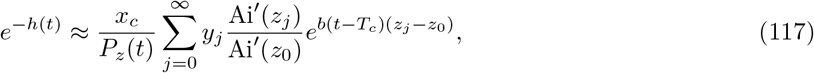

which, under the assumption *P_z_*(*t*) – *x_c_* ≪ *x_c_*, can be differentiated to yield

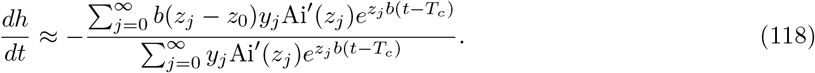

#### Long times: *b*(*t* – *T_c_*) ≫ 1

The analysis simplifies considerably at long times such that *b*(*t* – *T_c_*) ≫ 1. The sums in Eq. (115) and Eq. (117) are dominated by *j* =0 terms, with remaining terms exponentially suppressed. We therefore have

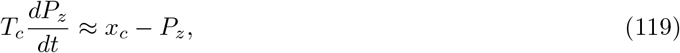

which is solved by

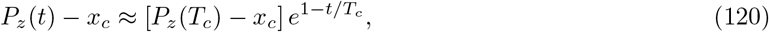

and the quantity *r*(*x*, *t*) simplifies to

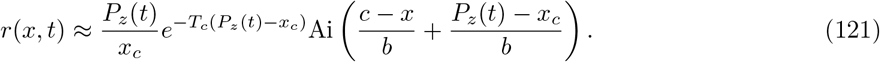

From Eq. (121) it follows that 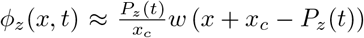; the quantity Φ_0_(*z, t*) ≡ ∫ *f*(*x*)*ϕ_z_*(*x*, *t*)*dx* can therefore easily be evaluated using the same calculation carried out in Appendix D to obtain *p*_fix_(*s*), with the result

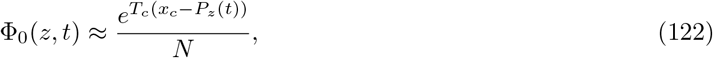

provided that |*P_z_*(*t*) – *x_c_*| ≪ *b*. Using the *P_z_*(*T_c_*) in Eq. (108), Eq. (122) further simplifies to

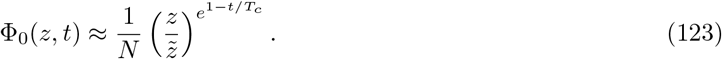

#### Intermediate times 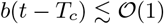

At intermediate times such that 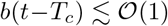, multiple terms contribute substantially to the sums in Eq. (115) and Eq. (117). As a result, we need to determine the appropriate coefficients *y_j_* in these sums, which can be done by matching our solution in Eq. (115) onto the solution for *t* < *T_c_* given in Eq. (105). To do so, we will integrate 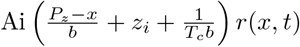 over the *x*-interval (–∞, *P_z_*(*t*) + 1/*T_c_*), using *r*(*x*, *t*) from both of these solutions, and exploit the orthogonality and completeness of appropriately shifted Airy functions. We will equate the two resulting integrals evaluated at a time *t* = *t*_0_ satisfying 1 ≪ (*T_c_* – *t*_0_)*b* ≪ *T_c_b*, and see that our final result will not depend on the precise choice of *t*_0_.

In particular, from the Airy equation it follows (using integration by parts) that

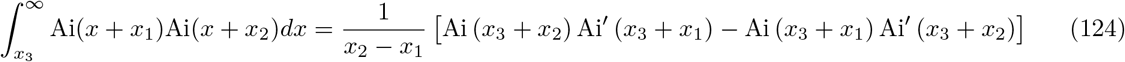

for arbitrary *x*_1_, *x*_2_ and *x*_3_. A further consequence is that 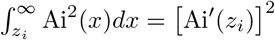, and also that

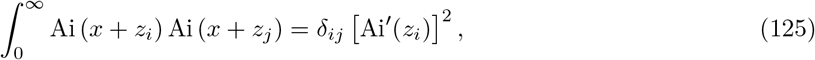

which establishes the orthogonality of the set of functions Ai(*x* + *z_j_*) on the interval (0, ∞). Using these properties, we can carry out the above-described matching, with the result

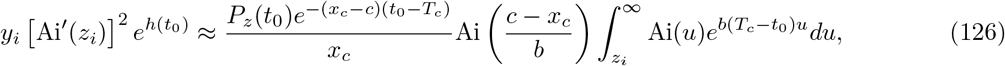

where, in obtaining the right-hand side, we made use of the relations 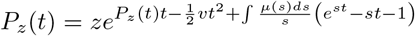 and *x_c_* ≈ *c* – *bz*_0_ – 1/*T_c_*. For sufficiently large *b*(*t*_0_ – *T_c_*), the integral on the right-hand side is dominated by *u* > *z*_0_, so that we can extend its lower limit of integration to –∞; the integral thus evaluates approximately to 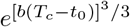. We can therefore take

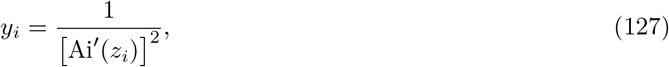

with

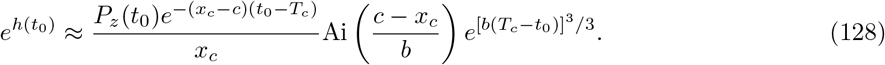

To simplify Eq. (128), we use the relation between *T_c_*, *N* and *x_c_* given in Eq. (31). This yields

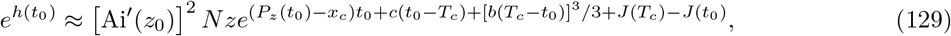

where 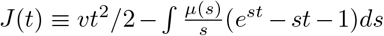. Expanding 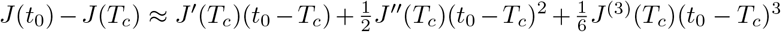, we then have

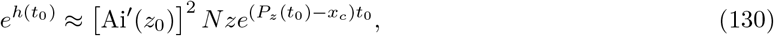

for –1 ≲ *b*(*t*_0_ – *T_c_*) < 0.

Eq. (127) can be substituted into Eq. (118) to yield

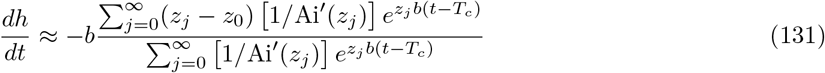

which is valid for intermediate times 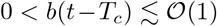 and long times *b*(*t* – *T_c_*) ≫ 1, and which determines the time-dependence of *P_z_*(*t*) through Eq. (116). Importantly, from Eq. (131) it follows that 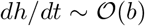 for 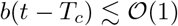; from Eq. (116), it then follows that

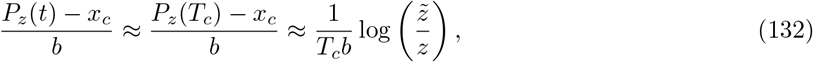

for times 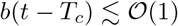. Eq. (116) can thus be integrated to yield, for times 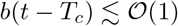,

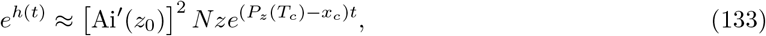

which does not depend on the choice *t*_0_. Substituting Eq. (133) and Eq. (127) into Eq. (115), we thus have

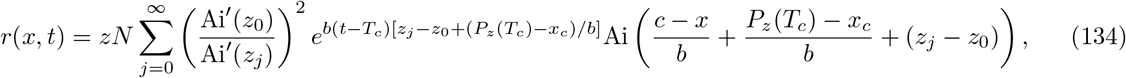

for 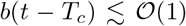. We can further simplify Eq. (134) by replacing (*P_z_*(*T_c_*) – *x_c_*)/*b* with 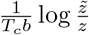. In particular, evaluating *ϕ_z_*(*x*, *t*) at *z* = *ζ*/*N*, we have

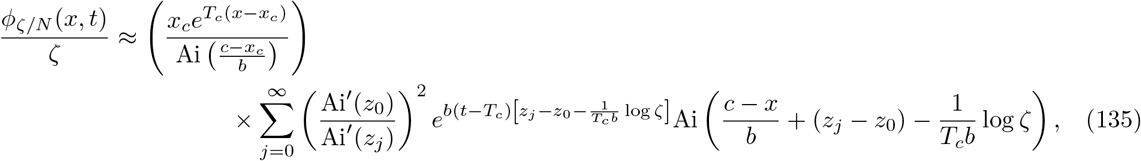

since (as discussed below) 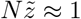 to logarithmic accuracy.

### A.4 Average rate of adaptation

Finally, we conclude this Appendix by briefly reviewing the procedure for computing *v* in terms of *N* introduced by Fisher (2013). While some of that work considers a model in rate of adaptation *v* is fixed and the population size can vary, Fisher (2013) notes that for a fixed population size *N*,

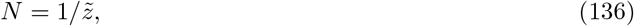

where 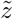 depends implicitly on *v*, and can be defined such that given the initial condition 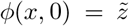, *ϕ*(*x*, *t*) converges to the eventual fixed-point *w*(*x*) with zero amplitude of the slowest eigenvector. Given the properties of the solutions *ϕ_z_*(*x*, *t*) discussed in the previous Subsection, it follows from this definition that 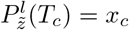, where 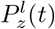 is the location of the time-dependent shoulder obtained by matching the linearized and saturated solutions in Eq. (105). That is,

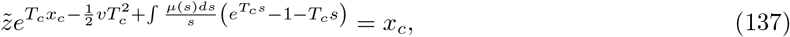

which yields

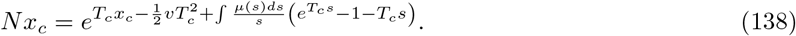

Apart from a factor of *T_c_b*Ai′(*z*_0_) multiplying *N*, Eq. (138) precisely matches our result obtained for the MSSM regime. Because of the logarithmic dependence of most quantities of interest on *N*, this is a relatively minor difference (even though *T_c_b* ≫ 1). As in our case, to solve for *v* using Eq. (138) requires another relation between *x_c_* and *T_c_* (as well as *μ*(*s*)). Within the “high-speeds” regime, Fisher (2013) obtains the same relation *x_c_* ≈ *c* + *b*|*z*_0_| – 1/*T_c_* that we obtain within the MSSM regime, using a solvability condition for *w*(*x*) which is exact asymptotically up to ambiguities of order 1/*T_c_* due to fluctuations.

## Appendix B DETERMINATION OF *v* AND *x_c_*

In this Appendix, we enforce the relevant self-consistency conditions to solve for *v* and *x_c_*, given the approximate solutions *f*(*x*) and *w*(*x*) obtained in Appendix C. Our presentation largely parallels the analysis presented by Tsimring et al. (1996) for the determination of *v* within the infinitesimal regime. We reproduce this derivation in detail because with a slight modification, this computation can be directly applied to yield *v* and *x_c_* within the MSSM regime.

In particular, we make use the following properties of *f*(*x*) identified within both limits: (i) *w*(*x*) ≈ *x* for *x* > *x_c_*, and (ii) within the region dominating 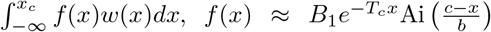 and 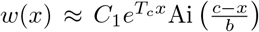, with constant factors *B*_1_ and *C*_1_ independent of *x*. In describing the infinitesimal approximation, (ii) is to be interpreted along with the substitutions 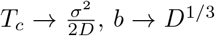, and 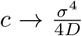 in describing the MSSM regime, (ii) is to be interpreted with the definitions for *T_c_*, *b*, and *c* used throughout the main text. These substitutions allow for an analysis of the two limits in parallel. In both cases, we assume *T_c_b* ≫ 1. In the MSSM approximation, *T_c_b* ≫ 1 is explicitly assumed, while in the infinitesimal approximation, *T_c_b* ≫ 1 follows from the additional assumption *ND*^1/3^ ≫ 1. The majority of work on the infinitesimal regime has focused on the case *ND*^1/3^ ≫ 1; we thus focus on this case in our consideration of the infinitesimal regime. An analysis of the *ND*^1/3^ → 0 limit, as well as a numerical consideration of the arbitrary *ND*^1/3^ case, is conducted by Good and Desai (2013) and Good et al. (2014). Finally, we make use of the normalization constants *B*_1_ obtained in Appendix C. With the substitutions assumed here,

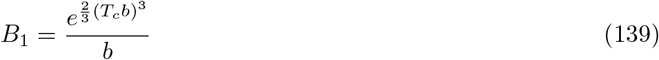

within the infinitesimal approximation, and

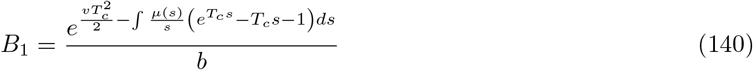

within the MSSM approximation.

The constant *C*_1_ and the interference threshold *x_c_* can be determined (in terms of *T_c_*, *b* and *c*) by enforcing the continuity of *w*(*x*) and *w*′(*x*) at *x* = *x_c_* (Good and Desai, 2014). Continuity of *w*(*x*) at *x_c_* requires that

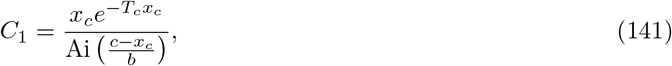

and continuity of *w*′ (*x*) at *x* = *x_c_* requires that

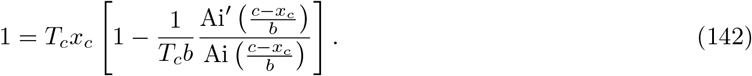

Given *T_c_* and *μ*(*s*), *x_c_* can be obtained as a numerical solution to Eq. (142). An explicit expression for *x_c_* in terms of *T_c_* and *μ*(*s*) can be obtained by making an ansatz that *T_c_x_c_* ≫ 1, such that

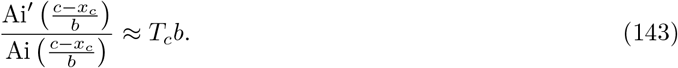

We will assume that *x_c_* < *c* + *b*|*z*_0_|, where *z*_0_ ≈ –2.34 is the least negative zero of Ai(*z*); this ensures that the patched *w*(*x*), as well as the cutoff *f*(*x*), are both nonnegative for all *x*. For *y* > *z*_0_, *Ai*′(*y*)/Ai(*y*) is only large and positive for *y* close to *z*_0_, motivating an expansion of 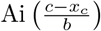 around 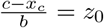. Retaining only the lowest-order term of this expansion yields

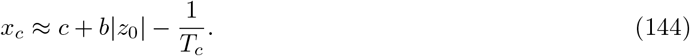

Eq. (144) is consistent with our ansatz *T_c_x_c_* ≫ 1, given our additional assumption that *T_c_b* ≫ 1 (along with the positivity of *c* implied by its definition), and is therefore justified.

The consistency condition 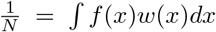, which enforces that a neutral mutation fixes with an unbiased probability 1/*N*, can be used to relate the quantities *T_c_*, *b* and *c*—and thus *v*—to *N* and *μ*(*s*). This consistency condition can be expressed as

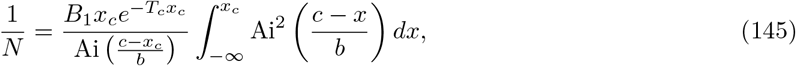

which evaluates to

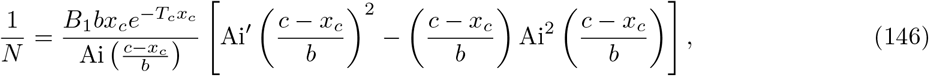

or, approximately, to

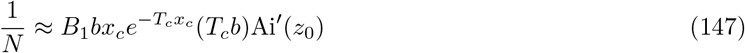

given the location of *x_c_* identified in Eq. (27). Substituting in the appropriate normalization constant *B*_1_ for the infinitesimal approximation, Eq. (147) simplifies to

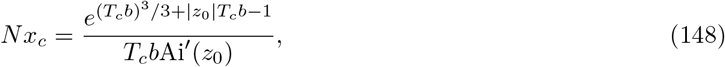

while for the MSSM approximation, Eq. (147) reduces to

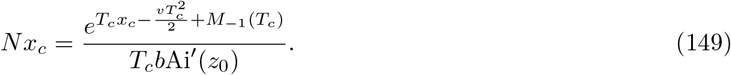

These are the key equations, in combination with Eq. (142), which relate *T_c_* (and thus *v*) to *N* and *μ*(*s*) in the two approaches. We discuss the numerical solution of these equations in Appendix I.

### B.1 An asymptotic approximation in the *N* → ∞ limit

Finally, we conclude this Appendix by analyzing Eq. (149) to obtain an asymptotic expression for *T_c_* in the limit *N* → ∞. We focus on the case in which the DFE consists of a single beneficial effect—that is, we assume *μ*(*s*) = *Uδ*(*s* – *s_b_*). In the limit *N* → ∞, *T_c_x_c_* → *T_c_v* and we have

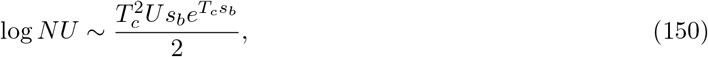

which has the solution

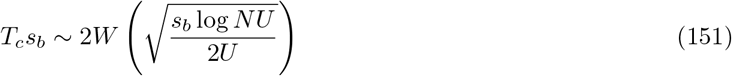

where *W* is the Lambert-*W* function satisfying *W*(*z*)*e*^*W*(*z*)^ = *z*. Using the asymptotic approximation *W*(*z*) ~ log *z* – log log *z* for large *z*, we have

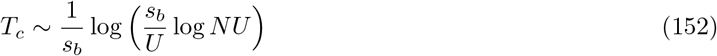

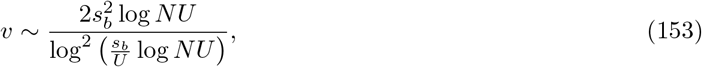

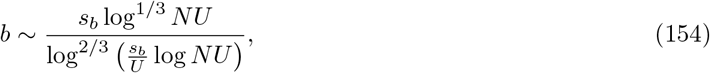

and

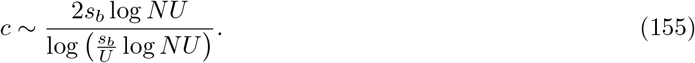

Note that in the limit *N* → ∞, the conditions of validity *s_b_* ≪ *b* and *T_c_b* ≫ 1 of the MSSM approximation are in fact satisfied.

## Appendix C APPROXIMATION OF *f*(*x*) AND *w*(*x*) USING TRANSFORM METHODS

In this Appendix, we use transform methods to approximate *f*(*x*) and *w*(*x*) within both the infinitesimal and MSSM regimes. To do so, we begin by reviewing the solutions, provided by Fisher (2013), for the Laplace transforms 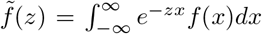 and 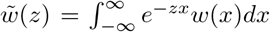. We then demonstrate that the infinitesimal approximation solutions *f*(*x*) and *w*(*x*) can be obtained from inverse Laplace transforms of 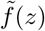 and 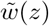, respectively, by approximation of the relevant contour integrals. Using a different approximation of these contour integrals, we reproduce the MSSM approximate solutions *f*(*x*) and *w*(*x*) found in the main text. In doing so, our main purpose is to obtain precise conditions of validity for both the infinitesimal and MSSM approximations, expressed as integrals involving *μ*(*s*).

Note that in the definitions we take of 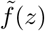 and 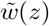, *f*(*x*) and *w*(*x*) are assumed to satisfy

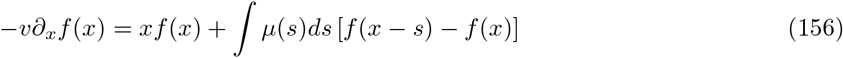

and

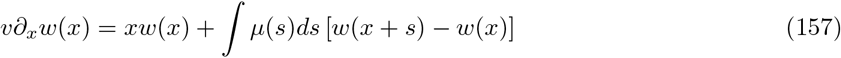

respectively. Here *f*(*x*) and *w*(*x*) denote the formal solutions to Eq. (156) and Eq. (157), respectively. That is, within the definition of 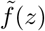, *f*(*x*) does not possess a “cutoff” at *x_c_*; similarly, within the definition of 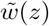, *w*(*x*) does not tend to *x* for *x* ≫ *x_c_* (which is the case for the solution to Eq. (10)). The transform methods used here are employed only to obtain approximate solutions Eq. (156) and Eq. (157). These approximate solutions can then be patched with the appropriate behavior for *x* > *x_c_* to obtain the full approximate solutions *f*(*x*) and *w*(*x*) which are used to compute quantities of interest.

The transforms 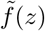 and 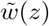 satisfy

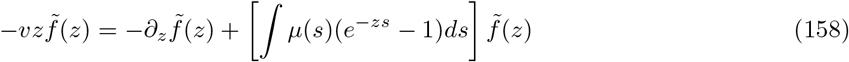

and

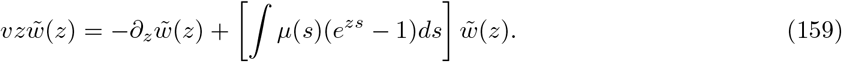

In obtaining Eq. (158) from Eq. (5) we assume that *e*^−*zx*^*f*(*x*) → 0 as *x* → ∞ and as *x* → –∞. Likewise, in obtaining Eq. (159) from Eq. (10) we assume that *e*^−*zx*^*w*(*x*) → 0 as *x* → ∞ and as *x* → –∞. Together, these conditions restrict the range of *z* for which Eq. (158) and Eq. (159), and thus our solutions 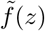 and 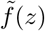, are valid. However, if *f*(*x*) and *w*(*x*) are reasonably well-behaved, these conditions are satisfied when 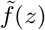 and 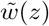 (which are defined by integrals over *e*^−*zx*^*f*(*x*) and *e*^−*zx*^*w*(*x*), respectively) take on finite values; this is in fact satisfied by our solutions 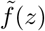 and 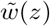, within the region of *z* of interest to us.

Eq. (158) and Eq. (159) can each be integrated to yield

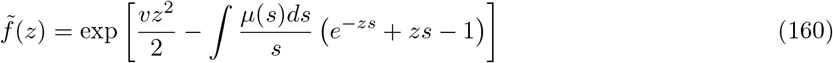

and

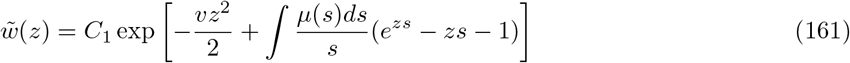

(Fisher, 2013), where the constant of integration for 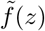 is set by the requirement that ∫ *f*(*x*)*dx* = 1. The constant of integration for 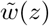 will be determined along with the location *x_c_* of the interference threshold by enforcing that *w*(*x*) and *w*′(*x*) are continuous at *x_c_*.

To enforce this condition—as well as to determine *v* and other dynamical quantities of interest—the functional forms of *f*(*x*) and *w*(*x*) are needed (as opposed to their transforms 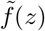 and 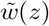)). The distribution *f*(*x*) follows from 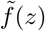, using the standard formula for an inverse Laplace transform, as

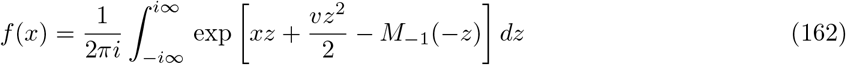

and

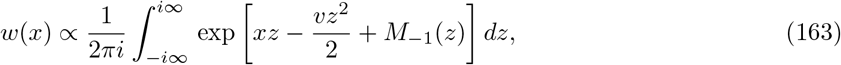

where, as defined by Fisher (2013), 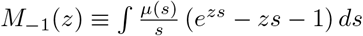. We will also define

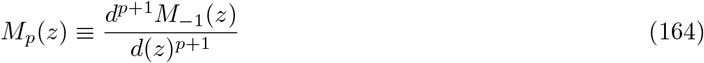

for *p* ≥ 0, so that *M*_0_(*z*) ≡ ∫ *μ*(*s*) (*e^zs^* – 1) *ds* and *M_p_*(*z*) = ∫ *μ*(*s*)*s^p^e^zs^ds* for *p* ≥ 1.

### C.1 Approximation of *f*(*x*) and *w*(*x*) within the Infinitesimal Approximation

We now turn to approximate Eq. (162) and Eq. (163) within the infinitesimal regime. Eq. (162) can be rewritten as

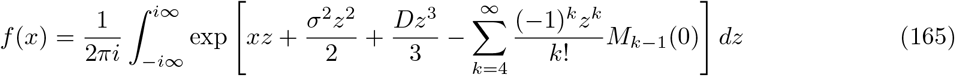

by expanding the exponential *e*^−zs^ within the integral over *s*, and defining 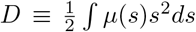, and *σ*^2^ ≡ *v* – *U*〈*s*〉. As in the rest of this article, we limit our attention to the case of the infinitesimal regime in which *ND*^1/3^ ≫ 1, or equivalently, in which *σ* ≫ *D*^1/3^ (for a brief discussion of the case in which *ND*^1/3^ is not large, see Appendix B). By making the substitution 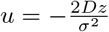, (165) can be alternatively expressed as

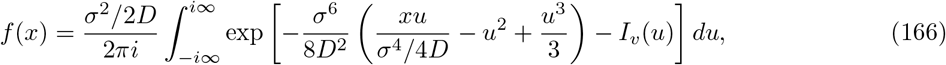

where we have defined

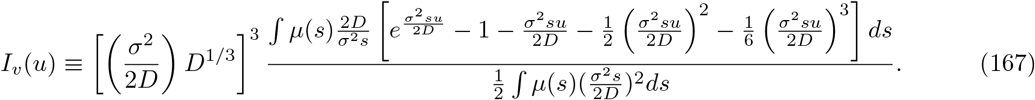

To approximate *f*(*x*) within the infinitesimal approximation, we neglect the contribution of *I_v_*(*u*) to the exponent of Eq. (166); the remaining integral can be evaluated, yielding

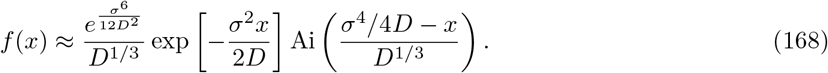

This can be justified by ensuring that *I_v_*(*u*) ≪ 1 throughout the region around saddle points dominating the integral in Eq. (166). To see this, note that the integration contour can be deformed to pass through one or more saddle points along a path of steepest descent, with the integral then dominated by a relatively narrow region around those saddle points. In general the suitability of this approximation may depend sensitively on *x*; below we focus on the region 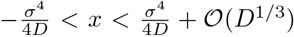, which will comprise the “bulk” of *f*(*x*) as well as the “fixation class” (the region dominating ∫ *f*(*x*)*w*(*x*)*dx*).

The relevant saddle points are located at *u*\* which satisfy

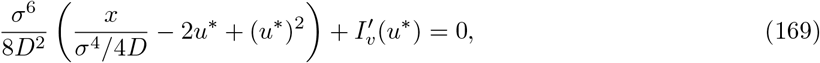

and which can be approximated as

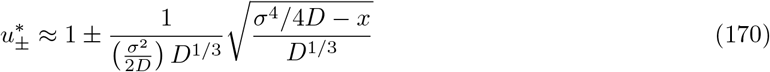

after further assuming that 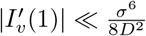 (or more precisely, that 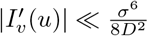 for 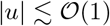, along with the assumption 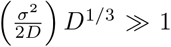, which turns out to be equivalent to the assumption *ND*^1/3^ ≫ 1). Upon making the appropriate deformation of the integration contour, the integral in Eq. (166) is dominated by a region of width Δ around 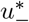 (for 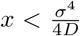) or a region of width Δ around *both* 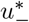 and 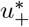 (for 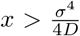), where

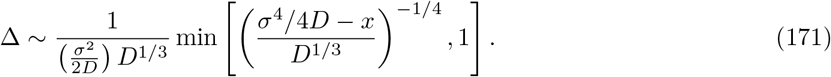

It follows that the region 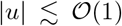 dominates the integral in Eq. (166), throughout the region of *x* considered here. The conditions |*I_v_*(*u*)| ≪ 1 and 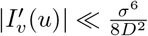 for 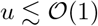 then justify the approximations yielding Eq. (168); these conditions are closely approximated by the conditions |*I_v_*(1)| ≪ 1 and 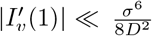.

A similar approximation can be made to obtain *w*(*x*); Eq. (163) can be rewritten as

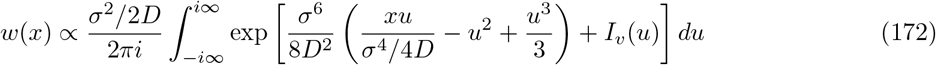

with *I_v_*(*u*) again defined by Eq. (167), which simplifies to

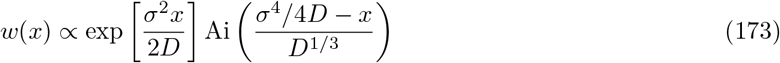

provided, again, that *I_v_*(*u*) is negligible within the region that dominates the integral in Eq. (172). For a particular *x*, this region may differ from the region of *u* dominating the integral in Eq. (166): for *x* > *σ*^4^/4*D*, these regions are similar, while for *x* < *σ*^4^/4*D*, the integral is dominated by the region around 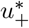 (as opposed to the region around 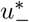, for *f*(*x*)). However, throughout the region 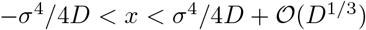 of *x* considered, the same conditions |*I_v_*(1)| ≪ 1 and 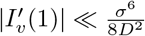 justify the neglect of *I_v_*(*u*) within the region dominating Eq. (172).

Note that the condition 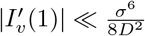 can be written

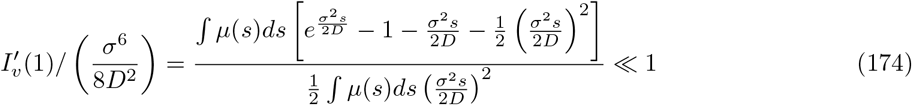

which is, roughly speaking, satisfied if *σ*^2^*s*/2*D* ≪ 1 throughout the region dominating ∫ *μ*(*s*)*s*^2^*ds*. Because, in the infinitesimal regime, *T_c_* ≈ *σ*^2^/2*D* (Neher and Hallatschek, 2013), Eq. (174) essentially requires that *T_c_s* ≪ 1 for a substantial majority of available fitness effects *s*, as stated in the main text. The other condition |*I_v_*(1)| ≪ 1 has a similar interpretation.

### C.2 Approximation of *f*(*x*) and *w*(*x*) within the MSSM regime

As in the previous Subsection, here we approximate the integrals in Eq. (162) and Eq. (163) by first approximating the integrand in the vicinity of its saddle points, and then employing an integral representation of the Airy function. The integrand in Eq. (162) possesses saddle points whose locations *z_s_*(*x*) satisfy

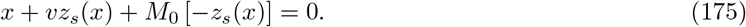

Validity of the infinitesimal approximation requires that the relevant saddle points *z_s_* are small enough in magnitude to justify a Taylor approximation of Eq. (175) to second-order in *z_s_*. Within the MSSM regime, this Taylor approximation is not necessarily justified. Instead, in the next Subsection we make a local approximation to Eq. (175), valid for *x* within 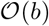 of *c*, and obtain approximate expressions for *f*(*x*) and *w*(*x*) within this fixation class. Using a similar approach, we then turn to obtain approximate solutions for *f*(*x*) for *x* within the “bulk” of the fitness distribution.

#### Solution within the Fixation Class: 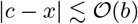

To approximate Eq. (162) for *x* within the fixation class, we shift our integration contour to the line from –*T_c_* – *i*∞ to –*T_c_* + *i*∞, and expand the exponent of our integrand around its value at *z* = –*T_c_*, the relevant (and only) saddle point of the integrand for *x* = *c*. That is,

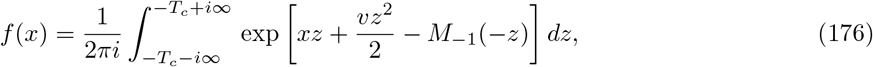

which can be rewritten as

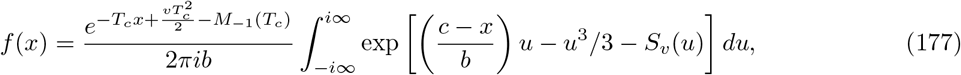

with

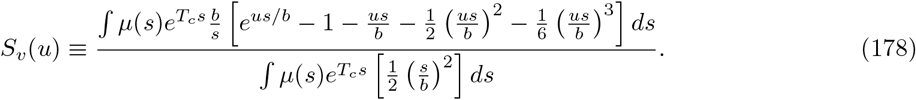

Note that Eq. (177) is obtained using the expansion 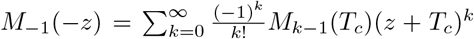 and the substitution *u* ≡ –*b*(*z* + *T_c_*). An integral representation of the Airy function, 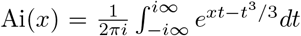, can then be applied to yield

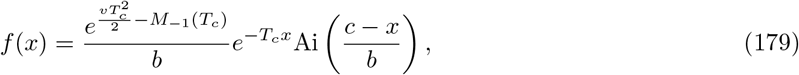

provided that *S_v_*(*u*) can be neglected throughout the region (within the vicinity of saddle points) that dominates the integral in Eq. (177). Saddle points are located at *u*\* which satisfy

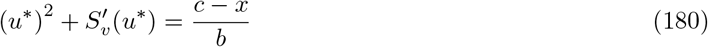

with

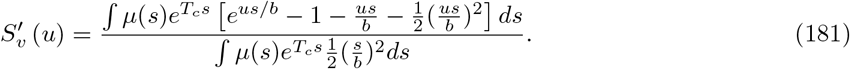

We approximate Eq. (180) by

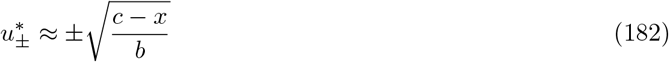

which, provided that 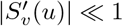 for 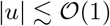, holds self-consistently throughout the region 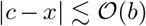 of *x* considered. The region of *u* within 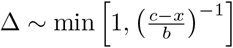 of *u*\* dominates the integral in Eq. (177); the approximation yielding Eq. (179) is then justified throughout the region of *x* considered if |*S_v_*(1)| ≪ 1 and 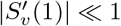. A similar calculation yields

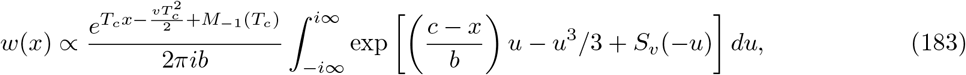

which we approximate by 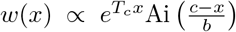 (when *S_v_*(–*u*) can be neglected. The same conditions |*S_v_*(1)| ≪ 1 and 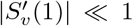 identified above can be shown to ensure that *S_v_*(–*u*) is negligible in the relevant regions.

#### Solution within the Bulk Class: 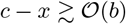

Here, we obtain an approximate solution for *f*(*x*) for *x below* the fixation class. Similar manipulations can be carried out to yield an approximate solution for *w*(*x*) within this region. Within this region, approximating the locations of saddle points—solutions *z_s_*(*x*) to Eq. (175)—as in Eq. (180) is not justified. Instead we obtain approximations to the integrals in Eq. (162) and Eq. (163) in terms of *z_s_*(*x*), which must be obtained numerically. Note that for each *x* < *c* there are two real solutions *z_s_*(*x*) to Eq. (175): one in which *z_s_*(*x*) < –*T_c_* and another in which *z_s_*(*x*) > –*T_c_*. Throughout the rest of this section, we denote by *z_s_* the solution *z_s_*(*x*) to Eq. (175) such that *z_s_*(*x*) > –*T_c_*, which will be the relevant saddle point through which it is useful to deform the integration contour.

After shifting the integration contour to the line from *z_s_* – *i*∞ to *z_s_* + *i*∞ and making the substitution 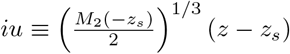, Eq. (162) can be rewritten

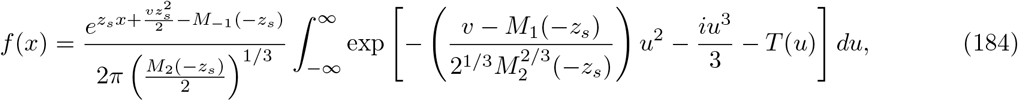

where 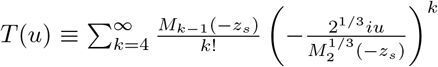. We wish to approximate *f*(*x*) by neglecting *T*(*u*). The integral representation 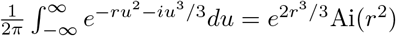—which assumes Re(*r*) ≥ 0—can then be applied to yield

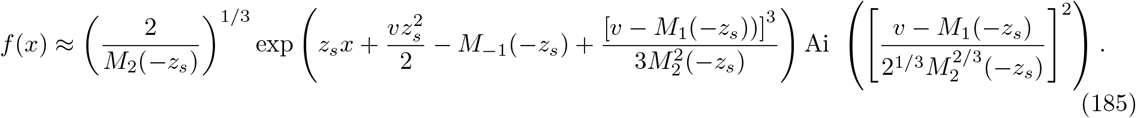

When 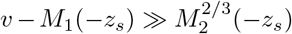, an asymptotic approximation of the Airy function can be applied to yield

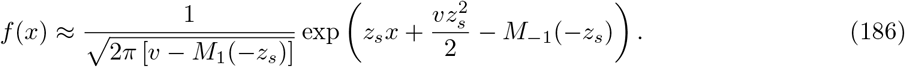

For *x* sufficiently close to 0, Eq. (186) reduces to the expression for *f*(*x*) obtained within the infinitesimal approximation—that is, Eq. (168). In particular, for sufficiently small *x*,

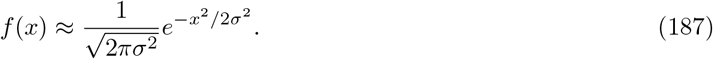

Note that Eq. (187) requires an approximation for *σ*^2^. Taking *σ*^2^ = *v* – *U*〈*s*〉, with *v* computed using the MSSM approximation as described in the main text, should yield a good approximation to *f*(*x*) within its bulk (provided the conditions of validity of the MSSM approximation are satisfied).

## Appendix D MUTATIONAL FIXATION PROBABILITIES

In this Appendix, we evaluate

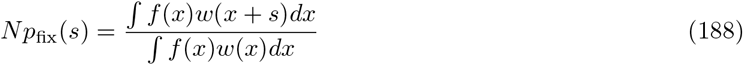

using the approximate expressions for *f*(*x*) and *w*(*x*) identified within the MSSM regime. In particular, we make use of our result that 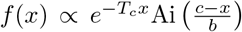, and 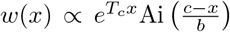, within the region of *x* dominating ∫ *f*(*x*)*w*(*x*)*dx* (in which 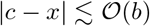). We focus throughout on the case |*s*| ≪ *b*; as discussed in the main text, validity of the MSSM approximation requires that |*s*| ≪ *b* for a substantial majority of fixed mutations. Given the assumption |*s*| ≪ *b*, the region dominating ∫ *f*(*x*)*w*(*x* + *s*)*dx* is approximately the same as the region dominating ∫ *f*(*x*)*w*(*x*)*dx*, justifying the use of our approximate results for *f*(*x*) and *w*(*x*).

We first consider the case in which *s* < 0, so that

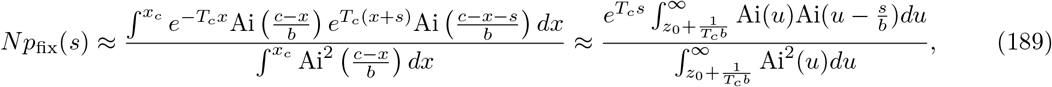

where in the last equality we used our approximate result for *x_c_* in Eq. (27). Expanding 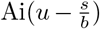 to second order in 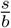,

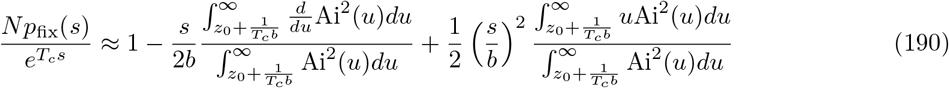

which simplifies to

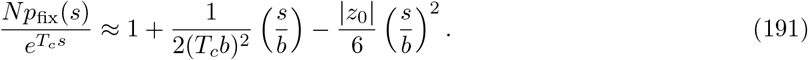

The *s* > 0 case is a bit more complicated, since ∫ *f*(*x*)*w*(*x* + *s*)*dx* receives a contribution from the region of *x* in which *x* + *s* > *x_c_*. In this case,

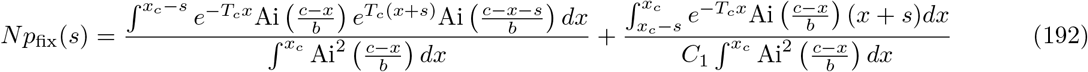

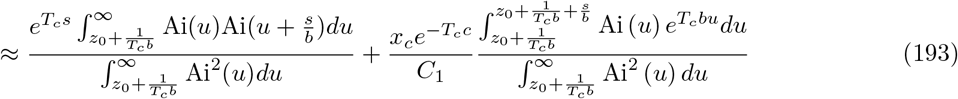

with 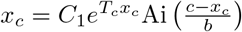. The first term above can be easily evaluated using our *s* < 0 result. The second term can be approximated by noting that the integrand Ai(*u*)*e^T_c_bu^* peaks at 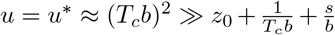. An application of Watson’s Lemma then yields

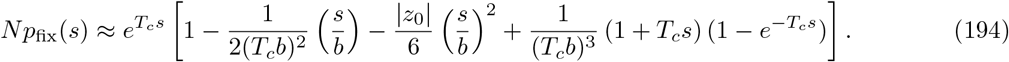

In both the *s* > 0 and *s* < 0 cases, then,

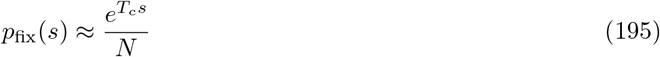

given the assumption *T_c_b* ≫ 1 made throughout (as well as the assumption *s* ≪ *b* made here).

We note that we have defined *T_c_* according to *N* ∫ *μ*(*s*)*se^T_c_s^ ds* ≡ *v* = *N* ∫ *μ*(*s*)*sp*_fix_(*s*)*ds*. Unless *μ*(*s*) consists of only a single effect size, this definition does not necessarily imply that *Np*_fix_(*s*) = *e^T_c_s^*. We have shown above that *Np*_fix_(*s*) ≈ *e^T_c_s^* in fact holds for *s* ≪ *b*; together with the fact that ∫ *μ*(*s*)*p*_fix_(*s*)*ds* is dominated by the region of *s* ≪ *b*, this implies that approximate self-consistency of *N* ∫ *μ*(*s*)*se^T_c_s^ ds* ≡ *v* = *N* ∫ *μ*(*s*)*sp*_fix_(*s*)*ds* is achieved. Our analysis only explicitly enforced that *p*_fix_(0) = 1/*N*, and yet a consequence of this requirement is that a self-consistent rate *v* is obtained. The equivalence of the self-consistency conditions *p*_fix_(0) = 1/*N* and *v* = *N* ∫ *μ*(*s*)*sp*_fix_(*s*)*ds* has been identified by Good et al. (2012) to hold *exactly* in the “tunable constraint” models introduced in Hallatschek (2011), provided that an analog of ∫ *f*(*x*)*w*(*x*)*dx* = 1/*N* is taken as the “tuned constraint”; here these conditions are approximately equivalent.

## Appendix E STATISTICS OF GENETIC DIVERSITY

In this Appendix, we provide derivations of the merger probabilities 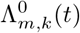 that follow from our evolutionary model. We use these results to conclude that, after an initial delay period during which negligible coalescence events occur, genealogies are well-described by the Bolthausen-Sznitman coalescent (Bolthausen and Sznitman, 1998). In a similar calculation, we demonstrate a more direct correspondence with the BSC through the *partition structure* (which we define below) following from our evolutionary model. We then provide an explicit calculation of the neutral and selected site frequency spectra (SFS), separately obtaining the contribution to the SFS from mutations with ages *t* such that *b*(*t* – *T_c_*) ≫ 1, and from mutations with ages *t* such that 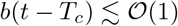. Finally, we conclude by briefly discussing how results from Fisher (2013) can be applied to obtain a *transition density* of the effective lead frequency *ν_L_*(*t*) defined in the main text.

### E.1 Merger Probabilities

The BSC is a type of Λ-coalescent, which in turn is a coalescent process that can be defined based on the merger rates λ_*m,k*_ at which any given *k*-tuple of *m* blocks coalesce (Berestycki, 2009). In the BSC,

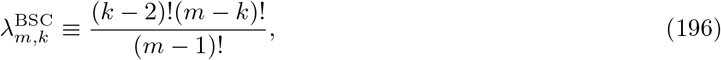

which can be contrasted to the Kingman coalescent, in which λ_*m,k*_ = *δ*_*k*,2_ (i.e. only pairwise merger events are allowed) (Berestycki, 2009). Following an approach similar to that taken by Neher and Hallatschek (2013), we can obtain the merger rates λ_*m,k*_ for our evolutionary model by first calculating the merger *probabilities* 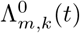. By 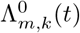, we denote the probability that of a sample of size *m*, a particular set of *k* individuals shares a common ancestor at *t* generations into the past, and the remaining *m* – *k* individuals do *not* trace back to that same ancestor. These can be computed as

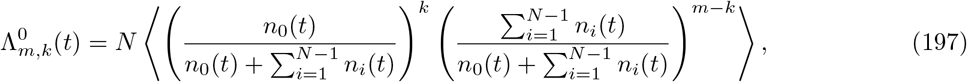

where the factor of *N* accounts for the *N* possible common ancestors of the *k* individuals which were alive at time *t* before the present generation. Here, the 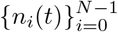 are the (stochastic) lineage sizes of the *N* individuals after a time *t*, which are identically distributed. Note also that our definition of 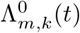 differs from the definition of 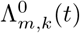 taken by Neher and Hallatschek (2013). In that work, 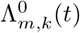 is defined as the probability that of a sample of *m* individuals, a given subset *k* of those individuals share a common ancestor, and that the remaining *m* – *k* individuals trace back to *m* – *k distinct* ancestors, at *t* generations. We take our definition because it enables us to obtain the same correspondence with the BSC observed by Neher and Hallatschek (2013), and because defined as such, the probabilities 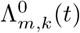 can be used to compute the site frequency spectrum.

To simplify Eq. (197), we employ the identity 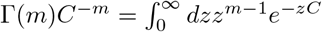 with 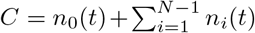, which yields

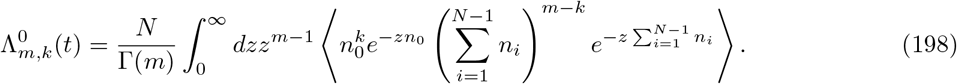

Assuming *n*_0_(*t*) and 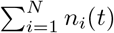 are independent of one another,

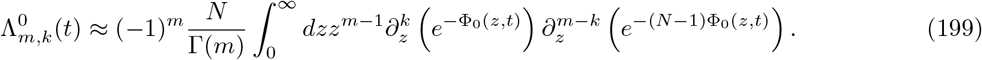

since, as we discuss in Appendix A, *e*^−Φ_0_(*z,t*)^ ≈ 〈*e*^−*zn*(*t*)^〉 (note that Φ_0_(*z,t*) ≡ ∫ *f*(*x*)*ϕ_z_*(*x, t*)*dx*). We discuss the behavior of Φ_0_(*z,t*) in Appendix A. The key results we use here are that, for *t* < *T_c_*, Φ_0_(*z,t*) is approximately linear in *z*, and for *t* > *T_c_*,

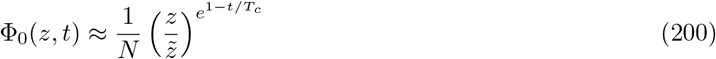

within the relevant region of *z* (note that for *t* > *T_c_*, our results primarily require only that Φ_0_(*z,t*) ≈ *A*(*t*)*z^*e*^1−t/T_c_^^* for some *z*-independent function *A*(*t*)).

Linearity of Φ_0_(*z,t*) in *z* implies that 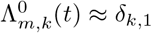 and thus that merger events are negligible for *t* < *T_c_*. For *t* > *T_c_*, Eq. (199) can be simplified by noting that Φ_0_(*z,t*) ≪ 1 in the relevant region of *z*, so that

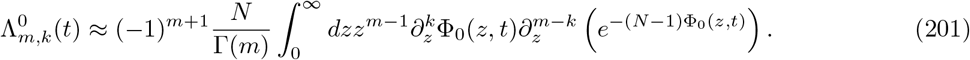

Substituting Eq. (200), and integrating by parts *b* – *k* times, yields

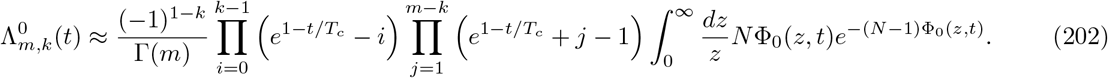

Using the substitution *u* ≡ *N*Φ_0_(*z, t*), the *z*-integral in Eq. (202) can be evaluated to give

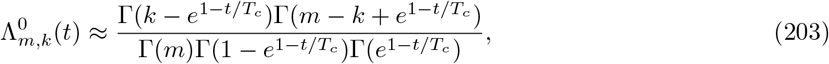

which, using the Euler reflection formula for Γ(*z*), can be further simplified to

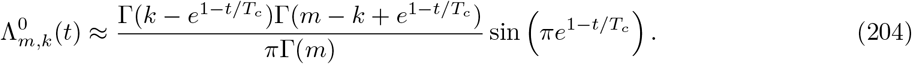

From Eq. (204) the merger *rates* 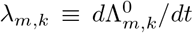 at *t* = *T_c_* (just after merger events occur at a non-negligible rate) follow as

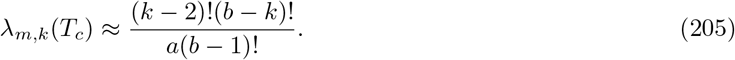

These are the BSC merger rates 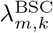, scaled by the overall timescale *T_c_*. The same correspondence is noted by Neher and Hallatschek (2013) to argue that genealogies within the infinitesimal regime resemble those of the BSC (although in that case, the overall timescale identified is *σ*^2^/2*D*, to which *T_c_* reduces in the infinitesimal regime), after an initial delay period during which few coalescence events occur. The initial delay period can be interpreted, looking backward in time, as the time required for the ancestors of typical individuals—those sampled from the “bulk” of the fitness distribution—to reach the high-fitness edge of the fitness distribution; once this occurs, coalescence proceeds as described by the BSC.

The probability *Q_k_*(*t*) ≡ Λ*_k,k_*(*t*) gives the probability with which *k* randomly chosen individuals share a common ancestor within *t* generations, and simplifies to

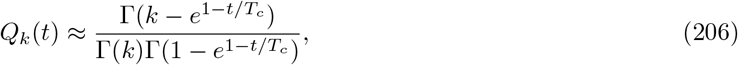

for *t* > *T_c_*. Apart from the replacement of *t* by (*t* – *T_c_*)/*T_c_*, these are precisely the full time-dependent *Q_k_*(*t*) which are obtained in the BSC (Mohle and Pitters, 2014; Pitman, 1999). Thus the correspondence with the BSC in our model extends beyond the correspondence of the instantaneous merger rates at *T_c_* generations. Note that *q_k_*(*T_k_*) ≡ *dQ_k_*/*dt* can be interpreted a probability distribution for the coalescence times *T_k_* of a sample of *k* individuals. In particular,

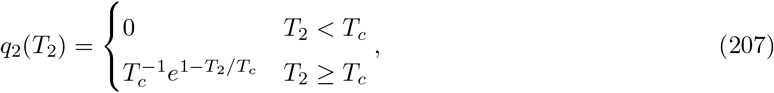

and therefore

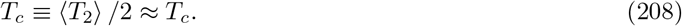

That is, the timescale *T_c_* of coalescence is the same as the *delay* timescale—the initial time during which negligible coalescence events occur. In the infinitesimal regime, the same correspondence (as well as the exponential distribution of pairwise coalescence times found in Eq. (207)) is inferred based on the instantaneous merger rates calculated at *T_c_* generations, and observed in simulations, by Neher and Hallatschek (2013).

We note that although our overall conclusions are quite similar to those of Neher and Hallatschek (2013), our calculation differs in an important way. In obtaining Φ_0_(*z, t*), Neher and Hallatschek (2013) approximate the solution *ϕ_z_*(*x, t*) to Eq. (62) using a dominant balance argument—in particular, by taking

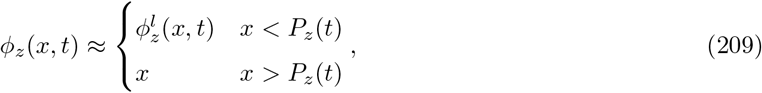

where 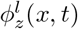 is the solution to Eq. (62) with the 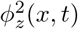 term neglected, and 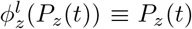 (so that the two solutions match up at *x* = *P_z_*(*t*)). Fisher (2013) argues that Eq. (209) is valid only for *t* < *T_c_*, and describes the *t* > *T_c_* behavior in detail (which we review in Appendix A). Neher and Hallatschek (2013) analytically demonstrate a correspondence with the BSC by calculating the instantaneous merger rates at *t* = *T_c_* (more precisely, at *t* = *σ*^2^/2*D* ≈ *T_c_* since they work within the infinitesimal regime), so that Eq. (209) may still be valid; however, extending their calculation to later times yields various pathological results (e.g diverging expectation values for coalescence times including 〈*T*_2_〉). Our calculation, which makes use of behavior of *ϕ_z_*(*x, t*) described by Fisher (2013), yields well-behaved predictions for the full time-dependence of the quantities *Q_k_*(*t*) and 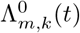, and therefore the site frequency spectrum. As a result, we are able to analytically demonstrate aspects of the correspondence with the BSC which are observed in simulations, and justified using heuristic arguments, by Neher and Hallatschek (2013).

We emphasize that in our demonstration of a correspondence with the BSC, a key step is noting that Eq. (200) holds over the region of *z* which dominates the relevant integrals. That is, we take

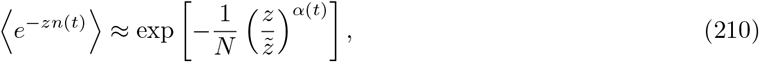

with *α*(*t*) = *e*^1−*t/T_c_*^. The same generating function is used by Fisher (2013) to describe fluctuations in the total size *N*(*t*) of a population, assuming its mean fitness increases at a fixed rate *v* (which can then be used to describe fluctuations in the rate of adaptation at fixed population size). As noted by Fisher (2013) for the case of a fluctuating population size, from Eq. (210) it follows that *n*(*t*) is drawn from a one-sided Lévy stable distribution with parameter *α*(*t*); the offspring number distribution—the pdf of *n*(*t*)—then falls off as *p*(*n*) ~ 1/*n*^1+*α*(*t*)^ (Nolan, 2018). In particular, at *T_c_* generations the offspring distribution falls off as 1/*n*^2^. This offspring number distribution—which is the same as that obtained for the Luria-Delbruck process (Yule, 1925)—is observed (on different timescales) for rapidly adapting populations lying in several regimes of the parameter space (Desai and Fisher, 2007; Kosheleva and Desai, 2013), and has been explained as arising from the exponential amplification of fit lineages (Neher and Hallatschek, 2013).

It is well-understood that the offspring number distribution *p*(*n*) ~ 1/*n*^2^ is related to the BSC (Schweinsberg, 2003); Hallatschek (2018) recently described a precise (neutral) stochastic process, involving an offspring number distribution of *p*(*n*) ~ 1/*n*^2^, which can be considered a forward-time dual of the BSC (much like Wright-Fisher diffusion can be considered a forward-time dual of the Kingman coalescent). This duality has been used to argue that, after coarse graining to the timescale on which *p*(*n*) ~ 1/*n*^2^, the genealogies of rapidly adapting populations can be described by the BSC. Interestingly, we have found above that Eq. (210) implies a correspondence with the BSC that goes beyond a correspondence on a coarse-grained timescale: after making reasonable approximations, the full time-dependence of the BSC partition structure follows from our evolutionary model. We provide further discussion of the relation between the dynamics under our model in the MSSM regime, and the dynamics under the model considered by Hallatschek (2018), in Subsection VI..3.

We note that the total population size *N* is fixed within our model, and therefore a 1/*n*^2^ offspring number distribution cannot strictly speaking apply. At minimum, the true offspring number distribution must have a cutoff at the total population size *N*. However, because only the relative *fractions* of the population comprised by any given lineage enter into our calculation of diversity statistics, we expect that Eq. (210) is still appropriate. We note also that Eq. (210) is only valid within a limited range of *z*; for instance, Eq. (210) yields pathological results both in the limit *z* → ∞ (which gives extinction probabilities) and in the limit *z* → 0 (which gives moments of *n*(*t*)). As we discuss in Appendix A, however, Eq. (210) is valid in the region of *z* which dominates integrals used to compute statistics of genetic diversity—roughly speaking, for *z* corresponding to *n*(*T_c_*) in the range *e^−T_c_b^* ≪ *n*(*T_c_*)/*N* ≪ *e^T_c_b^*. Lineages of this size—while atypically large, relative to the expected value 〈*n*(*T_c_*)〉 = 1—make the dominant contribution to coalescence probabilities.

#### Selected Merger Probabilities

Using this framework, we can also compute the probability 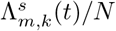 that, of a sample of *m* individuals, a given set of *k* individuals trace back to a *specific* common ancestor at *t* generations (and that the remaining *m* – *k* individuals do not trace back to that ancestor), conditioned on that common ancestor of the *k* individuals having acquired a mutation of effect size *s* at time *t* in the past. As noted in the main text, the probability 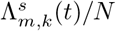 can be directly applied to compute the selected site frequency spectrum. For *t* > *T_c_*, we have that

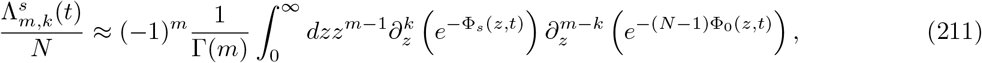

where Eq. (211) differs from Eq. (199) only by the replacement of 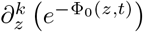 by 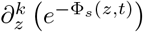. Here

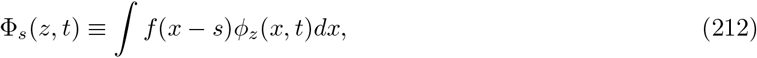

such that *e*^−Φ_*s*_(*z,t*)^ ≈ 〈*e*^−*zn*_s_(*t*)^〉; that is *e*^−Φ_*s*_(*z,t*)^ is the generating function for the (stochastic) size of a lineage seeded by a mutation effect *s*, after *t* generations. The properties of *ϕ_z_*(*x, t*) and Φ(*z,t*) discussed in Appendix A imply that Φ_*s*_(*z,t*) ≈ *e^T_c_s^*Φ_0_(*z,t*) for *s* ≪ *b*. Note that even with an additional factor of *e^T_c_s^* (compared to Φ_0_(*z,t*)) we still have that Φ_*s*_(*z,t*) ≪ 1 in the relevant region of *z*, and thus that 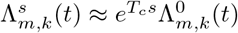.

The *t* < *T_c_* and *k* =1 case is also relevant for computing the heterozygosity. To simplify 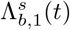 for *t* < *T_c_*, we note that *ϕ_z_*(*x, t*) has *x*-dependence *ϕ_z_*(*x, t*) ∝ *e^xt^* for the region of *x* and *z* of interest (at least until just before *t* = *T_c_*). As a result, we can make the approximation Φ_*s*_(*z,t*) ∝ *e^st^* Φ_0_(*z,t*), which yields 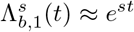. That is, an individual descends from an ancestor at time *t* (which acquired a mutation of effect *s* at that same time) with probability 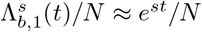.

### E.2 Partition Structure

We can see a more direct correspondence with the BSC by considering the *partition structure* implied by our model. In doing so, we will define the *partition* Π(*t*) of a sample of *m* individuals, at a time *t* into the past, as follows. The partition Π(0) consists of *m* blocks of size 1 (with each block corresponding to a different individual). As the time *t* recedes into the past, blocks of Π(*t*) merge together once their corresponding individuals share a common ancestor, with their sizes summing up (so that at time *t*, each block of Π(*t*) consists of a group of individuals related by common ancestry at *t* generations). Thus once the entire sample shares a common ancestor, Π(*t*) will consist of a single block of size *m*. For a sample of size *m*, the *partition structure* is then given by the probability *p*(*l*_1_, *l*_2_, …, *l_m_*; *t*) that the partition Π(*t*) of the sample consists of *l*_1_ blocks of size 1, *l*_2_ blocks of size 2, and so on.

To compute the partition structure *p*(*l*_1_, *l*_2_, …, *l_m_*; *t*), we first compute the coalescence configuration probabilities

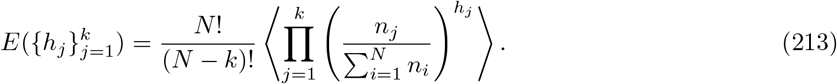

The quantity 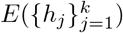 gives the probability that at time *t*, the sample traces back to a total of *k* distinct ancestors, and further, that a particular set of *h*_1_ individuals trace back to one of those ancestors, a particular set of *h*_2_ individuals trace back to another, and so on (with 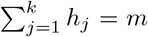). These can be calculated using manipulations similar to those used to calculate 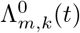; note that a similar calculation is also performed by Desai et al. (2013), though that work is primarily focused on tracing the ancestors of a sample of individuals as they move from one discrete fitness class to another by acquiring a single beneficial mutation.

The probability *E* is given by

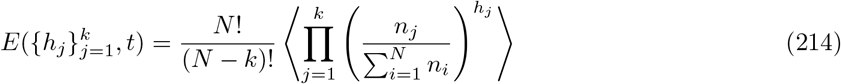

Note that the combinatorial factor *N*!/(*N* – *k*)! is included because we do not specify the common ancestors of the *k* “families” which share common ancestry. Using the identity 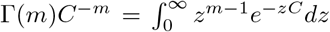 with 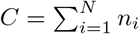, and assuming the *n_i_* are independent of one another, yields

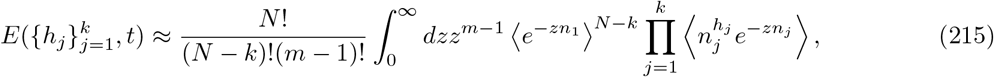

which can be expressed as

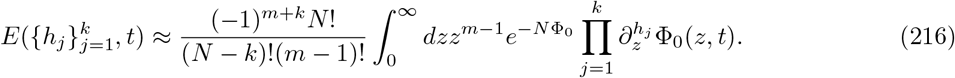

As in our computation of 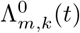, we use the properties of Φ_0_(*z,t*) discussed by Fisher (2013). For *t* < *T_c_*, linearity of Φ_0_(*z,t*) in *z* implies that *E* is negligible unless *h_j_* = 1 for all *j* (which describes the case in which each individual in the sample traces back to a distinct common ancestor). For *t* > *T_c_*, Eq. (200) can be simplified to

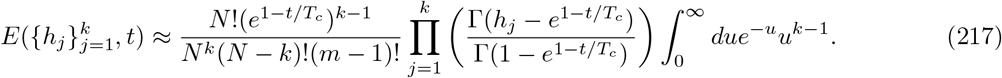

by using the Φ_0_(*z, t*) in Eq. (200) and making the substitution *u* ≡ *N*Φ_0_. The integral in Eq. (217) evaluates to Γ(*k*); using Stirling’s approximation to approximate *N*!/(*N* – *k*)! ≈ *N^k^* then yields

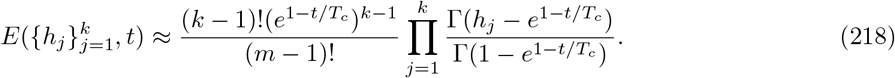

Note that *E* depends only on the *block counts l_i_*, where *l_i_* counts the number of blocks of size *i* (so that 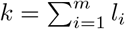 and 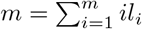). We can thus rewrite *E* as

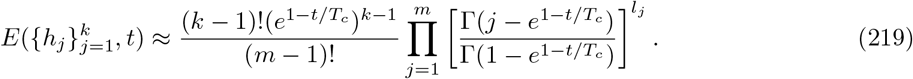

The quantity *E* gives the probability of a particular partition with block counts 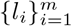. The total number of partitions with *l*_1_ blocks of size 1, *l*_2_ blocks of size 2, and so on, is 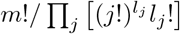 (note that by definition a partition is an *unordered* collection of blocks). The probability *p*(*l*_1_, *l*_2_, …, *l_m_*; *t*) then follows as

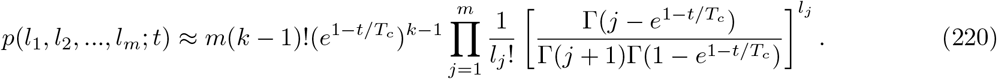

The partition structure *p*(*l*_1_, *l*_2_, …, *l_n_*; *t*) in Eq. (220) is precisely that of a partition drawn from a two parameter Poisson-Dirichlet distribution *PD*(*α, θ*) with *α* = *e*^1−*t/T_c_*^ and *θ* = 0 (Pitman, 1995; Pitman and Yor, 1997). Importantly, the partition structure of the BSC is *also* described by a Poisson-Dirichlet distribution *PD*(*α*, 0) with *α* = *e^−t^* (Berestycki, 2009). Thus, given the approximations made above, for *t* > *T_c_* the partition structure under our evolutionary model corresponds precisely to that under the BSC (up to the replacement of *t* by (*t* – *T_c_*)/*T_c_*, which reflects the delay period and coalescence timescale arising within our model).

The correspondence of the partition structure for our evolutionary model with that of the BSC does not imply that genealogies within our model are perfectly described by the BSC. For example, a further assumption of the BSC is that successive merger events are independent of one another; in our evolutionary model, heritable fitness variation among individuals may result in non-independence among merger events (though Neher and Hallatschek (2013) suggest, by a heuristic argument, that lineages within the appropriate region of fitness space equilibrate on a timescale much faster than the timescale of coalescence, such that successive merger events *are* largely independent of one another). However, the partition structure contains much more information than simply the instantaneous merger rates 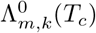 or the time-dependent probabilities *Q_k_*(*t*), and its correspondence implies a relatively strong resemblance between genealogies of our model and those of the BSC.

### E.3 The Site Frequency Spectrum

As discussed in the main text, the selected site frequency spectrum *P_s_*(*k|m*) can be computed as

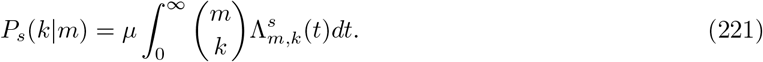

Substituting into Eq. (221) the 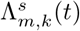 found above, we have

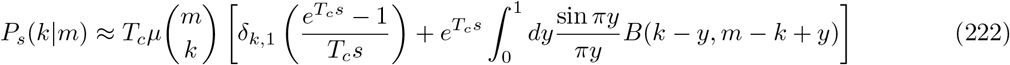

where we made the substitution *y* ≡ *e*^1−*t/T_c_*^ in the integral; note that *B*(*x*_1_, *x*_2_) denotes the Beta function which satisfies *B*(*x*_1_, *x*_2_) = Γ(*x*_1_)Γ(*x*_2_)/Γ(*x*_1_ + *x*_2_). The first term in Eq. (222) denotes a contribution from observed mutations with ages *t* < *T_c_*; consistent with our approximate result that coalescence events do not occur until *T_c_* generations into the past have elapsed, this term is only present (up to the level of our approximation) for the case *k* = 1, corresponding to singletons. Apart from an overall scale factor, the contribution from *t* > *T_c_* (the second term in Eq. (222)) exactly matches the SFS of the true BSC, as calculated by Kersting et al. (2019). Kersting et al. (2019) computes the SFS of the true BSC directly from its known partition structure; a similar calculation is carried out by Neher and Hallatschek (2013) for the common-allele portion of the BSC SFS. For completeness, here we simplify Eq. (222) for the case *s* = 0 (which gives the neutral site frequency spectrum); results for the selected site frequency spectrum follow immediately.

First, we note that *P*_0_(1|*m*) simplifies to

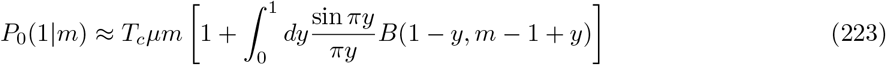

while *P*_0_(1|2), the pairwise heterozygosity, further simplifies to

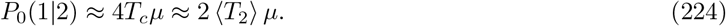

That is, we recover the well-known “molecular clock” result for *P*_0_(1|2): the average number of (neutral) pairwise differences among individuals is given by the average time since they share a common ancestor, times the per-locus mutation rate *μ*. Note that the contribution of Λ_2,1_(*t*) with *t* < *T_c_* is important for computing *P*_0_(1|2) (otherwise, our calculation would disagree with the “molecular clock” result by a factor of 2). For large *m* and *k* = 1, the contribution from *t* < *T_c_* dominates and *P*_0_(1|*m*) ~ *T_c_μm*; this is precisely the contribution of mutations which occur on the *m* terminal branches of the sample, none of which merge until *T_c_* generations into the past have elapsed.

#### The BSC SFS in the large sample size limit

To analyze the more general case with *k* > 1, it is useful to consider the large *m* limit, such that the allele frequency *ν* ≡ *k/m* can be treated as a continuous variable. The contribution from *t* < *T_c_* vanishes for large *N*, and the distribution of neutral site frequencies *h*_0_(*ν*) follows as

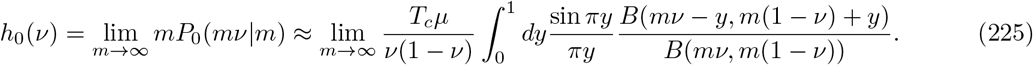

The integral on the right-hand side of Eq. (225) has been analyzed by Kersting et al. (2019); we briefly reproduce this analysis here. The right-hand side of Eq. (225) can be further simplified using Stirling’s approximation, which yields

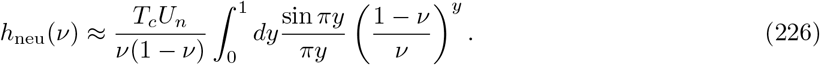

The integral in Eq. (226) can be evaluated exactly in terms of functions arctan and the exponential integral Ei, yielding

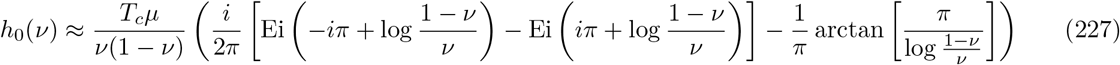

where in Eq. (227), the range of arctan must be taken as (0, *π*) instead of the usual (−*π*/2, *π*/2). For rare alleles 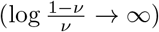 Eq. (227) simplifies to

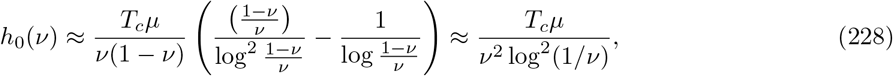

while for common alleles 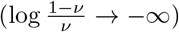,

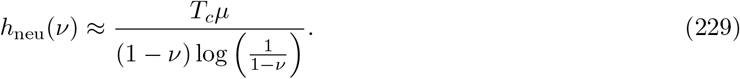

Note that the integral in Eq. (226)—and in particular, the region of *y* which dominates that integral—can be examined to identify the typical allelic ages of mutations observed at a given frequency. For example, if *ν* > 1/2, the integrand peaks at *y* = 0 and the integral is dominated by the region of width 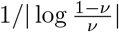 above 0. As a result, alleles at a frequency 1 – *ν* ≪ 1 have typical ages 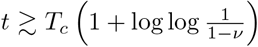. Alternatively, if *ν* < 1/2, the integrand peaks at an intermediate *y*\* between 0 and 1 which satisfies 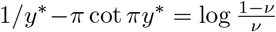. In the limit *ν* → 0, 1 – *y*\* → 1/log(1/*ν*) so that *t*\* → *T_c_*(1 + 1/log(1/*ν*)). In particular, *b*(*t*\* – *T_c_*) > 1 when log(1/*ν*) < *T_c_b*, or equivalently, when *ν* > *e^−T_c_b^*.

#### Deviations from the BSC SFS at lower frequencies

For lower frequencies such that log(1/*ν*) > *T_c_b*, the time integral yielding the SFS receives a substantial contribution from times *t* such that 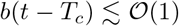. At these times the approximation 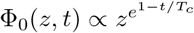 used to obtain 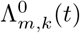, and thus the SFS, begins to break down. In Appendix A, we analyze the behavior of *ϕ_z_*(*x, t*) at intermediate times such that 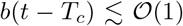, extending the analysis of *ϕ_z_*(*x, t*) carried out by Fisher (2013) at long times such that *b*(*t* – *T_c_*) ≫ 1. Because the corresponding Φ_0_(*z,t*) does not have a simple power law dependence on *z*, the manipulations carried out above to obtain the merger rates 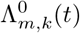, and in turn the contribution to the SFS, are difficult to extend to these times. To obtain the contribution to the SFS of these mutations, we will instead directly extract *p*(*ν*; *t*, *x*), the probability that a lineage has frequency *ν* at time *t*, given it was seeded at time 0 with relative fitness *x*, from 〈*e*^−ζ*ν*(*x, t*)^〉, the generating function for the frequency *ν*(*x, t*) of the lineage. The generating function 〈*e*^−ζ*ν*(*x, t*)^〉 can in turn be obtained from the quantity *ϕ_z_*(*x, t*) computed in Appendix A, according to

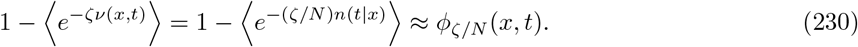

We will then compute the contribution of these mutations to the SFS by integrating *p*(*ν; t, x*) over both times *t* and initial fitnesses *x*.

We will define the function *E*[*R*; *t, x*] according to

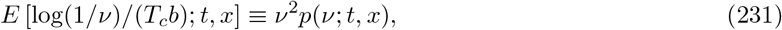

with the scaling variable *R* ≡ log(1/*ν*)/(*T_c_b*), so that

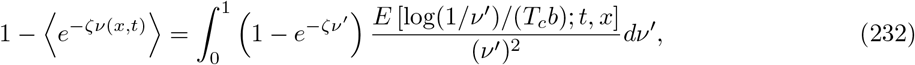

which can be written as

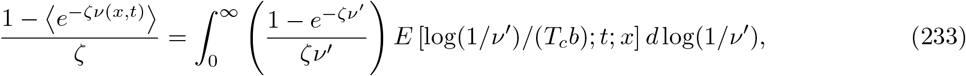

by making a change of variables. For log(1/*ν*) ≪ log *ζ*, the factor (1 – *e^−ζν^*)/(*ζν*) is exponentially suppressed, while for log(1/*ν*) ≫ log*ζ*, (1 – *e^−ζν^*)/(*ζν*) ≈ 1. Upon making the ansatz that *E*[*R*; *t, x*] varies on *R* scales of 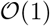 (which can later be checked to be self-consistent), we then have

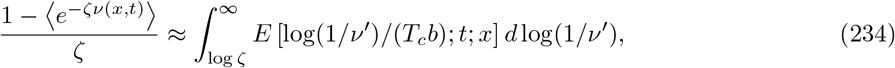

and thus

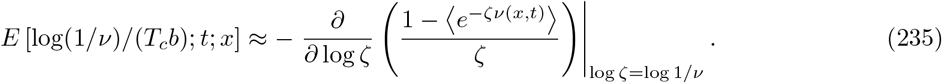

Using our result from Appendix A for *ϕ_ζ/N_*(*x, t*) when 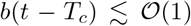, given in Eq. (135), we can evaluate the derivative on the right-hand side of Eq. (235) as a contour integral. We then have

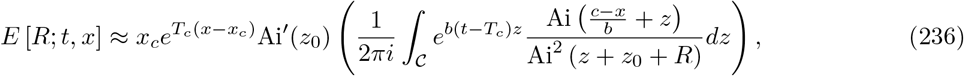

where the contour 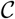 runs from *e*^−2*πi*/3^∞ through 0 to *e*^2*πi*/3^∞. To see this, note that the integral in Eq. (236) has second order poles at *z* = *z_j_* – (*z*_0_ + *R*) with corresponding residues

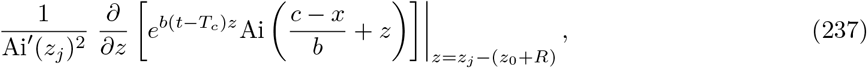

so that Eq. (236) follows from Eq. (135) and an application of the residue theorem. Integrating over times from *b*(*t – T_c_*) = −∞ to *b*(*t – T_c_*) = ∞, we have

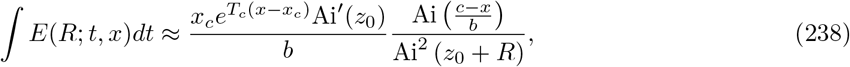

which can then be integrated against the fitness distribution to yield

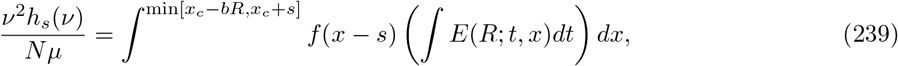

the SFS of mutations with fitness effect *s*. We can consider the cases *s* > −*bR* and *s* < −*bR* separately, in each case using the expression for *f*(*x*) given in Eq. (30). If *s* > −*bR*,

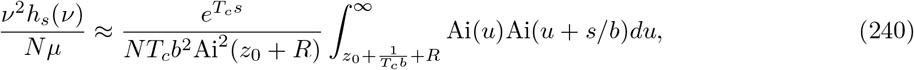

which evaluates to

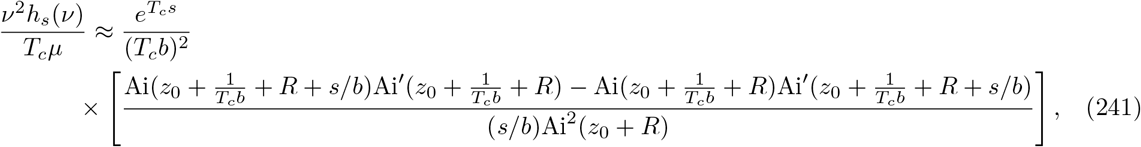

and, in the limit *s/b* → 0, to

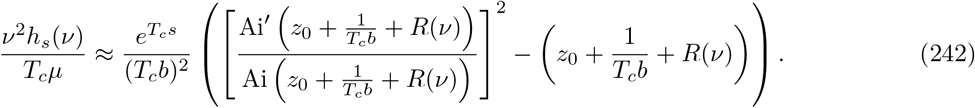

For small *R* (and large *T_c_b*), Eq. (242) simplifies to

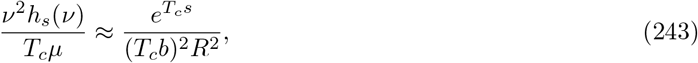

and thus

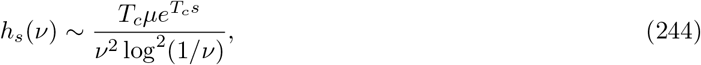

which is precisely the asymptotic behavior of the BSC SFS at low frequencies. In the limit of large *R* + *z*_0_, Eq. (242) simplifies to

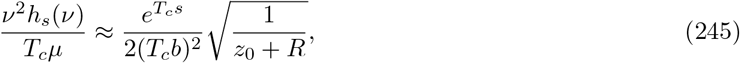

and thus

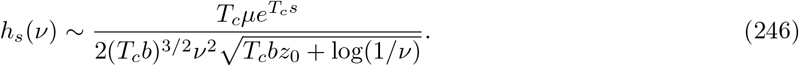

Note, however, that when *z*_0_ + *R* ≫ 1, the contribution to the SFS is dominated by lineages with initial fitnesses (*c*–*x*)/*b* ~ *z*_0_ + *R* ≫ 1; this can be seen from the integral in Eq. (240), which is boundary-dominated in this case. These fitnesses lie well below the fixation class, where the approximation 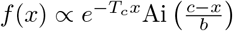, used to obtain Eq. (240), begins to break down.

If *s* < −*bR* (which requires that the mutation is deleterious) we have

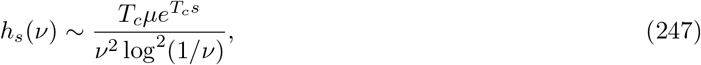

for small *R*, which again matches the low-frequency behavior of the BSC SFS.

#### The contribution to the SFS from deterministic lineage trajectories

To obtain the contribution to the SFS from mutations with ages *t* < *T_c_* (which will be important for low frequencies) we will treat the growth of lineages deterministically. For simplicity, we focus our attention on the neutral SFS *h*_0_(*ν*), which can be computed as

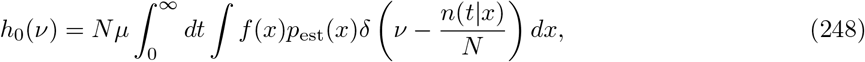

where *p*_est_(*x*) denotes the *establishment* probability of a lineage founded at relative fitness *x*, and *n*(*t*|*x*) denotes the size of a deterministically growing lineage at time *t*, given it was established at time 0 with relative fitness *x*. A solution for *n*(*t*|*x*) is reviewed, and the dynamics of deterministically growing lineage trajectories are discussed, in Appendix F (see Eq. (275)). Here we will use the result that *n*(*t*|*x*) = *n*_0_(*x*)*e*^*xt−J*(*t*)^, where 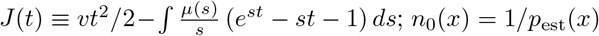 is an additional factor we include here which denotes the size of an *established* lineage—the size of a lineage at the point such that its future growth is largely described by the deterministic forces of mutation and selection, as opposed to genetic drift.

The time integral in Eq. (248) can easily be carried out, yielding

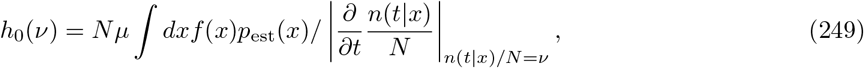

where the time derivative in Eq. (249) is evaluated at *t* such that *n*(*t*|*x*)/*N* = *ν*, for a given *x* and *ν*. To simplify Eq. (249), we note that the quantity *n*(*t*|*x*) peaks at time *t_x_* solving

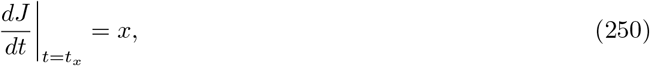

with peak size *n*_peak_(*x*) given by

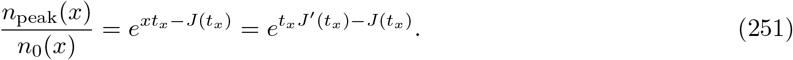

For *t* near the peak—which will turn out to often dominate the contribution of a lineage to the SFS—we have

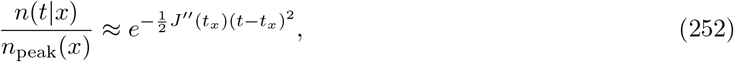

from which it follows that

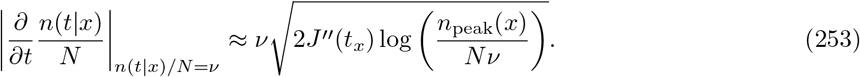

Assuming the contribution of lineages near their peak dominates the SFS, we then have

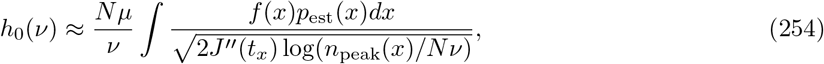

where the lower limit of integration in Eq. (254) is such that *n*_peak_(*x*) > *Nν*. A simplification occurs in that

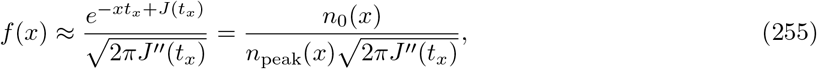

(see Eq. (186) in Appendix C) so that

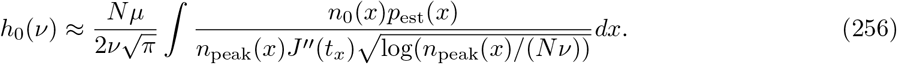

The factors *n*_0_(*x*) and *p*_est_(*x*) cancel one another in the numerator of Eq. (256), and thus contribute only logarithmic corrections to our final result. For simplicity, we therefore neglect the *x*-dependence of *n*_0_(*x*) throughout the rest of this calculation; at the end of our calculation we replace *n*_0_ with the appropriate value based on the dominant range of *x* contributing to the SFS at a given frequency. Noting that *d* log *n*_peak_(*x*) = *t_x_dx*, Eq. (256) can be rewritten as

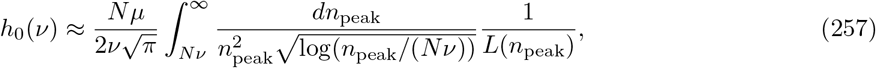

with

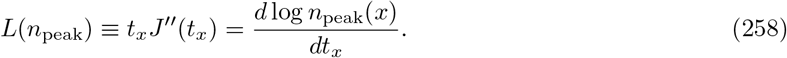

Note that in Eq. (257), and in our definition of *L*(*n*_peak_), we have replaced the dependence of *n*_peak_ on *x* with its dependence on *t_x_*. The quantity *L*(*n*_peak_) can be simplified to *t_x_*[*v* – ∫ *μ*(*s*)*dsse^st_x_^*], where *t_x_* is the time a lineage size peaks, given it peaks at a size *n*_peak_. Changing the variable of integration to *u* ≡ *n*_peak_/(*Nν*), we have

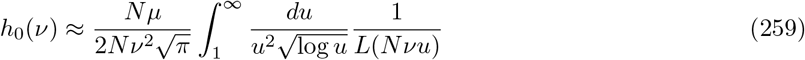

The integral over *u* is dominated by *u* close to 1 on the logarithmic scale over which *L* varies; we thus have

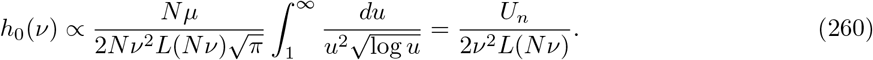

For concreteness, we now consider two special cases. First, provided that *st_x_* ≪ 1 for relevant *s, L*(*n*_peak_) ≈ *σ*^2^*t_x_* and 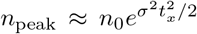, so 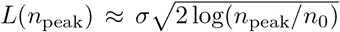. Mutations observed at these frequencies occurred in the “bulk” of the fitness distribution, with relative fitness 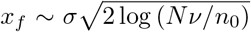; we can thus take *n*_0_ ~ 1/*σ*, to logarithmic accuracy. We thus have

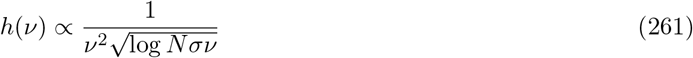

if 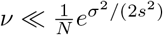 for relevant *s*..

Alternatively, if *b*(*T_c_* – *t_x_*) ≪ 1, then *L*(*n*_peak_(*x*)) ≈ 2*T_c_b*^3^(*T_c_* – *t_x_*) and *b*^2^(*t_x_* – *T_c_*)^2^ ≈ (*c* – *x*)/*b*. We then have that

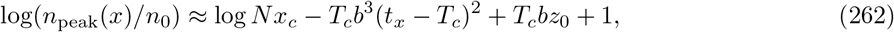

with *n*_0_ ~ 1/*x_c_*, and thus

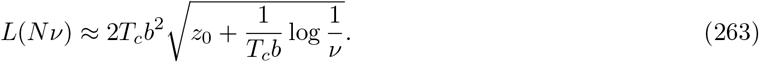

It follows that

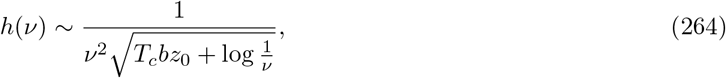

and that the dominant relative fitness *x_f_* at which observed mutations occurred is given by 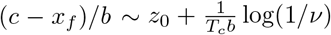. Note also that this approach breaks down if log(1/*ν*) < *T_c_b*|*z*_0_|. Mutations observed at frequencies such that log(1/*ν*) < *T_c_b*|*z*_0_| are likely to have originated at relative fitnesses *x_f_* > *c*, for which trajectories monotonically increase with time, under a deterministic approximation.

We note that the above approach resembles an approach taken by Neher and Shraiman (2011) to compute the rare-allele portion of the SFS. In that work, lineages contributing to the SFS are assumed to be founded by individuals within the Gaussian portion of the fitness distribution *f*(*x*) ~ *e*^−*x*^2^/2σ^2^^, with sizes growing deterministically according to *n*(*t*|*x*) ∝ *e*^*xt*−*σ*^2^*t*^2^/2^. We can carry out a similar calculation to obtain the SFS at very low frequencies (such that the SFS is not necessarily dominated by mutations at the peaks of their deterministic trajectories). Given these assumptions, in contrast to the previous case we can more easily carry out the *x* integral in Eq. (248) before the *t* integral, which yields

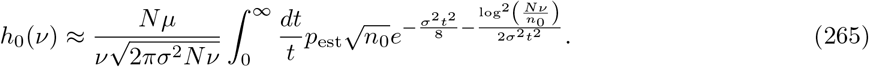

As in the above case, both *p*_est_ and *n*_0_ depend on *t* (via the *x* that yields a lineage of size *Nν* at time *t*); we will neglect this dependence and see that, under this assumption, the factor *p*_est_ in the numerator cancels with a factor 1/*n*_0_, leaving only a logarithmic dependence on *n*_0_. We can then evaluate the integral in Eq. (265) exactly, yielding

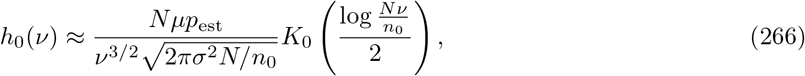

where *K*_0_ is the modified Bessel function of the second kind. Using the asymptotic expansion 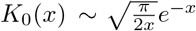 for large *x* then gives

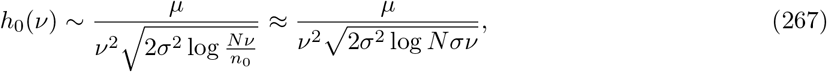

if *Nσν* ≫ 1. Note that in the final equality above we replaced *n*_0_ by 1/*σ*, since the SFS is dominated by mutations which arose at relative fitness 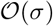 (i.e., in the “bulk” of the fitness distribution).

### E.4 Transition Density of the Effective Lead Frequency

To obtain the transition density, we first consider the generating function

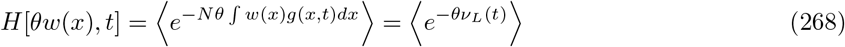

for the effective lead frequency *ν_L_*(*t*) ≡ *N* ∫ *g*(*x, t*)*w*(*x*)*dx*. Note that the effective lead frequency *ν_L_* can also be interpreted as the fixation probability of a lineage. Using the method of Appendix A, this can be done by solving Eq. (93) with the initial condition *ϕ*(*x*, 0) = *θw*(*x*). Fisher (2013) considers essentially the same quantity as in Eq. (268), with the only difference being that in that work, *g*(*x, t*) is replaced by the population-wide fitness distribution, which can fluctuate. In our case, *g*(*x, t*) is the fitness distribution of a lineage (which can fluctuate), evolving in competition with a population whose mean fitness changes at rate *v*. We can thus directly apply the following result from Eq. (68) of Fisher (2013):

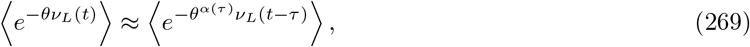

with *α*(*t*) = *e^−t/T_c_^* (in our notation). As discussed by Fisher (2013), Eq. (269) implies that *ν_L_*(*t*) is drawn from a (one-sided) Lévy stable distribution with a tail falling off as 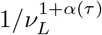. Eq. (269) is valid for *τ* > log(1/*θ*)/*x_c_* and 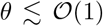. As a result, Eq. (269) does not adequately describe the probability distribution of *ν_L_* for *ν_L_* ≪ 1; our results below thus only apply to lineages with “macroscopic” effective lead frequency *ν_L_* (i.e., with a significant probability of taking over the population).

The result for 〈*e*^−*θν_L_*(*t*)^ in Eq. (269) is essentially the same as the result for a similar quantity obtained by Kosheleva and Desai (2013) (though that case involves transitions in frequencies among discrete fitness classes at discrete points in times; the analog of *α*(*τ*) also differs slightly). The derivation in Kosheleva and Desai (2013) can be carried over to our case, yielding, in our notation,

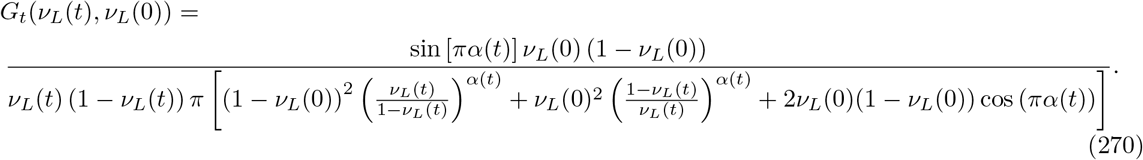

We note that a key aspect of the derivation of Eq. (270) is essential in allowing its application to our case. In particular, the calculation of Kosheleva and Desai (2013) computes Eq. (270) essentially by integrating over the transitions of *ν*_*L*,1_(*t*) (given a starting value *ν_L_*(0)) as well as *ν*_*L*,2_(*t*) (given a starting value, 1–*ν_L_*(0), equal to the complement of *ν_L_*(0)), with *ν_L_*(*t*) then equal to *ν*_*L*,1_/(*ν*_*L*,1_(*t*) + *ν*_*L*,2_(*t*)). This can be thought of as a way to explicitly enforce that the *ν_L_*(*t*) of the different lineages in a population sum up to 1, as they should in our model, and resembles the way in which our calculation of genetic diversity statistics only involves *relative* sizes of lineages. In both cases, the generating functions used correspond to Lévy distributions with diverging mean and variance, and so care must be taken to deal only with “relative” quantities in considering a population of fixed size.

## Appendix F DETERMINISTIC LINEAGE TRAJECTORIES

In this Appendix, we review the dynamics of a lineage whose fitness distribution evolves deterministically and in competition with a population steadily increasing in fitness at rate *v*. More precisely, we consider the size trajectory *n_l_*(*x*_0_, *t*) of a lineage founded at relative fitness *x*_0_ as well as its time-dependent distribution *g*(*X, t*) of fitnesses. For clarity, we proceed by writing down and directly analyzing an equation for *g*(*X, t*) with number fluctuations neglected; alternatively, the same results can be obtained using a solution for the generating function 〈*e*^−*zn*(*t*|*x*_0_)^〉 from Appendix A, which treats *n*(*t|x*_0_) as a random variable. These results facilitate a comparison of the behavior obtained within the infinitesimal and MSSM approximations (which are stochastic treatments) and the behavior obtained within a purely deterministic treatment, which is provided in the Discussion. These results are also useful in motivating an interpretation of the timescales *T_c_* and *σ*^2^/2*D* as *sweep* timescales with the infinitesimal and MSSM regimes, respectively; interpretations of *σ*^4^/4*D* and *c* are also possible.

In treating the deterministic behavior of a lineage, apart from neglecting any stochasticity in births or deaths, we assume that the deterministic growth of the lineage does not perturb the steady advance of the mean fitness of the population from its expected value *vt*. Without loss of generality, we assume a population-wide mean fitness of 0 when *t* = 0. Our focus is on the quantity *g*(*X, t*), with *g*(*X, t*)*dX* denoting the number of individuals within the lineage whose absolute fitness lies between *X* and *X* + *dX*, at a time *t* since the foundation of the lineage. The equation for *g*(*X, t*) is then given by Eq. (87) with the noise term *η_g_*(*X*) further neglected:

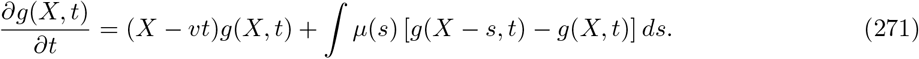

Our approach to analyzing Eq. (271) is similar to the approach carried out in Fisher (2013) to analyze the population-wide fitness distribution, assuming it evolves deterministically at fixed rate *v* (with the total population size allowed to vary in that model). We solve for the the Laplace transform 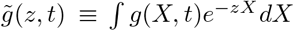, which evolves according to

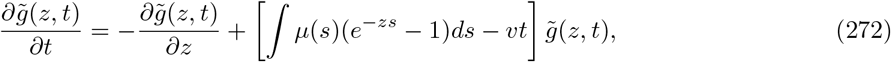

subject to the initial condition

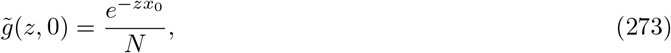

assuming the lineage is founded by a single individual with relative fitness *x*_0_. Eq. (272) can be solved using the method of characteristics, yielding

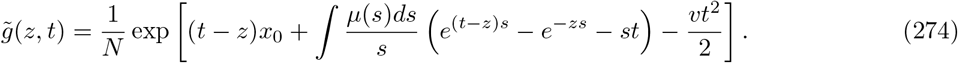

At time *t*, the total lineage size 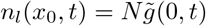 is

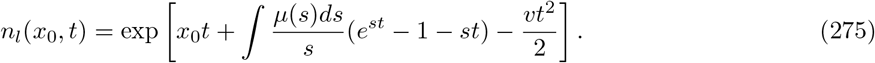

The same result for *n_l_*(*x*_0_, *t*) could be obtained using the formalism of Appendix A by substituting into 〈*e*^−*zn*(*t*|*x*)^〉 = *e*^−*ϕ_z_*(*x, t*)^ the approximate solution for *ϕ_z_*(*x, t*) obtained in Appendix E, and then evaluating *∂_z_*〈*e*^−*zn*(*t*|*x*)^〉|_*z*=0_.

The mean relative fitness 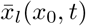 of the lineage, along with the lineage-wide fitness variance 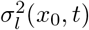, can be similarly extracted by taking *z*-derivatives of 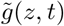:

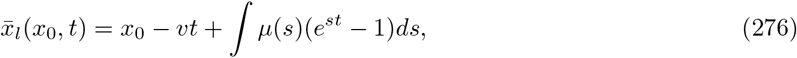

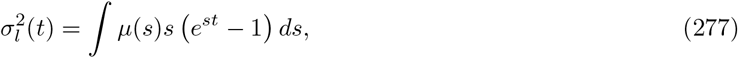

and for *p* ≥ 2,

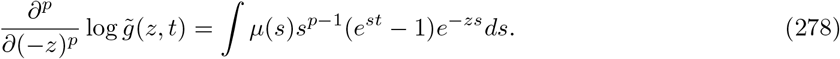

Of course, a consequence of Eq. (271) is that the lineage size *n*(*t|x*_0_) grows as 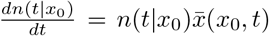. That is, the instantaneous growth rate of the lineage is equal to its mean relative fitness. Similarly, the rate of change of the mean relative fitness of the lineages matches what would be expected from Fisher’s fundamental theorem: 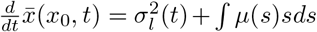. Of note is that the scale *c*, as defined in the main text, demarcates the region *x*_0_ > *c* for which lineage sizes increase in size indefinitely from the region *x*_0_ < *c* within which they do not. Lineages founded at 0 < *x*_0_ < *c* will increase initially in size but attain a maximum size at some point *t*\*, with *t*\* < *T_c_* (and with *T_c_* as defined in the main text by Eq. (17)). For *x*_0_ sufficiently close to *c*, this local maximum will be attained at 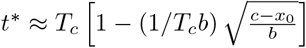. Lineages founded at *x*_0_ < 0 decrease in size indefinitely. We note also that 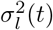 is independent of *x*_0_, and increases from 0 to match *σ*^2^, the population-wide fitness variance, over the course of precisely *T_c_* generations. For *t* > *T_c_*, 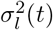 exceeds *σ*^2^, suggesting that the approximations made above break down.

The above expressions for *n_l_*(*x*_0_, *t*), 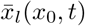 and 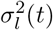 involve integrals over *ρ*(*s*). In the next subsection, we restrict our attention to the infinitesimal regime, in which the above expressions can be simplified in terms of *σ*^2^ and *D*. We then turn to the MSSM regime, in which simplifications can be made in terms of the scales *T_c_*, *b* and *c*.

### F.1 Lineage Trajectories in the Infinitesimal Regime

Assuming validity of the infinitesimal approximation, for 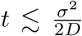, Taylor approximations can be made within Eq. (275), Eq. (276) and Eq. (277), yielding

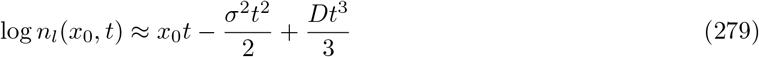

with the quantities *σ*^2^ and *D* as defined in the main text (Neher et al., 2014b). The mean and variance in fitness of the lineage-wide fitness distribution can be computed in a similar way, yielding

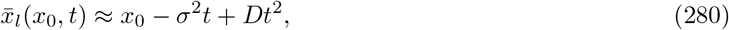

and

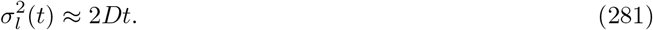

Under the additional assumption 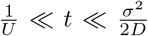, the (relative) fitness density *g*(*x, t*) of the lineage can be approximated as

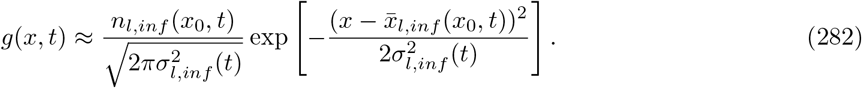

for 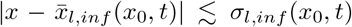. This can be done by writing down *g*(*X, t*) as the inverse Laplace transform of Eq. (274), identifying the regions dominating the integral (saddle points, which are located at 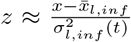), and arguing that the relevant terms can be neglected within those regions, provided that *Ut* ≫ 1. Alternatively, we can note that Eq. (282) solves

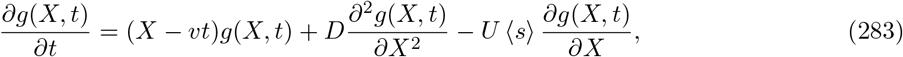

which can be obtained from (271) by a Taylor approximation of *g*(*X – s, t*) to second order in *s* (similar to the expansion of *f*(*x* – *s*) in the usual presentation of the infinitesimal approximation). Within the region 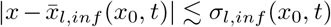 of interest, *g*(*X, t*) varies on a scale of *σ_l_*(*t*), suggesting validity of this Taylor approximation when *s* ≪ *σ_l_*(*t*) throughout the support of *ρ*(*s*), or equivalently, when *Ut* ≫ 1. Eq. (283) motivates the following description of the deterministic dynamics of lineages within the infinitesimal regime: lineages grow in size at at a rate given by their overall mean (relative) fitness advantage, advect upwards in absolute fitness at rate *U*〈*s*〉, and diffuse outward in fitness with diffusion constant *D*.

The above analysis considers the behavior of a single lineage evolving deterministically, given its initial fitness *x*_0_. We can easily extend our analysis to consider how, under the deterministic approximation, the collective set of descendants of a *class* of individuals evolves. In particular, given the results from the main text for the steady-state relative fitness distribution *f*(*x*), we can compute the number *n*(*t*; *x*_1_, *x*_2_) of descendants of the class of individuals with fitnesses between *x*_1_ and *x*_2_, after *t* generations. At the particular time *t* = *σ*^2^/2*D*, *n*(*t*; *x*_1_, *x*_2_) simplifies to

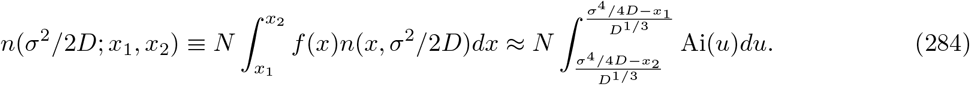

As we discuss in the main text, within the infinitesimal regime, the region *σ*^4^/4*D* < *x* < *x_c_* contributes a future common ancestor of the population with probability 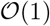 (which motivates us to refer to this region as the fixation class). Under the deterministic approximation, after *σ*^2^/2*D* generations, the lineage seeded collectively by the fixation class has reached a size of

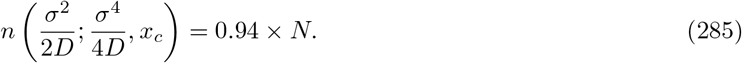

individuals. This in turn motivates our interpretation of *σ*^2^/2*D* as a *sweep* timescale: over the course of *σ*^2^/2*D* generations, the fixation class collectively sweeps through to comprise a macroscopic fraction of the population, under the deterministic approximation. A timescale *σ*^2^/2*D* has also been interpreted as a sweep timescale by

As discussed in the main text, *D*^1/3^ corresponds to the width of the fixation class. The above results results further suggest an interpretation of *D*^1/3^ as a *diffusion* scale. To see this, we consider the ratio of lineage sizes seeded simultaneously with initial fitnesses *x*_0_ and *x*_1_ > *x*_0_, which evaluates to *n*(*t|x*_1_)/*n*(*t|x*_0_) = *e*^(*x*_1_–*x*_0_)*t*^. The lineage seeded at initial fitness *x*_1_ is favored by selection on the timescale *τ_sel_* = 1/(*x*_1_ – *x*_0_). Over this same timescale, both lineages acquire fitness variance of the amount 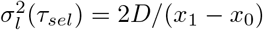. The lineage-wide fitness distributions overlap substantially at times *before t* = *τ_sel_* only if 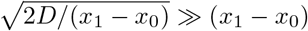, or equivalently, only if (*x*_1_ – *x*_0_) ≪ *D*^1/3^. The fitness scale *D*^1/3^ then corresponds roughly to the maximum fitness difference of two lineages such that their fitness distributions overlap substantially *before* selection acts to amplify the frequency of the more-fit lineage. The overlap of the fitness distributions can be thought of as resulting from *diffusion* of the lineages through fitness space; this diffusion is particularly important on fitness scales smaller than *D*^1/3^.

### F.2 Lineage Trajectories in the MSSM regime

Within the MSSM regime, analogous approximations can be made for log *n*(*t|x*_0_) and its derivatives. Here, however, a Taylor approximation of *e^st^* for *t* < *T_c_* is not necessarily justified, since *T_c_s* is not necessarily small throughout the bulk of *μ*(*s*). In fact, even the smaller quantity *s/b* need not be small throughout the bulk of *μ*(*s*); *s/b* is only required to be small within the region dominating ∫ *μ*(*s*)*e^T_c_s^ ds*. For deleterious mutations, the condition that *s/b* ≪ 1 for *s* dominating ∫ *μ*(*s*)*e^T_c_s^ ds* is strictly *weaker* than the condition that *s/b* ≪ 1 for *s* dominating ∫ *μ*(*s*)*ds*.) Instead, given that *s* ≪ *b* for *s* dominating ∫ *μ*(*s*)*e^T_c_s^ ds*, an approximation of Eq. (275) can be made for 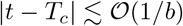, yielding

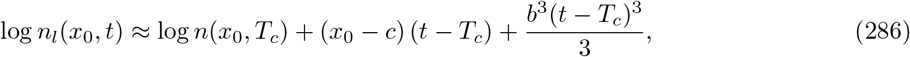

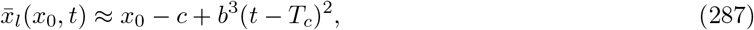

and

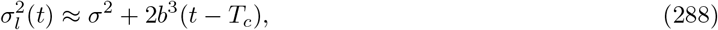

with

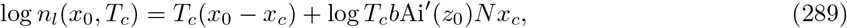

where the last equality makes use of Eq. (31) from the main text.

As in the previous subsection, the quantity *n*(*T_c_*; *x*_1_, *x*_2_) can be evaluated, in this case by making use of Eq. (275) along with Eq. (179) for *f*(*x*) obtained within the MSSM regime. Since Eq. (179) is accurate only for *x* within 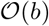 of *c*, *x*_1_ and *x*_2_ must also be within 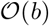 of *c* to yield reasonable accuracy in *n*(*T_c_*; *x*_1_, *x*_2_). Provided this condition is met,

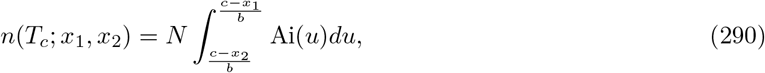

and in particular,

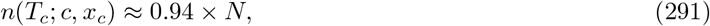

such that *T_c_* can be interpreted as a *sweep* timescale, in the sense that the fixation class collectively sweeps to comprise a macroscopic fraction of the population over *T_c_* generations. The timescale 1/*b* can be interpreted as the *rate* at which *n*(*T_c_*; *c*, *x_c_*) approaches *N*; to see this, we note that for *c* < *x*_0_ < *x_c_* and 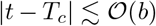, the quantity *n_l_*(*x*_0_, *t*) varies on the timescale 1/*b*. The fitness scale *b* also corresponds to the width of the fixation clas; however, the interpretation given in the previous subsection to *D*^1/3^ as a diffusion fitness scale does not extend to this case.

## Appendix G PARTICULAR DISTRIBUTIONS OF FITNESS EFFECTS

In this Appendix, we consider two classes of DFEs: stretched-exponential distributions, of the form

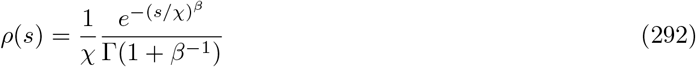

considered by Desai and Fisher (2007), Good et al. (2012) and others, as well as (more briefly) gamma distributions, of the form

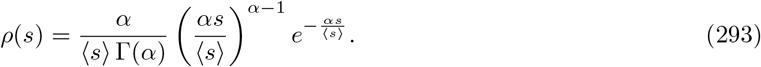

For these classes of DFEs, we simplify many of the expressions provided in the main text, focusing particular attention on the scale 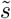 and its magnitude relative to the scales 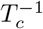 and *b*. In doing so, our primary goal is to evaluate the suitability of the infinitesimal and MSSM approximationes, particularly in the *N* → ∞ limit. We limit our attention here to DFEs consisting only of beneficial mutations, although our analysis can easily be extended to consider DFEs consisting of deleterious mutations or a combination of the two.

We begin by considering the case of an exponential DFE. Attention on the case of a beneficial exponential DFE can be motivated by arguments from extreme value theory (Gillespie, 1984; Orr, 2003). We then briefly consider more general gamma distributions, and conclude by considering the case of a stretched-exponential DFE with *β* > 1. The requirement that *β* > 1 is a key requirement for validity of our approach; otherwise, the integrals *M_p_*(*T_c_*), which we consider throughout, do not converge.

### G.1 Beneficial Exponential DFE

The special case in which mutations are beneficial and exponentially-distributed yields particularly simple analytical results. We assume in this Subsection that 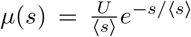. We will compute quantities of interest by first computing the moments *M_p_*(*T_c_*), defined as *M_p_*(*T_c_*) ≡ ∫ *μ*(*s*)*e^T_c_s^ s^p^ ds* for *p* ≥ 1, *M*_0_(*T_c_*) ≡ ∫ *μ*(*s*)(*e^T_c_s^* – 1) *ds*, and 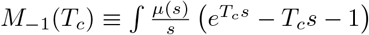. These evaluate to

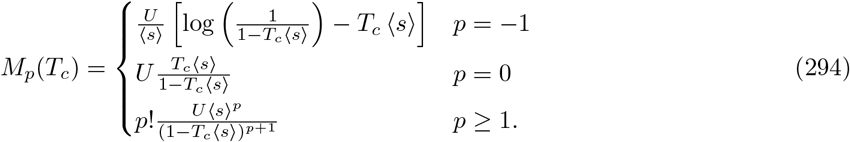

In particular,

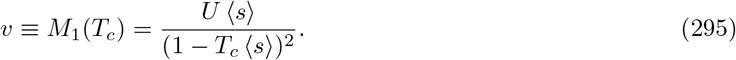

Consequently, *T_c_*〈*s*〉 approaches 1 from below as *N* → ∞ (note this follows from the fact that *v* increases monotonically, without bound, with *N*). The fact that *T_c_*〈*s*〉 < 1 ensures convergence of the *M_p_*(*T_c_*) for all *p*, and implies that 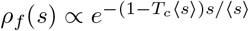 is a strictly decreasing function of *s* on the interval (0, ∞). The average effect size 〈*s_f_*〉 of a fixed mutation simplifies to

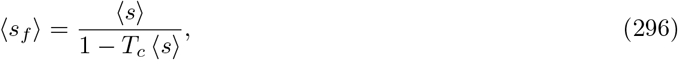

such that

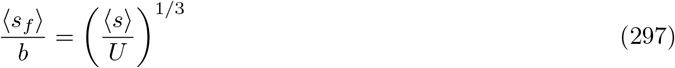

is independent of *N*. Since *ρ_f_*(*s*) is an exponential distribution, the standard deviation Δ*s_f_* of fixed fitness effects equates to 〈*s_f_*〉, and thus 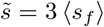 given the definition of 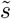 in Section II. Additionally,

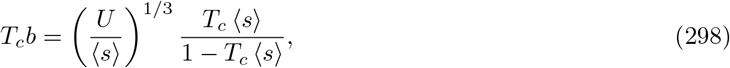

so that the conditions 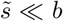 and *T_c_b* ≫ 1 are both met (implying suitability of our MSSM approximation) in the *N* → ∞ limit, as long as 〈*s*〉 ≪ *U*.

Conditions of validity can be considered more precisely by considering the quantities *S_v_*(1) and 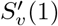 discussed in Appendix C. These quantities evaluate to

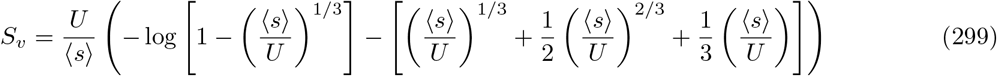

and

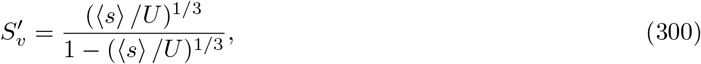

with both *S_v_* and 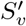 tending to 0 as 〈*s*〉 /*U* → 0, and diverging for 〈*s*〉 ≥ *U*. We note that in the limit 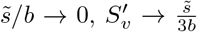 (which is the same result for 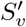 obtained when *ρ*(*s*) is *not* an exponential distribution, but *ρ_f_*(*s*) is sufficiently sharply peaked—as in the case *ρ*(*s*) = *δ*(*s* – *s_b_*), for instance). This can be seen as a motivation for taking the particular definition 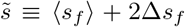; with this definition, the (arguably) simpler and more interpretable quantity 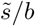 can be used instead of 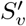 in determining the suitability of the MSSM approximation. The quantity 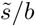 behaves the same way as 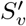 in the opposing limiting cases that *ρ_f_*(*s*) is very sharply peaked, and that *ρ_f_*(*s*) is an exponential distribution. The same holds for the quantity *S_v_*; 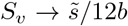 in the limit 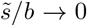, both when *ρ_f_*(*s*) is sharply peaked and when *ρ_f_*(*s*) is an exponential distribution.

#### An asymptotic approximation for the *N* → ∞ limit

We can obtain asymptotic results in the limit *N* → ∞ at fixed 〈*s*〉 /*U* ≪ 1. As noted above, in this limit *T_c_*〈*s*〉 → 1, so this limit can alternatively be thought of as the limit *T_c_b* → ∞ at fixed 〈*s*〉 /*U* ≪ 1. We wish to obtain the leading behavior of *T_c_b* in terms of *N* by analyzing Eq. (31) in this limit. In doing so, it is useful to simplify Eq. (31) in terms of the large parameter *y* ≡ (*U*/ 〈*s*〉)^1/3^*T_c_b*. Using the integrals evaluated in Eq. (294), we have, in the limit *T_c_* 〈*s*〉 → 1,

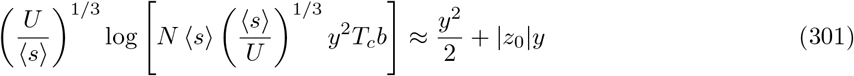

which can be further approximated as

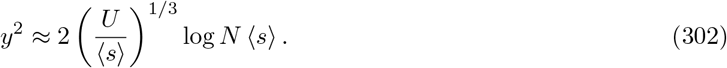

if *y* ≫ 1. This yields

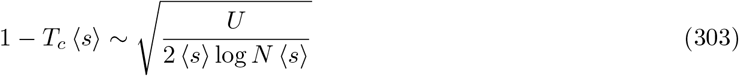

and therefore

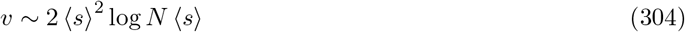

which is approximately valid if log *N*〈*s*〉 ≫ *U*/ 〈*s*〉. Note that for *U*/ 〈*s*〉 ≫ 1, an unreasonably large population size *N* may be required for this asymptotic approximation to be useful (and at the same time, *U*/ 〈*s*〉 ≫ 1 is required for validity of the MSSM approximation). In particular, caution must be taken in interpreting Eq. (303) or Eq. (304) as a function of *U* or 〈*s*〉, since changes in either quantity can cause the asymptotic approximation to break down. In practice, the asymptotic approximation *v* ~ 2(*U* 〈*s*〉^2^)^1/3^ (*s*) log *N* 〈*s*〉 generally provides better accuracy throughout the parameter regime we have considered in simulations. Additionally, in this limit

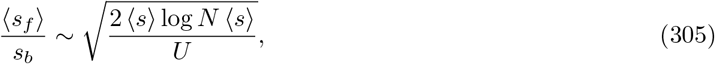

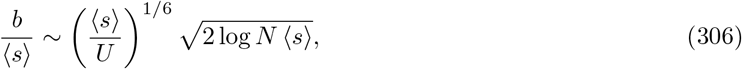

and

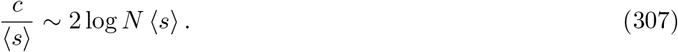

#### Single-s equivalence

Finally, we note that in the limit *N* → ∞ the dynamics can be captured an effective mutational fitness spectrum *μ_eff_*(*s*) = *U_eff_δ*(*s* – *s_eff_*), with

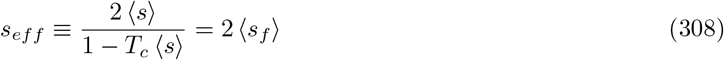

and

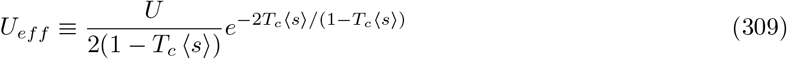

With these definitions of *U_eff_* and *s_eff_*, 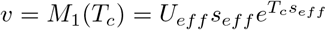 and 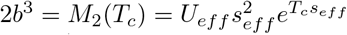, matching the corresponding quantities that are obtained for *μ*(*s*) = *U_eff_ δ*(*s* – *s_eff_*). In the limit *N* → ∞, the quantities *T_c_* and *v* depend on *μ*(*s*) only through *M*_1_(*T_c_*) and *M*_2_(*T_c_*); the dynamics can thus be described by the effective mutation rate *U_eff_* and effective DFE *δ*(*s* – *s_eff_*).

### G.2 Beneficial Gamma DFE

Much of the above analysis can be straightforwardly generalized to the case of gamma-distributed beneficial fitness effects, with

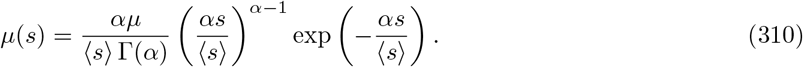

Here 〈*s*〉 denotes the mean effect size and *α* > 0 denotes the shape parameter of the gamma distribution (such that the variance in effect sizes is 〈*s*〉^2^ /*α*).

The moments *M_p_*(*T_c_*) evaluate to

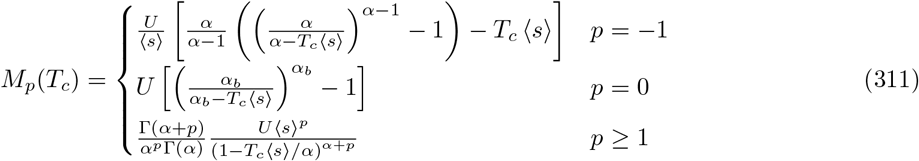

with *v* = *M*_1_(*T_c_*) → ∞ as *T_c_*〈*s*〉 → *α*; consequently, *T_c_* 〈*s*〉 approaches *α* from below as *N* → ∞. (The expression for *M*_−1_(*T_c_*) assumes *α* ≠ 1.)

The distribution of fixed effects in the MSSM regime is just *ρ_f_*(*s*) ∝ *ρ*(*s*)*e^T_c_s^*, which is a gamma distribution with shape parameter *α* and mean effect 〈*s_f_*〉 = 〈*s*〉 /(1 – *T_c_* 〈*s*〉 /*α*); the variance in fixed effects is given by 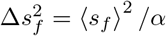. The ratio 〈*s_f_*〉 /*b* simplifies to

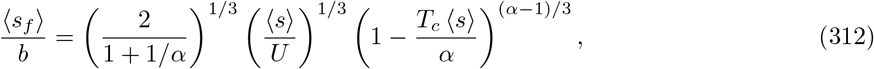

and so as long as *α* > 1, 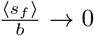 (and thus 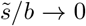) in the *N* → ∞ limit. Further, *T_c_b* → ∞ as *N* → ∞, suggesting that the MSSM regime is approached in the *N* → ∞ limit as long as *α* > 1. In the opposite case *α* < 1, the MSSM approximation is expected to break down as *N* → ∞ since 〈*s_f_*〉 /*b* → ∞. Note that this can be confirmed by simplifying the more precise conditions of validity 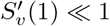 and *S_v_*(1) ≪ 1. Additionally, as might be expected, the infinitesimal approximation breaks down in the limit *N* → ∞.

### G.3 Beneficial Stretched-Exponential DFE

Here we consider beneficial stretched-exponential DFEs of the form

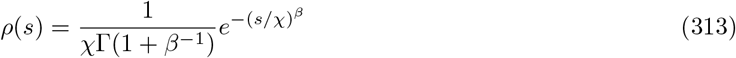

for *s* > 0, with *β* > 1. To do so involves evaluating integrals of the form 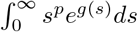, with 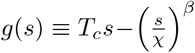. In contrast to the *β* = 1 case, simple closed-form expressions for these integrals are not available when *β* > 1. In the limit that *T_c_χ* → 0, these integrals can be approximated as

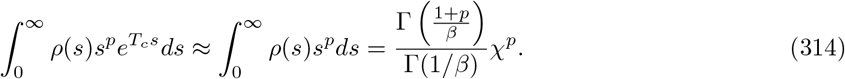

We focus here instead on the case in which *T_c_χ* ≫ 1—which is obtained in the limit *N* → ∞—in which these integrals and related expressions can be approximated using Laplace’s method.

The quantity *g*(*s*) peaks at *s*\* satisfying

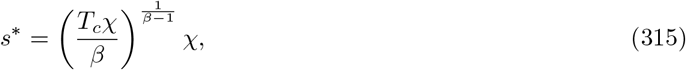

with

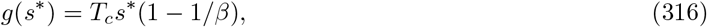

and

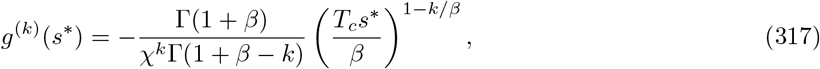

for *k* = 2. The width Δ ≡ |*g*^(2)^(*s*\*)|^1/2^ of *e*^*g*(*s*)^ scales, relative to *s*\*, as

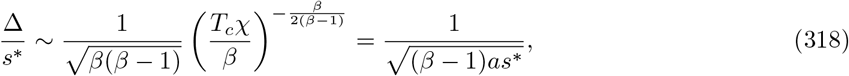

and

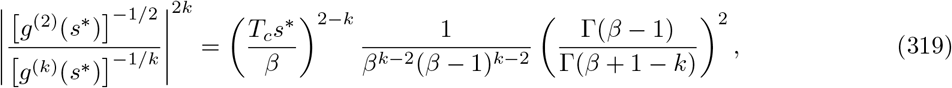

motivating the use of Laplace’s method when *T_c_s*\* ≫ *β* and *β* – 1 ≫ 1. Laplace’s method yields

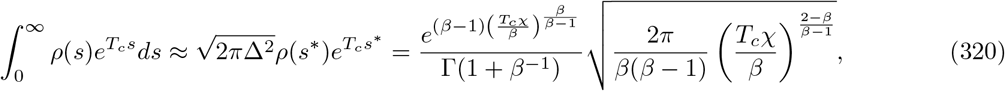

and more generally,

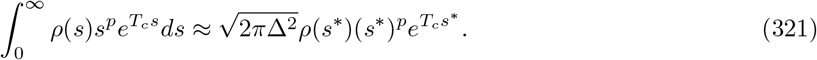

Under this application of Laplace’s method, 〈*s_f_*〉 is approximated to precisely *s*\*. Further, this application of Laplace’s method essentially implies a *single-s equivalence* of the type discussed in Good et al. (2012), whereby the dynamics arising from the full *μ*(*s*) can be approximated by the dynamics arising within a population subject to mutations which confer only a single effect *s_eff_* = *s*\*, occurring at rate 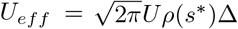.

The magnitude of *s*\* can be compared to *b*, with the result

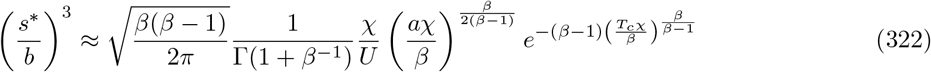

From Eq. (322) it follows that *s*/b* decreases with *T_c_* (and therefore with *N*) at large *T_c_* (and therefore at large *N*). Together with the fact that *T_c_b* increases monotonically with *N* without bound, this suggests that the MSSM regime is approached in the *N* → ∞ limit. In contrast with the case of exponentially-distributed beneficial fitness effects (*β* = 1), no requirement needs to be made that 〈*s*〉 ≪ *U*.

## Appendix H SIMULATION METHODS

In this Subsection, we provide details on the implementation of our Wright-Fisher simulations. In conducting individual-based simulations, we track the absolute fitness of, as well as the mutations carried by, each individual over the duration of the simulation. The population is initialized as a clonal population with initial absolute fitness 0. Each generation consists of a mutation step and and a stochastic birth/death step. In the mutation step, each individual acquires a Poisson-distributed number of mutations (with mean *U*), with the effect sizes of those mutations drawn from *ρ*(*s*) and incrementing the log-fitness of an individual by *s*. Individuals are also subject to completely neutral mutations at rate *U_n_* = 0.1 (per-individual, per-generation). In the stochastic birth/death step, *N* individuals are sampled with replacement from the population, with an individual’s sampling probability proportional to its absolute exponential (as opposed to log) fitness.

Our simulations implement an “infinite-sites” model of mutation, in which each new mutation is assigned a unique identity upon its occurrence. Each time a mutation occurs, we record its fitness effect. When a mutation is shared by all individuals in the population, we mark the mutation as fixed. This enables us to empirically measure the distribution of fixed fitness effects, which can be compared to our theoretical predictions *ρ_f_*(*s*). Additionally, we measure the time elapsed before the *first* fixation of a neutral mutation, which we take as the length *T_eq_* of an epoch. From that point on, we record the state of the population each epoch until the conclusion of the simulation. In particular, at each epoch, we record the mean fitness of the population (and in some cases a histogram of relative fitnesses present in the population), the mutations which have fixed (and their fitness effects), as well as a measurement of the population’s site frequency spectra, both of neutral mutations and of selected mutations. From measured site frequency spectra we obtain measurements of the pairwise heterozygosity, both of neutral mutations and of selected mutations. We discard the first 10 recorded states of the population, which may still reflect the transient behavior of the population after its initialization as a clonal population. The code implementing our simulations is available upon request.

## Appendix I NUMERICAL METHODS

To obtain theoretical predictions for comparison with simulation results, we numerically solve the system

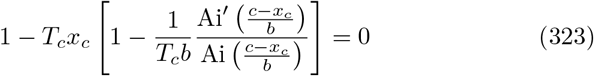

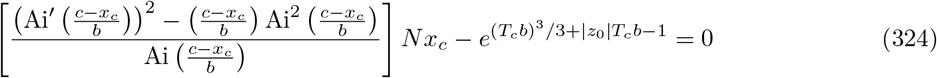

within the infinitesimal approximation; in the MSSM approximation, we solve Eq. (323) along with

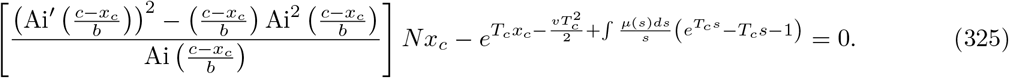

In doing so, the definitions for *T_c_, b* and *c* differ between the two approaches; in the infinitesimal approximation,

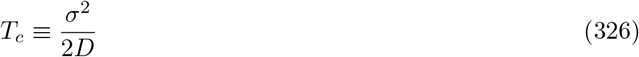

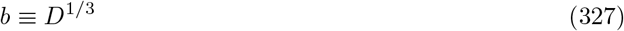

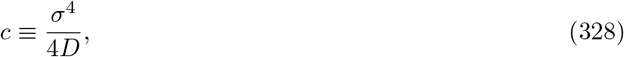

with *σ*^2^ and *D*^1/3^ defined by *v* ≡ *σ*^2^ + *U*(*s*) and 2*D* ≡ *U*〈*s*^2^〉, respectively. In contrast, within the MSSM approximation,

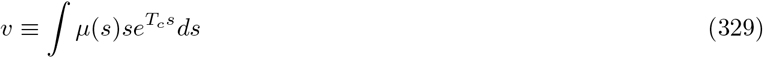

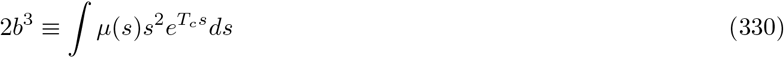

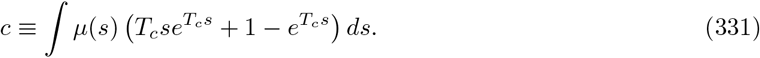

The parameters *N* and *μ*(*s*) can be thought to be the “known” inputs to a system of two equations with the two unknowns *T_c_* and *x_c_*.

Numerical solution is made more straightforward by enforcing upon the solution that *T_c_* > 0 (to ensure that *v* > ∫ *μ*(*s*)*sds*) and *T_c_* < *N* (given the interpretation of *T_c_* as a coalescence timescale, necessarily shorter than the timescale *N* of neutral pairwise coalescence). Additionally, we enforce that *x_c_* > 0 (since otherwise, our approach would neglect the fitness density of all individuals with greater than average fitness) and *x_c_* < *c* – *bz*_0_ (since otherwise, our approach would predict negative values of fitness density for individuals with relative fitness exceeding *c* – *bz*_0_).

To solve the relevant systems, an array of test *T_c_* values, logarithmically spaced between 1 and *N*, are considered. For each *T_c_* value, the corresponding *x_c_* solving Eq. (323) is identified. This is done by evaluating the left-hand side of Eq. (323) for linearly spaced *x_c_* between 0 and *c* – *bz*_0_, identifying zero crossings and employing the scipy function *brentq*. In cases where multiple solutions *x_c_* might be obtained, the solution closest to *c* – *bz*_0_ is taken. This permits evaluation of the left-hand side of Eq. (324) or Eq. (325) for a particular *T_c_*. Once this is done for the entire array of test *T_c_* values, zero crossings are identified and the scipy function *brentq* is employed. In cases where no zero crossings are identified, the scipy function *fsolve* is employed. The initial guess is taken as the test *T_c_* value which minimizes the absolute value of the left-hand side of Eq. (324) or Eq. (325). In cases where multiple zero crossings are identified, the scipy function *brentq* is employed for each zero crossing, and the solution with the largest value *T_c_b* is taken.

In practice, our simulations are carried out for parameter combinations lying on a grid with specified *T_c_b* and *T_c_* 〈*s*〉 (or, equivalently, *T_c_* 〈*s_f_*〉) values. We thus invert the above equations to solve for a corresponding *N* and *μ*(*s*), given *T_c_b* and *T_c_* 〈*s*〉. A given combination of *T_c_b* and *T_c_* 〈*s*〉 values is compatible with a range of *N* values (with *N*, *T_c_b* and *T_c_* 〈*s*〉 fully determining *μ*(*s*); note that properly scaled dynamical quantities depend on *N*, *U* and 〈*s*〉 only via the scaled quantities *NU* and *N* 〈*s*〉.) We thus choose a value of *N* that ensures feasibility of our individual-based simulations.

---

**FIG. S1.**
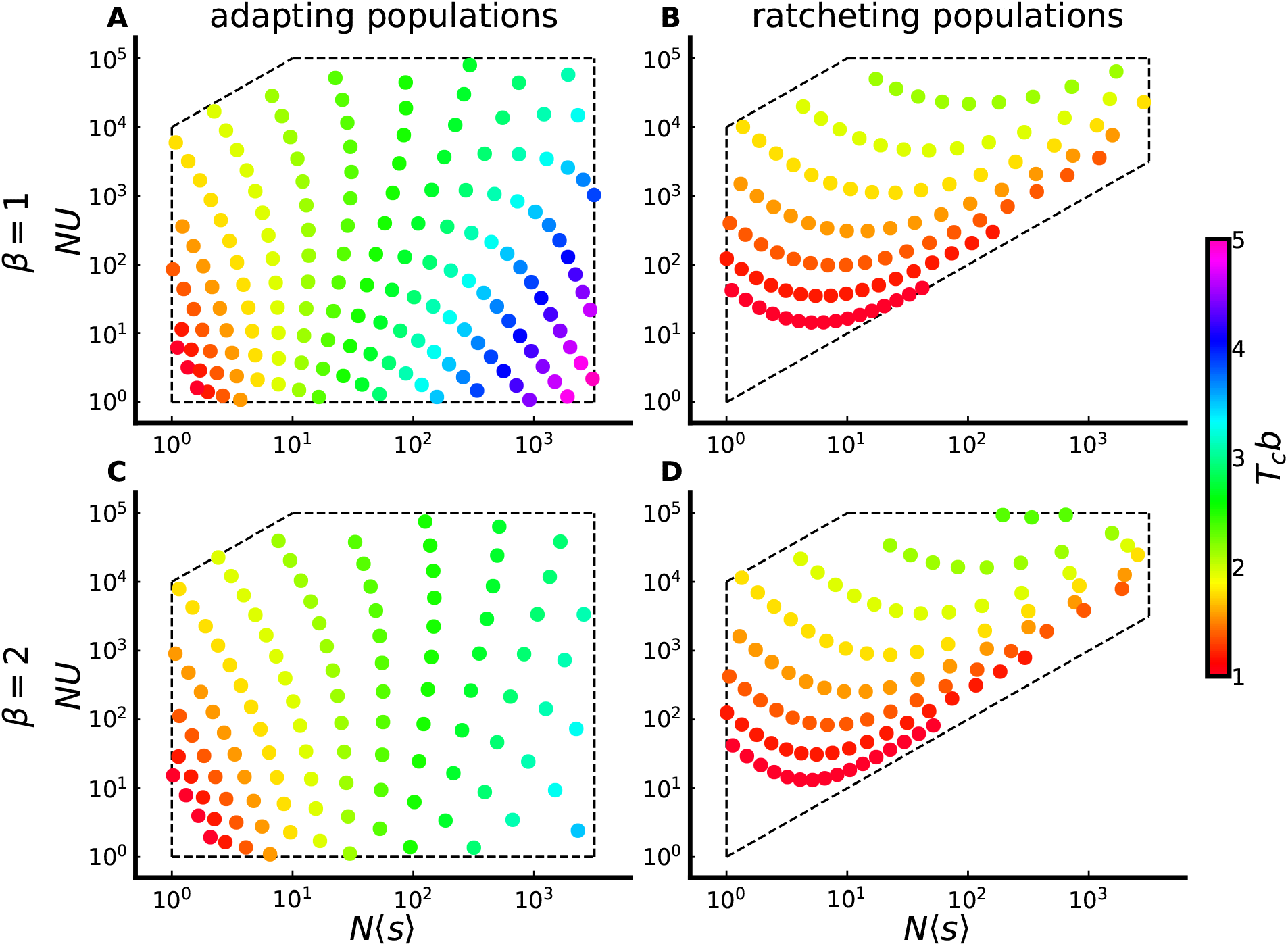
Simulated parameter combinations with an exponential or *β* = 2 stretched exponential DFE, plotted as a function of *NU* and *N*〈*s*〉, and colored by values of *T_c_b*. Dashed lines represent constraints imposed on a grid of parameter combinations with linearly spaced *T_c_b* values, and logarithmically spaced *T_c_*〈*s_f_*〉 values (for adapting populations) or *T_c_*〈*s*〉 values (for ratcheting populations). These constraints are chosen to limit attention to the MSSM regime (and the region of parameter space just beyond its regime of validity) and to ensure feasibility of individual-based simulations.

**FIG. S2.**
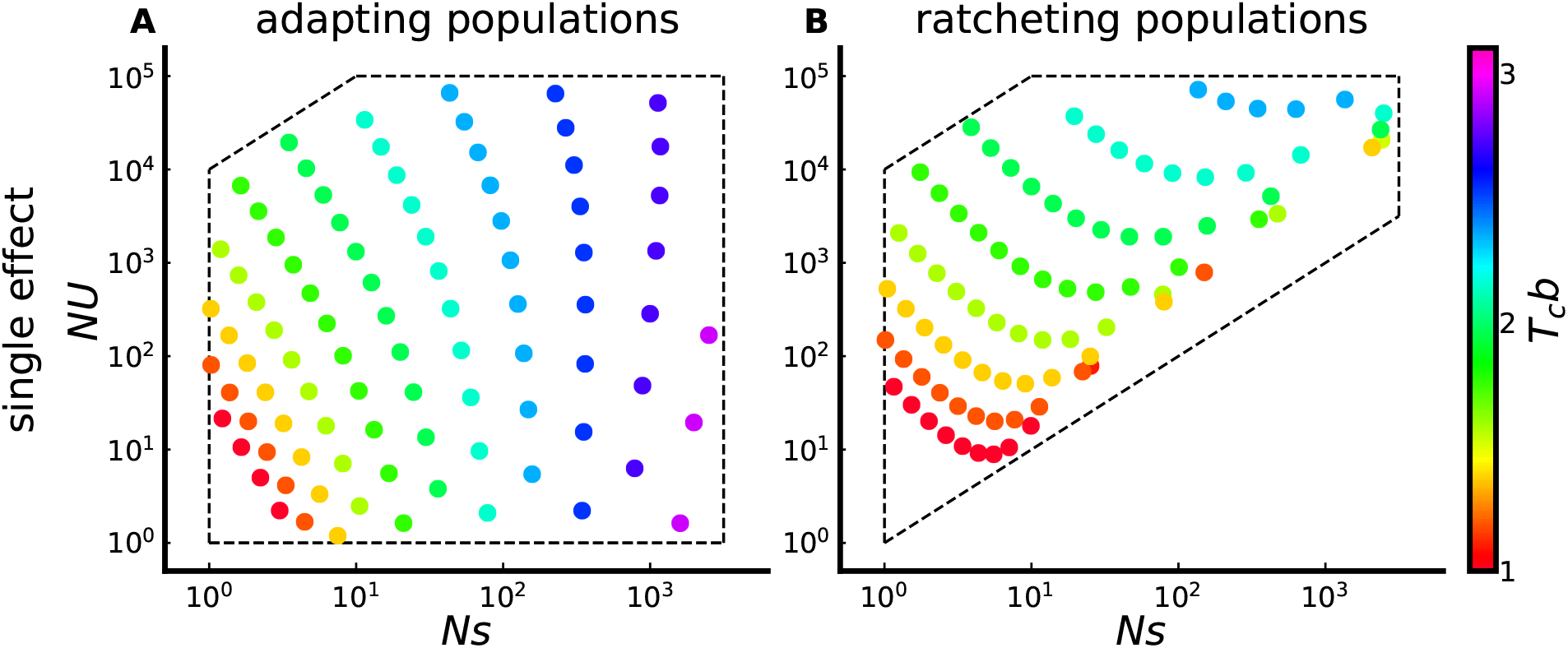
Simulated parameter combinations with a single-effect DFE, plotted as a function of *NU* and *Ns*, and colored by values of *T_c_b*. As in Fig. S1, dashed lines represent additional constraints imposed on the grid of parameter combinations.

**FIG. S3.**
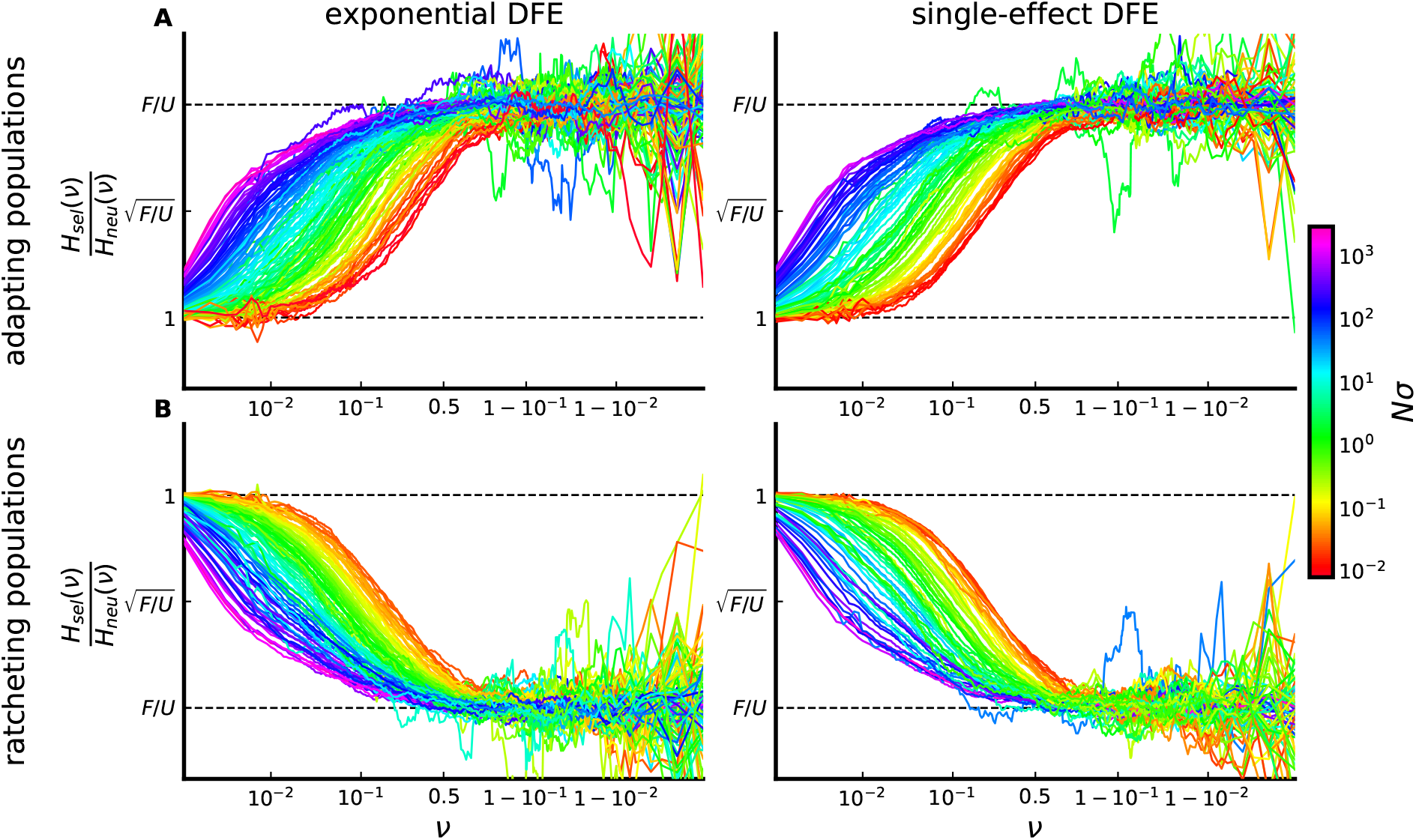
Color plot of SFS ratio curves for adapting populations (A) and ratcheting populations (B). SFS ratio curves for the entire range of simulated *T_c_*〈*s*〉 values are plotted with the same scaling of the vertical axis as in Fig. 5, colored by *Nσ* values. Displayed curves lie on the constrained grids for exponential DFEs (left) or single-effect DFEs (right), with the additional restriction that 〈*s_f_*〉 < 3*b*.

**FIG. S4.**
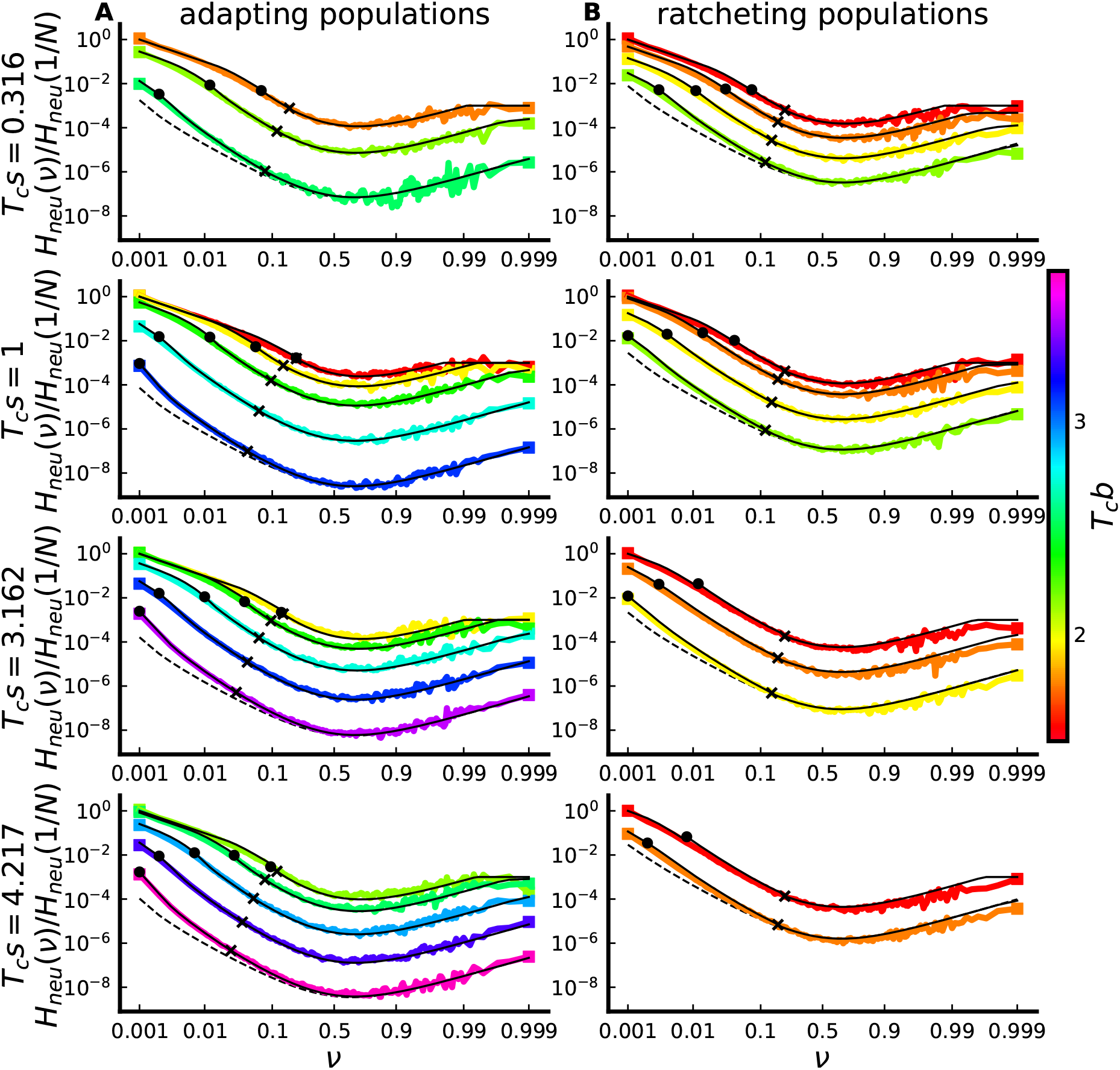
Comparison between predicted and simulated neutral SFSs, for populations subject to an exponential DFE. Solid black lines denote theory predictions; dashed black lines denote predictions of the BSC SFS for one parameter combination in each panel. The left side corresponds to adapting populations, while the right column corresponds to ratcheting populations. The *T_c_s* values denoted on the left-hand side are the values of *T_c_*〈*s_f_*〉 for adapting populations, and the values of *T_c_*〈*s*〉 for ratcheting populations. For computational reasons, SFSs are observed with a sample size of 1000 individuals, in some cases smaller than the population size *N*. The expected SFS depends on sample size (in addition to the frequency *ν*); theory curves are appropriately downsampled versions of the piecewise-defined function given in Eq. (75). Parameter combinations lie on the same grid used for comparison between simulation and theory in the main text.

**FIG. S5.**
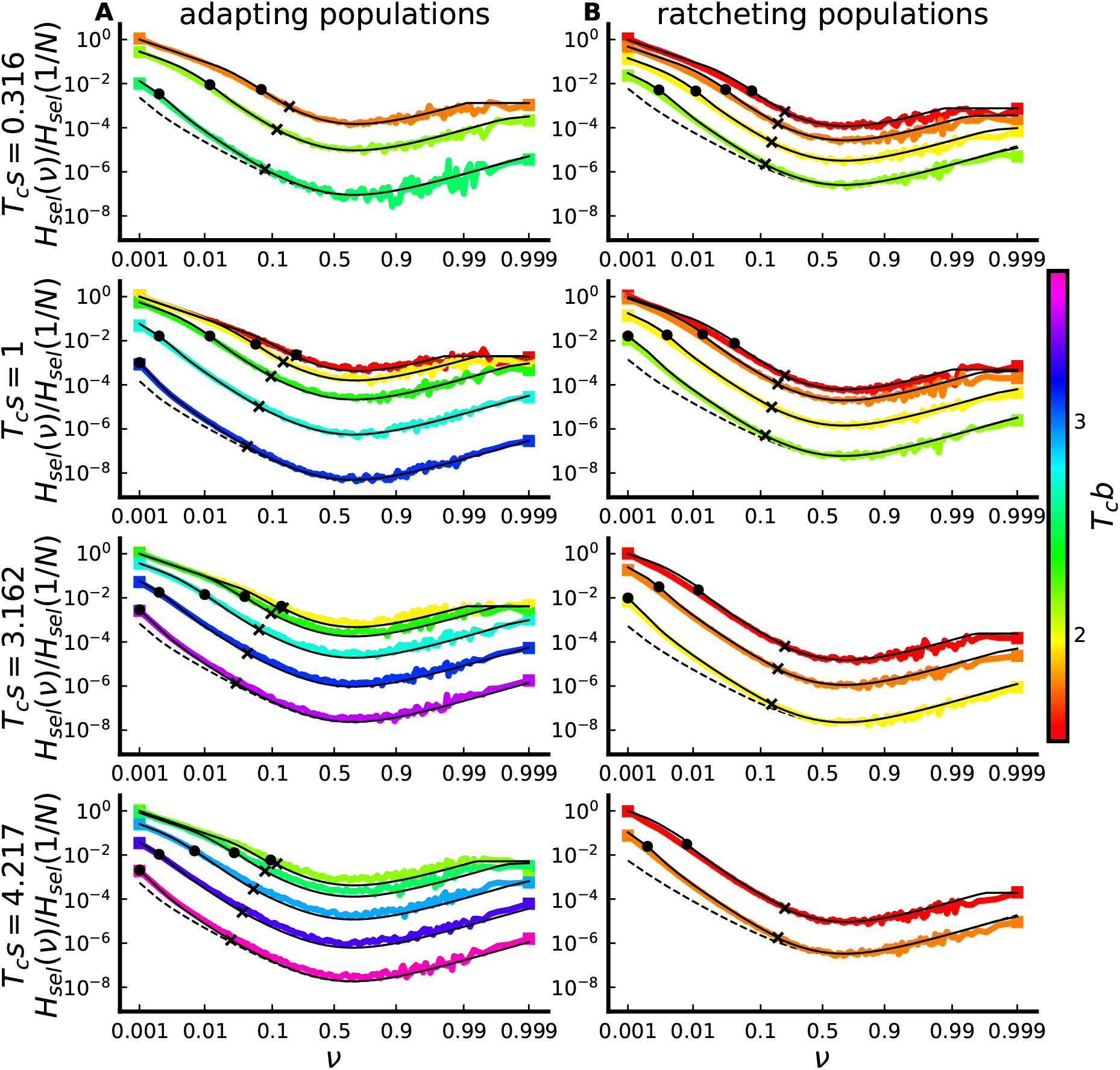
Comparison between predicted and simulated selected SFSs, for populations subject to an exponential DFE. Solid black lines denote theory predictions; dashed black lines denote predictions of the BSC SFS for one parameter combination in each panel. As in Fig. S4, the *T_c_s* values denoted on the left-hand side are the values of *T_c_*〈*s_f_*〉 for adapting populations, and the values of *T_c_*〈*s*〉 for ratcheting populations.

**FIG. S6.**
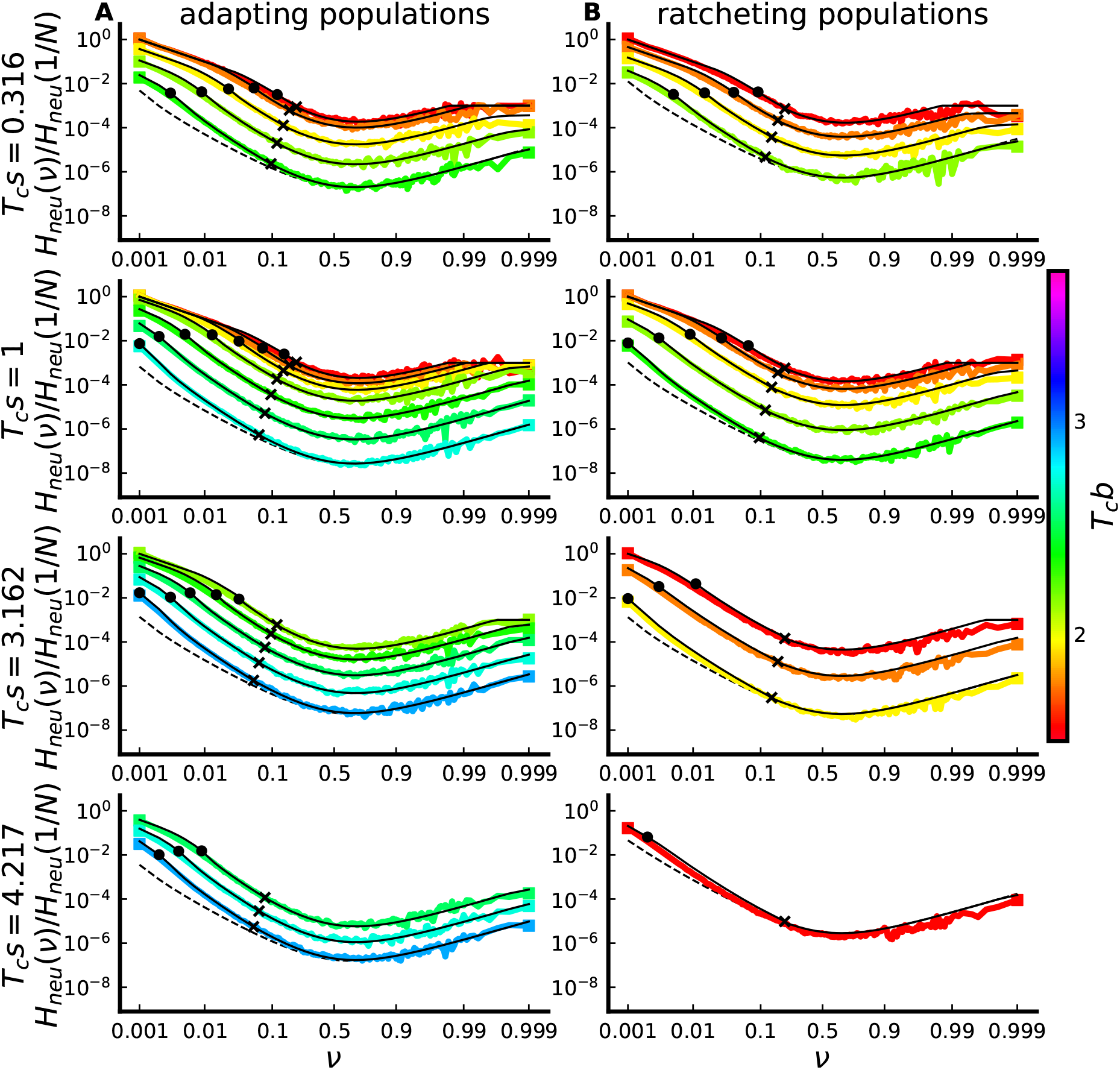
Comparison between predicted and simulated neutral SFSs, for populations subject to an single-effect DFE. Solid black lines denote theory predictions; dashed black lines denote predictions of the BSC SFS for one parameter combination in each panel. As in Fig. S4, the *T_c_s* values denoted on the left-hand side are the values of *T_c_*〈*s_f_*〉 for adapting populations, and the values of *T_c_*〈*s*〉 for ratcheting populations.

**FIG. S7.**
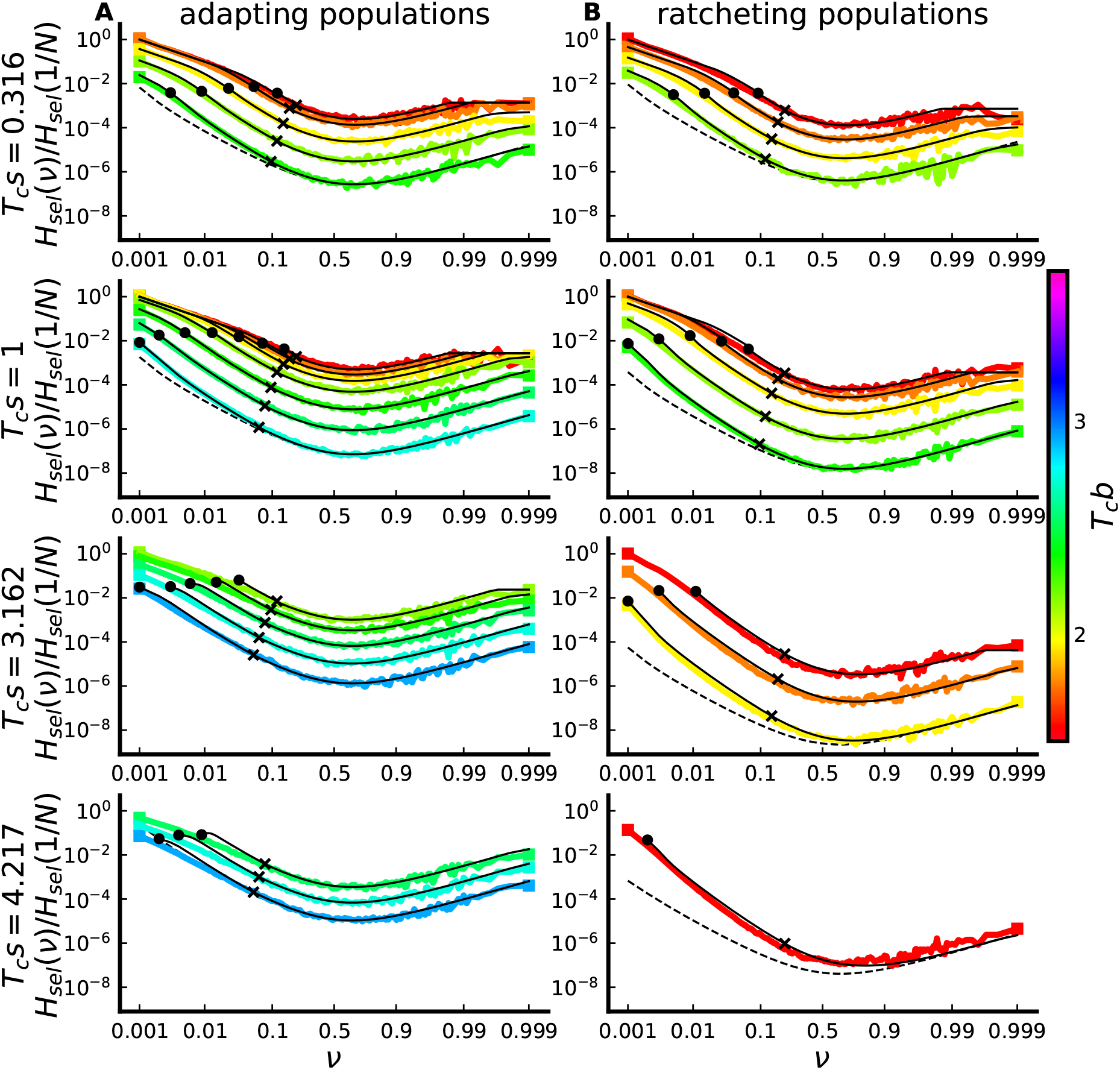
Comparison between predicted and simulated selected SFSs, for populations subject to an single-effect DFE. Solid black lines denote theory predictions; dashed black lines denote predictions of the BSC SFS for one parameter combination in each panel. Theory predictions break down for *ν* < 2/(*Nσ*) when *s* > *b*; for clarity, we have omitted these predictions. As in Fig. S4, the *T_c_s* values denoted on the left-hand side are the values of *T_c_*〈*s_f_*〉 for adapting populations, and the values of *T_c_*〈*s*〉 for ratcheting populations.

**FIG. S8.**
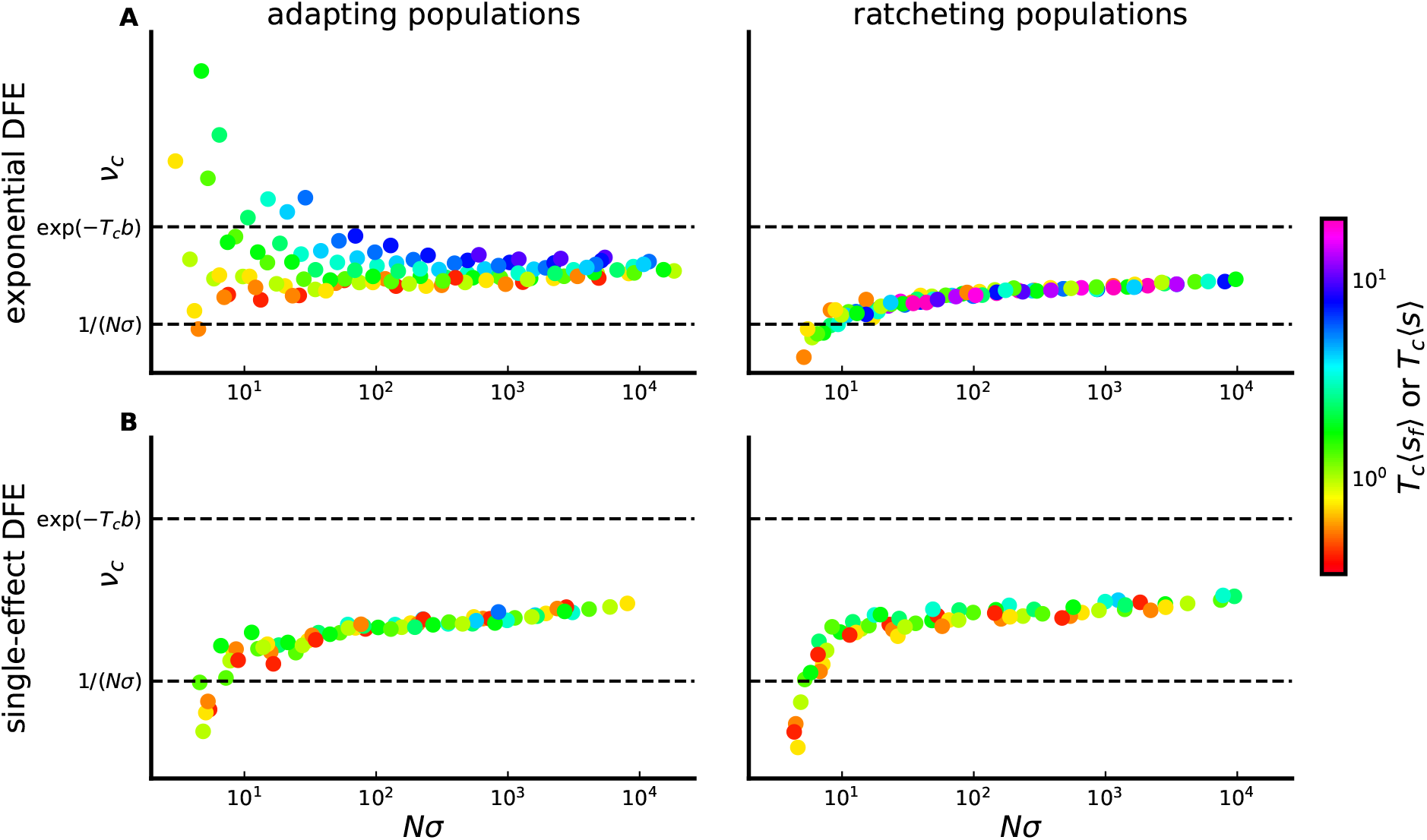
Bounds on the crossover frequency *ν_c_*. The location of a point along the vertical axis denotes the crossover frequency *ν_c_* of that simulated population, extracted from its SFS ratio curve using spline interpolation. The vertical axis is on a logarithmic scale, scaled such that dashed lines correspond to *ν_c_* = 1/(*Nσ*) and *ν_c_* = *e*^−*T_c_b*^. Note that *T_c_b* differs even among simulated populations with the same value of *Nσ*. For large *Nσ*, we can see that 1/(*Nσ*) < *ν_c_* < *e*^*T_c_b*^. Points are colored according to their values of *T_c_* 〈*s_f_*〉 (for adapting populations, in A) or *T_c_*〈*s*〉 (for ratcheting populations, in B). Simulated populations are identical to those considered in Fig. 6.

**FIG. S9.**
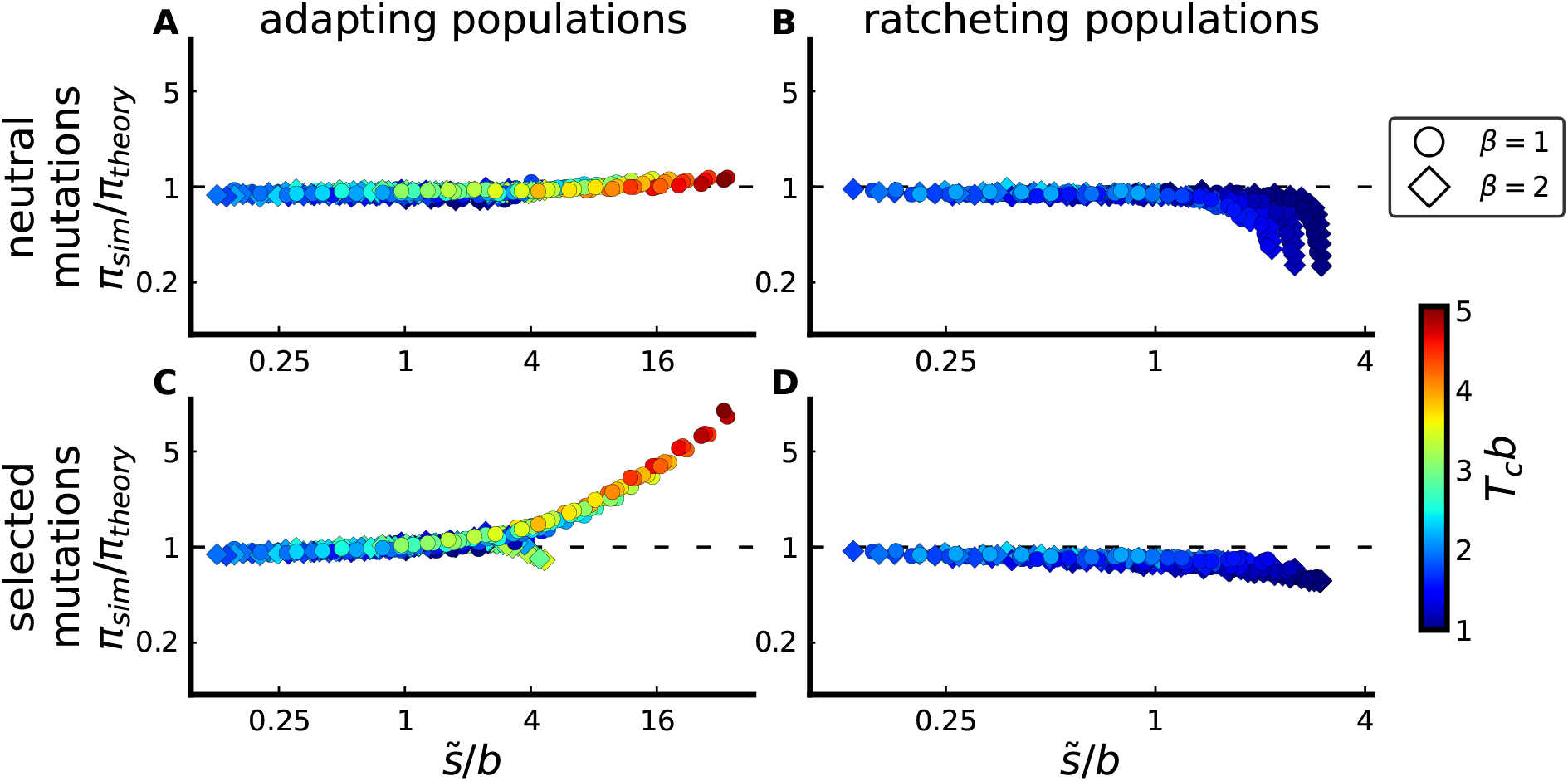
Comparison between simulated and predicted heterozygosity π of neutral mutations (A and B), and between simulated and predicted heterozygosity π of selected mutations (C and D), for adapting populations (A and C) and ratcheting populations (B and D). Parameters are identical to those simulated in Fig. 4. Predictions are obtained by integrating over the piecewise-defined approximation to the SFS given in the main text.

